# Glycan recognition by a plant sentinel immune receptor

**DOI:** 10.1101/2025.09.28.679030

**Authors:** Pedro Jiménez-Sandoval, Caroline Broyart, Owen Kentish, Hyun Kyung Lee, Klara Culjak, Uwe Osswald, Meriem Aitouguinane, Emanuele Tettamanti, Manon Schmidli, Lu Zhang, Charles Roussin-Lévéillée, Diego José Berlanga, Marina Martin-Dacal, Miguel Angel Torres, Varun Kumar, Patricia Fernández-Calvo, José M. Jimenez-Gomez, Alberto P. Macho, Fabian Pfrengle, Lucía Jordá, Antonio Molina, Julia Santiago

**Author notes:** Corresponding author. Current addresses: Patricia Fernández-Calvo, Misión Biológica de Galicia-Consejo Superior de Investigaciones Científicas, Av. de Vigo, s/n. Campus Vida. 15705 Santiago de Compostela, Spain Varun Kumar, Centre for Life Sciences, Mahindra University, Hyderabad, Telangana 500043, India. These authors contributed equally to this work (Caroline Broyart, Owen Kentish, Hyun Kyung Lee).

## Abstract

Pathogens target and degrade the extracellular matrix surrounding plant cells. A central question is how cell wall–derived damage-associated molecular patterns (DAMPs) are recognized and integrated to trigger immune responses. We address this question by determining the structure of the multidomain receptor IGP1 in both apo form and bound to the cellulose-derived DAMP cellotriose. Structural analyses reveal that constitutive Leucine Rich Repeat–malectin interactions preconfigure IGP1 for ligand recognition and that the receptor features a highly specific sugar-binding pocket in the LRR domain capable of distinguishing fine variations in glycan structures. By directly sensing cello-oligomers, IGP1 acts as a cell wall sentinel that links pathogen-induced wall degradation to immune alerting, equipping plants to mount rapid and robust defense responses.

## INTRODUCTION

Plants rely on their cell walls not only as structural scaffolds but also as dynamic frontlines where environmental adaptation and immunity start (*1–4*). Composed largely of cellulose, hemicelluloses, and pectins, cell walls form the first barrier that pathogens must overcome (*5*). To breach this defence, many pathogenic microbes secrete cell wall–degrading enzymes (CWDEs), releasing polysaccharides fragments into the extracellular space (*6–8*). While they can act as nutrients for microbes these wall-derived glycans, serve as damage-associated molecular patterns (DAMPs), signalling danger to the host, and activating its immune defenses (*6*, *9–11*). Cell wall-derived DAMPs include β-1,4-glucan fragments of cellulose, oligogalacturonides (OGs) from homogalacturonan, xyloglucan- and mannan-derived oligosaccharides, as well as arabinoxylan- and xylan-derived fragments, which trigger immune outputs when applied to plants (*12–18*). Despite their importance, however, the molecular mechanisms by which plants perceive cell wall glycans and integrate these signals into immune pathways remain poorly understood. Here, we address this longstanding question by structurally and functionally characterizing the multidomain receptor kinase Impaired in Glycan Perception 1 (IGP1)/Cellulose Oligosaccharide Receptor Kinase 1 (CORK1), which directly binds cellulose-derived oligosaccharides (*17*). These findings establish a new framework for DAMP perception and immune signaling activation in plants.

### Constitutive LRR–malectin interaction preconfigures IGP1 for ligand sensing

The Leucine-Rich Repeat-malectin receptor kinase (LRR-Mal) IGP1/CORK1 acts as a direct receptor of the cello-oligomers cellotriose (CEL3) and cellopentaose (CEL5) (*17*, *18*). The IGP1 ectodomain comprises a canonical plant LRR domain (*19–21*), followed by a malectin domain (*22*, *23*) connected via a flexible loop (Fig. 1A). This unique IGP1 multidomain architecture prompted us to determine its apo structure to reveal its intrinsic configuration. Crystallization of the apo receptor yielded diffracting crystals at 2.16 Å resolution (PDB: 9HHU) (Table S1). The structure revealed a compact and stable multidomain architecture in which the metal-bound malectin domain (*23*) forms an extensive interface with the LRR domain via a network of polar and hydrophobic contacts, with the connecting loop integrating directly into this interface reinforcing structural integrity (Fig. 1, A and B, and fig. S1). Collectively, the apo structure shows that these domain interactions shape the IGP1 ectodomain into a preconfigured state for specific ligand recognition.

**Fig. 1.**
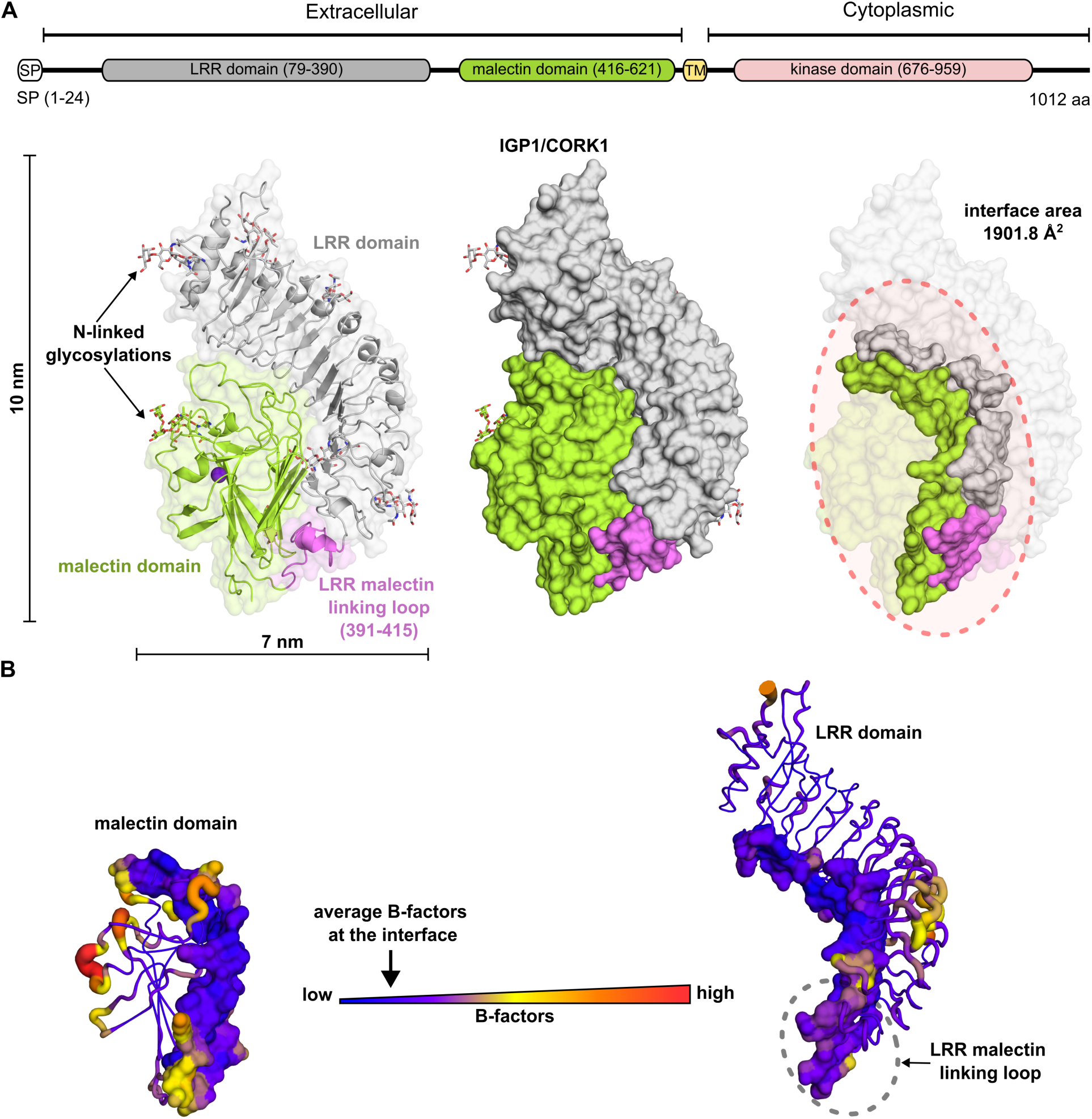
The apo IGP1 structure reveals a constitutive LRR–malectin interaction that preconfigures the receptor for ligand sensing. (**A**) The gene product of IGP1 is illustrated in a domain topology scheme, which includes a signal peptide for secretion (SP), an ectodomain with an LRR domain linked to a malectin domain, a single transmembrane helix (TM), and a cytoplasmic region containing a kinase domain. The crystal structure of the ectodomain of apo IGP1 reveals a large interface between the LRR domain (grey), the linking loop (pink) and the malectin domain (lime green) that stabilizes tertiary structure of the receptor. N-linked glycosylations are represented in stick format, with different colors indicating their locations on the LRR (grey) and malectin domains (lime green). The purple sphere highlights a structural metal atom (likely Ca^2+^). (**B**) Temperature factors (B-factors) analysis of the apo IGP1 ectodomain structure, shows that the interface formed by the malectin domain with both the LRR domain and the linking loop, exhibits relatively low values consistent with a structurally stabilized multidomain configuration.

### The specificity of the IGP1 receptor surface is determined by stereoselective carbohydrate recognition

To uncover the molecular basis of IGP1 cellotriose sensing, we next determined the crystal structure of IGP1 bound to cellotriose (a molecule of three β-(1→4)-linked D-glucose units; CEL3) at 2.62 Å resolution (PDB: 9HHX) (Table S1), allowing direct visualization of carbohydrate recognition and ligand-induced remodelling of the receptor surface. Contrary to expectations that the malectin domain binds cellotriose (*22*), the sugar ligand is instead fully accommodated within the LRR domain, with the malectin domain primarily shaping the ligand-binding pocket (Fig. 2, A and B, and fig. S2). Notably, the two conserved phenylalanine residues (Phe520 and Phe539), previously suggested to contribute to the carbohydrate ligand-binding pocket (*18*), are buried within the hydrophobic core of the malectin domain, where they play a critical role in maintaining the structural integrity and stability of the receptor extracellular domain (fig. S3). Mutation of these residues to alanine would likely destabilize the entire ectodomain fold, providing a structural explanation for the impaired Ca²⁺ signaling observed upon CEL3 treatment in plant complementation assays by Tseng et al., (*18*). Similar to other receptor–ligand complexes (*24*–*27*), the IGP1 ligand-binding surface is free of N-linked glycosylations (Fig. 2A). Remarkably, comparison of the apo and CEL3-bound IGP1 structures revealed minimal rearrangements of the extracellular domain upon ligand binding (global RMSD 0.231 Å). That indicates, as suggested by the apo structure, that the IGP1 ligand-binding pocket is largely preformed, with only small structural adjustments around the pocket refining ligand accommodation (Fig. 2, B and C; and fig. S2). While cellotriose in solution is conformationally flexible due to torsional freedom of glucopyranose rings (Phi, (φ), (Psi), ψ angles) (fig. S4) (*28*); in IGP1 it adopts a fully extended conformation that spans the entire pocket. The reducing-end hemiacetal is deeply accommodated against the malectin-facing wall of the pocket, coordinated by a network of polar interactions limiting its flexibility. This planar geometry is stabilised by a combination of polar contacts and CH–π interactions with electron-rich aromatic residues (Fig. 2, D and E). Specifically, IGP1 Trp146 and Tyr196 engage the terminal glucose rings through face-to-face CH–π contacts, which exploit the partial positive charge of glucose C–H bonds and the π-electron density of the aromatic rings to create highly directional and energetically favourable interactions (*29–32*). The aromatic clamps anchor the first (subsite 1) and third (subsite 3) glucosyl residues, while the central sugar (subsite 2) is further stabilized by direct and water-mediated polar contacts with side-chain residues lining the binding cleft (Fig. 2, D and E, and fig. S4). The coordinated action of the three subsites enforces the planar conformation of the trisaccharide and locks it into a precise orientation within the binding cleft, effectively creating a new receptor surface. This structural arrangement suggested that IGP1 achieves high ligand selectivity by simultaneously requiring full occupancy of the three subsites and the correct stereochemical configuration. To test this, we carried out isothermal titration calorimetry (ITC) assays with purified IGP1 ectodomain and structurally related oligosaccharides. While cellotriose bound with high affinity (Kd ∼1 µM; Fig. 2, H and J; fig. S5) (*17*), cellobiose (lacking one glucose unit) showed no detectable binding. Likewise, maltotriose, which preserves chain length but carries α-(1→4) instead of β-(1→4) linkages of D-glucose, failed to bind (Fig. 2, F and H; fig. S5). To further probe sugar discrimination, we also tested the DAMPs xylotriose and xylotetraose, containing five-carbon backbones instead of six, and again observed no binding (Fig. 2, G and H; fig. S5). Thus, IGP1 operates as a highly selective sensor, integrating chain length and stereochemistry to achieve precise recognition of cellotriose. We next validated this structural model by targeted mutagenesis of ligand-binding pocket residues. In agreement with the crystallographic interpretation, substitutions in residues shaping the outer malectin pocket surface (Cys592Ala, Gln596Ala) had no measurable effect on binding. By contrast, mutations of the aromatic anchors at subsites 1 and 3 (Trp146Ala, Tyr196Ala) completely abolished cellotriose interaction, confirming their essential role in sugar recognition. Moreover, disruption of the hydrogen-bonding network lining across the binding pocket, including the Asp74 in the N-terminal interacting loop, (IGP1^D74A.N126A. D174A^) likewise eliminated detectable binding, highlighting the cooperative contribution of polar contacts to sugar sensing (Fig. 2, I and J; and fig. S6).

**Fig. 2.**
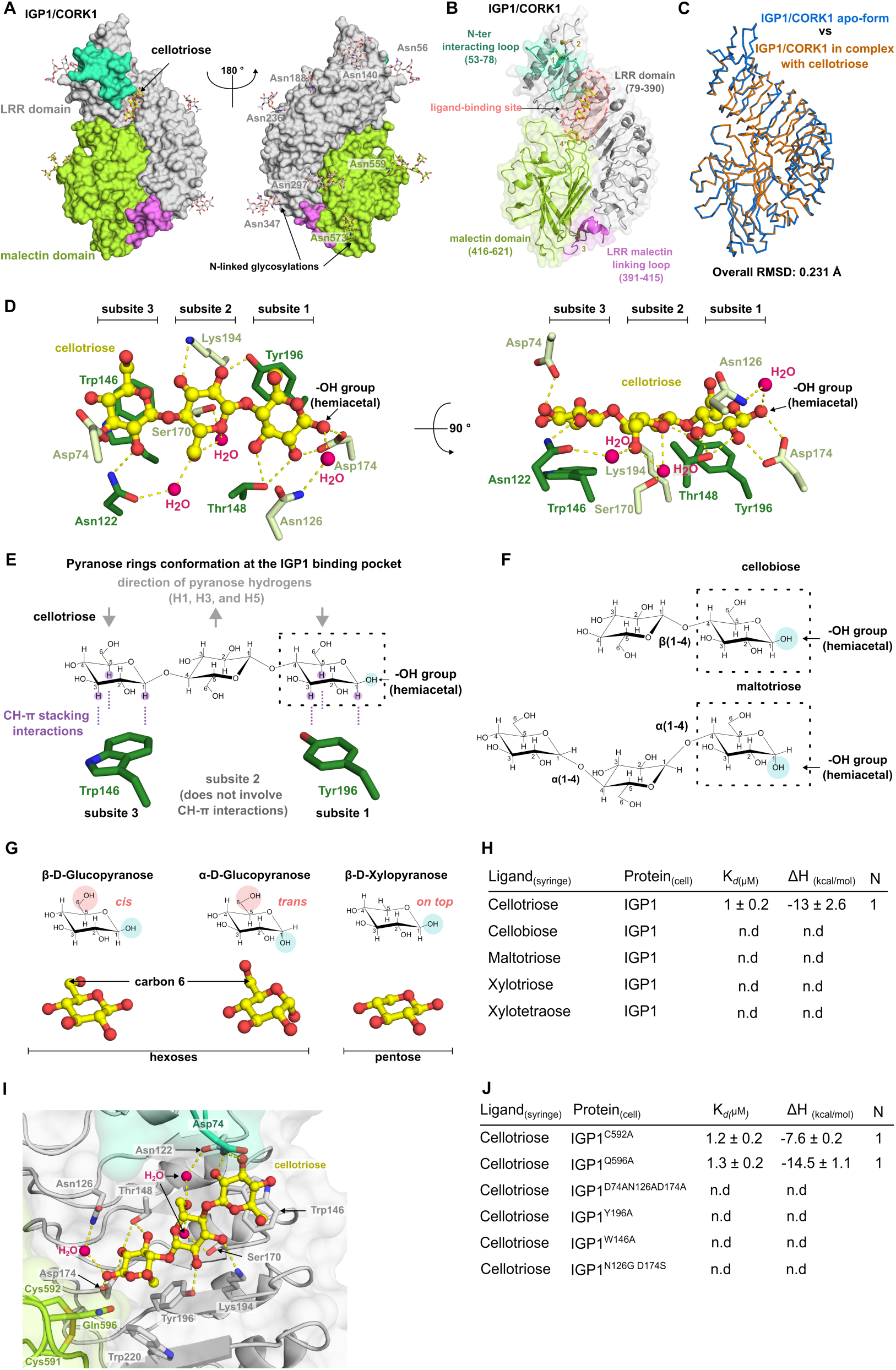
The cellotriose-bound IGP1 structure reveals stereoselective carbohydrate recognition as the basis for receptor surface specificity. (**A**) Surface representation of the IGP1 ectodomain showing cellotriose accommodated within a ligand-binding pocket formed by the LRR scaffold, with the malectin domain shaping one side of the cavity. N-linked glycans (sticks) are positioned away from the binding site. The LRR domain is depicted in gray, the malectin domain in lime green, the linking loop in magenta and the N-terminal interacting loop in cyan. **(B)** Cartoon model highlighting domain organization and structural features of the binding pocket. The LRR– malectin connecting loop is shown in pink, and conserved disulfide bonds are indicated in yellow and labeled. (**C**) Superposition of ligand-free (blue) and cellotriose-bound (orange) structures demonstrates minimal global rearrangement upon binding, consistent with a preconfigured architecture (RMSD shown). (**D, E**) Close-up views of the three subsites that accommodate the glucopyranose rings of cellotriose. CH–π stacking interactions mediated by Tyr196 and Trp146 coordinate rings 1 and 3, respectively, and stabilize the linear orientation of the ligand (yellow sticks). Water-mediated hydrogen bonds stabilize the central sugar (rotated view shown on right). The hemiacetal hydroxyl group faces the malectin domain. Hydrogen bonds (yellow dashed lines) and water molecules (pink spheres) are indicated. (**F**) 2D representation of maltotriose and cellobiose, α- and β-oligomers of glucose (D-glucopyranose), respectively. The hydroxyl group (-OH) of the hemiacetal is highlighted with a blue circle and the reducing end is enclosed within the dashed rectangle. The reducing end is enclosed within the dashed rectangle. (**G**) 3D representation of α- and β-anomers of D-glucose, and β-anomer of D-xylose. Hemiacetal –OH (blue) and terminal–CH₂OH (pink) are highlighted. (**H**) ITC assays show that binding requires full occupancy of all three subsites by β(1→4)-linked glucopyranose rings (DP ≥ 3), conferring high stereoselectivity of IGP1 for cellotriose. (**I**) In addition, cellotriose is tightly coordinated within the IGP1 LRR-domain binding pocket through a dense network of hydrogen bonds. The LRR and malectin domains are shown in grey and lime green; the Asp74 in the N-terminal interacting loop (cyan); cellotriose is in yellow sticks, water molecules in hot pink, and hydrogen bonds as yellow dashed lines. (**J**) Isothermal titration calorimetry (ITC) reveals that a structurally intact binding pocket is required for the specific recognition of cellotriose. Binding thermodynamics are shown for wild-type and mutants of IGP1; n.d., no binding detected. ITC table summarizes Kd (µM), stoichiometry (N=1), and mean ± SD from ≥2 experiments; n.d., no binding detected. Data are presented as mean ± SD from at least two independent experiments

Having validated the structural model through mutagenesis, we then asked whether these glycan recognition features were conserved across the broader LRR–Malectin family. Pocket conservation analysis identified four of thirteen *Arabidopsis* members, including IGP4 (At1g56140) and IGP3 (At1g56130), previously implicated in glycan-triggered immunity (*14*, *17*), sharing key IGP1 architectural features such as the aromatic clamps at subsites 1 and 3. We therefore classified this subset as the IGP1-like subfamily (Fig. 3, A and B; fig. S7). In contrast, while the remaining family members preserve the canonical LRR–Malectin domain architecture and a highly conserved domain interface, their ligand-binding pockets are highly divergent (fig. S7 and S8), suggesting that this receptor family has evolved distinct carbohydrate or other ligand specificities. To probe the basis of receptor specificity within the IGP1-like subfamily, we engineered IGP1 variants in which two polar residues were exchanged with those from IGP4, a close paralog (Fig. 3, A and B; fig. S7). Despite retaining the aromatic clamps, the IGP1^N126G.D174S^ variant lost all binding activity (Fig. 2J), aligning with the lack of interaction previously observed for native IGP4 and cellotriose in ITC assays (*17*). These results demonstrate that both the aromatic anchoring residues and the precise polar network are indispensable for bona fide glycan recognition, and that even limited substitutions are sufficient to shift specificity across IGP1-like receptors. Extending the analysis across angiosperms, we identified IGP1-like orthologs in multiple plant species. Remarkably, the aromatic residues that coordinate the β-pyranose rings at subsites 1 and 3 are highly conserved, suggesting that the molecular mechanism for β-D-glycan recognition has been maintained across flowering plants as a common strategy for perceiving cell wall–derived oligosaccharides (figs. S9 and S10).

**Fig. 3.**
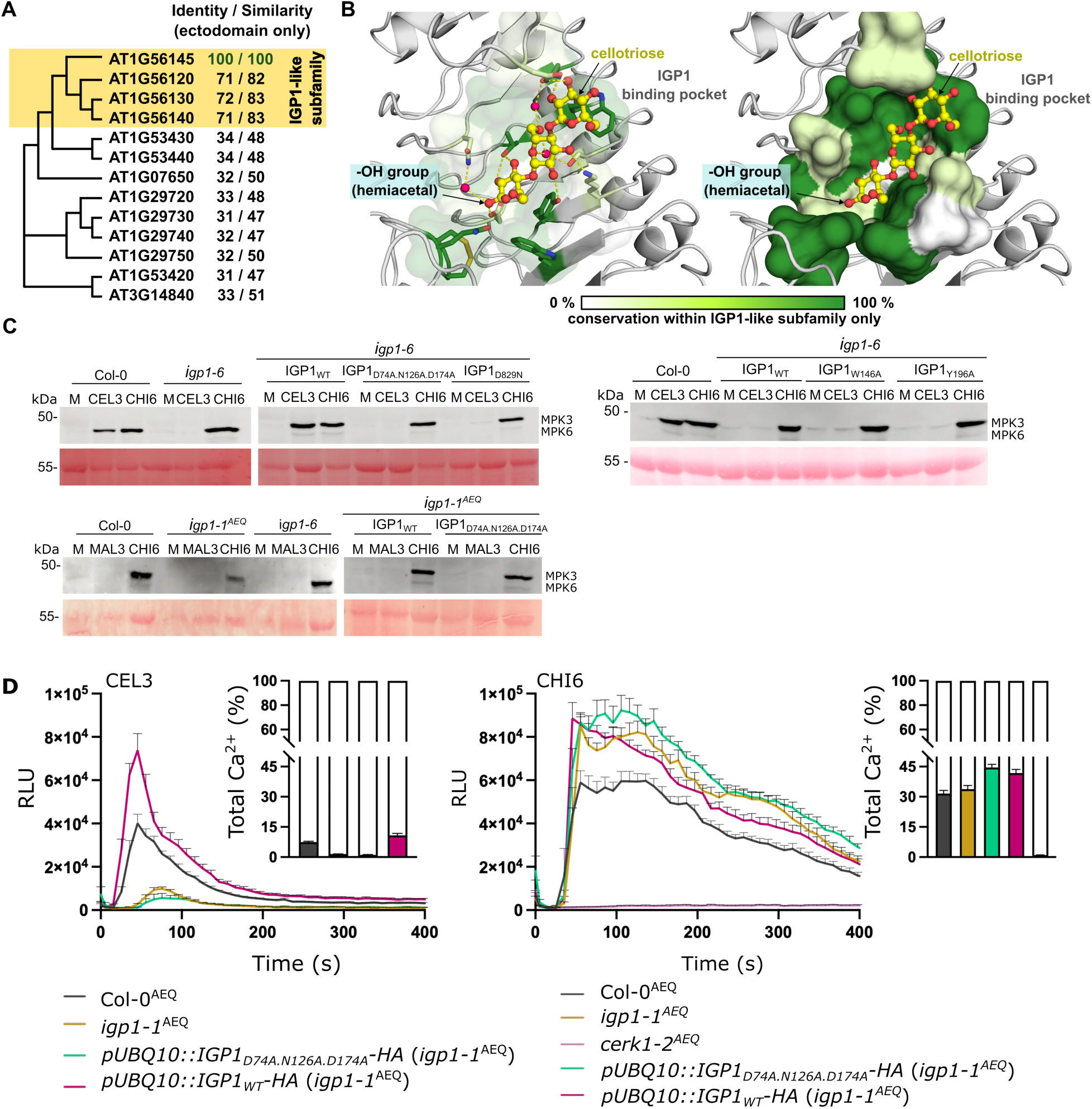
Precise ligand recognition is required for IGP1 signaling activation. (**A-B**) Conservation of ligand-recognition features within the IGP1-like receptor subfamily. Phylogenetic tree of Arabidopsis LRR–malectin receptors with the IGP1-like clade highlighted in yellow. (**B**) Residue conservation mapped onto the IGP1 pocket (cartoon and stick, left; surface, right). The hemiacetal –OH group of cellotriose faces the malectin domain; water molecules (pink) and hydrogen bonds (yellow dashed) are indicated. (**C**) Ligand binding is essential for downstream IGP1 MAPK activation. Cellotriose, but not maltotriose, triggers MAPK phosphorylation in Col-0 seedlings; this response is lost the *igp1-6* mutant and restored by wild-type IGP1 but not by ligand-binding or dead-kinase variants. Ponceau S serves as a loading control. Data shown are from one representative experiment out of three that yielded similar results. (**D**) Cellotriose-induced Ca²⁺ bursts require direct IGP1 ligand recognition. Aequorin-based Ca^2+^ assays show that CEL3 elicits rapid Ca²⁺ influx in Col-0^AEQ^ and in complemented *igp1-1*^AEQ^ lines expressing wild-type IGP1, but not in *igp1-1^AEQ^* mutant or lines expressing a binding-deficient IGP1 variant. CHI6-induced Ca²⁺ bursts, which are CERK1-dependent, remain unaffected in the *igp1-1^AEQ^* mutant and IGP1 complementation lines. *cerk1-2*^AEQ^ mutant, impaired in CHI6 perception, was used as control for CHI6-mediated responses. Total Ca²⁺ discharge was normalized after addition of 1 mM CaCl₂. Data represent mean ± SEM (n=8); results are from one of three independent experiments with similar outcomes.

### Precise ligand recognition drives IGP1 signaling

Cellotriose perception activates rapid immune-related responses in plants, such as cytosolic Ca²⁺ influx and MAPK phosphorylation (*17*, *18*, *33–37*). In our previous work, the *igp1-1^AEQ^* allele, which constitutively expresses the aequorin Ca^2+^ sensor protein (AEQ), was shown to be impaired in the rapid Ca²⁺ burst triggered by cellotriose stimulation compared with Col-0^AEQ^ control plants (*17*). Building on this, we identified in a genetic screen additional independent *igp1^AEQ^* mutant alleles that abolished cellotriose-induced Ca²⁺ responses (fig. S11) (*38*). These findings, consistent with our biochemical and structural analyses, establish IGP1 as a dedicated receptor for cello-oligomer perception in *Arabidopsis thaliana*. To directly link structure to function, we generated transgenic lines overexpressing either wild-type IGP1, a kinase dead version or the binding-pocket variants in *igp1-1^AEQ^*and *igp1-6* backgrounds ensuring comparable expression across lines (fig. S11, S12 and Table S2). All mutant variants expressed and localized to the plasma membrane, indicating that the introduced mutations did not impair receptor secretion (fig. S12, S13 and Table S2). Only lines expressing wild-type IGP1 restored cellotriose-induced Ca²⁺ burst and MAPK activation in *igp1* alleles, whereas the kinase-dead or the binding-pocket mutants failed to complement signaling (Fig. 3, C and D, and fig. S6). Consistent with the stereoselectivity observed in our vitro assays (Fig. 2H), maltotriose treatment did not trigger MAPK activation in an IGP1-dependent manner, underscoring the requirement for β-(1→4) linkage specificity (Fig. 3C). Moreover, cellopentaose elicited Ca²⁺ and MAPK responses comparable to cellotriose in native IGP1 complemented lines, in agreement with our previous biochemical data showing that IGP1 can bind cello-oligomers longer than cellotriose (fig. S6) (*17*). Together, these results demonstrate that IGP1 signaling requires both precise carbohydrate recognition at the extracellular binding pocket and an intact cytoplasmic kinase domain to couple ligand engagement with immune signaling outputs. This establishes a direct mechanistic link between molecular recognition of cell wall β-glucan fragments and the activation of immune-related responses, thereby positioning IGP1 as a key player in cell wall–derived danger perception.

Many plant receptor kinases require heteromerization with shape-complementary co-receptors for activation. The best-studied examples are members of the SOMATIC EMBRYOGENESIS RECEPTOR KINASE (SERK) family, which function as versatile co-receptors across diverse signaling pathways (*24–26*, *39*). BAK1/SERK3, in particular, acts as a co-receptor for several immune receptors (*27*, *40–42*), and has been reported to undergo phosphorylation upon cellotriose treatment, with *bak1* mutants showing partially impaired CEL3 responses (*36*), suggesting a potential direct role for SERK proteins in IGP1 signaling. To address this possibility, we examined whether ligand binding to IGP1 generates a co-receptor docking surface comparable to those observed in other receptor–ligand complexes. Structural analysis revealed that cellotriose binding with IGP1 creates a newly exposed surface of ∼200 Å², well within the range of surfaces (∼100– 200 Å²) that are typically recognized by SERK proteins in receptor complexes (fig. S14). We expressed and purified the ectodomains of several SERK proteins and tested their direct interaction with IGP1–CEL3 using ITC assays. However, no binding was detected, indicating that SERK proteins are unlikely to function as direct co-receptors for IGP1 (fig. S14). This suggests that IGP1 may employ alternative partners or an unconventional activation mechanism, highlighting a potentially distinct mode of receptor activation in cell wall damage surveillance.

### IGP1 alerts plant immunity through cello-oligomer perception

Notably, although cello-oligomers such as cellotriose rapidly trigger early defense-associated signaling events—including cytosolic Ca²⁺ influx and MAPK phosphorylation (Fig. 3, C and D)— they fail or weakly activate other hallmark immune outputs typically induced by canonical MAMP elicitors, such as, callose deposition, high reactive oxygen species (ROS) production, and growth inhibition (Fig. S15) (*16*, *17*, *43*). This selective response pattern suggested that IGP1-mediated cello-oligomer perception may function as a surveillance system, allowing the plant to detect early signs of cell wall damage associated with pathogen invasion. To explore this possibility, we first mapped the spatial distribution of *IGP1/CORK1* expression using a nuclear-localized GFP reporter driven by the *IGP1* endogenous promoter (*pIGP1::3xNLS-mGFP*) (Table S2). Reporter analyses revealed broad expression in both roots and shoots, with signal detected across all roots developmental zones—including the meristem, elongation, and differentiation zones— as well as in multiple cell layers and emerging lateral roots (fig. S16) (*18*). Expression was also evident in cotyledons and mature leaves, indicating that *IGP1* is widely expressed throughout Arabidopsis tissues, supporting the idea of a generalized role in cell wall surveillance (fig. S16).

To determine whether cellotriose perception modulates immune competence, we analyzed different transcriptional MAMP-reporter lines such (e.g., *pFLS2::NLS-3xmVenus*) (*44*). Confocal microscopy revealed that cellotriose perception enhanced the expression of key immunity receptors, including FLS2, CERK1, and EFR, across different tissues (Fig. 4, A and B, and fig. S17). To extend these observations, we interrogated public RNA-seq datasets of Arabidopsis seedlings exposed to cellotriose (fig. S18 and S19) (*14*, *36*). Notably, cellotriose induced a broad transcriptional program characterized by the upregulation of a wide spectrum of immunity receptor kinases, defense-related enzymes (e.g., chitinases and LOX3), and signaling peptides implicated in plant immunity and disease resistance responses (*11*, *41*, *45–48*). Cellotriose also up-regulated the expression of the DAMP peptides PEP1/2 and SCOOPs, both known amplifiers of Pattern-Triggered Immunity (PTI) (*42*, *49–52*) (fig. S18 – S20). Notably, while *IGP1* expression itself was not boosted by cellotriose, expression of *IGP4*, a member of the *IGP1-like* family, was induced in wild-type plants but not in the *igp1-6* knock-out, suggesting coordinated regulation among this receptor subfamily in monitoring cell wall integrity and danger perception (*14*, *17*) (fig. S20). These findings suggest that IGP1 signaling prepares the immune system by reinforcing receptor and signaling networks at the transcriptional level, thereby enhancing the plant’s readiness to respond to subsequent challenges.

**Fig. 4.**
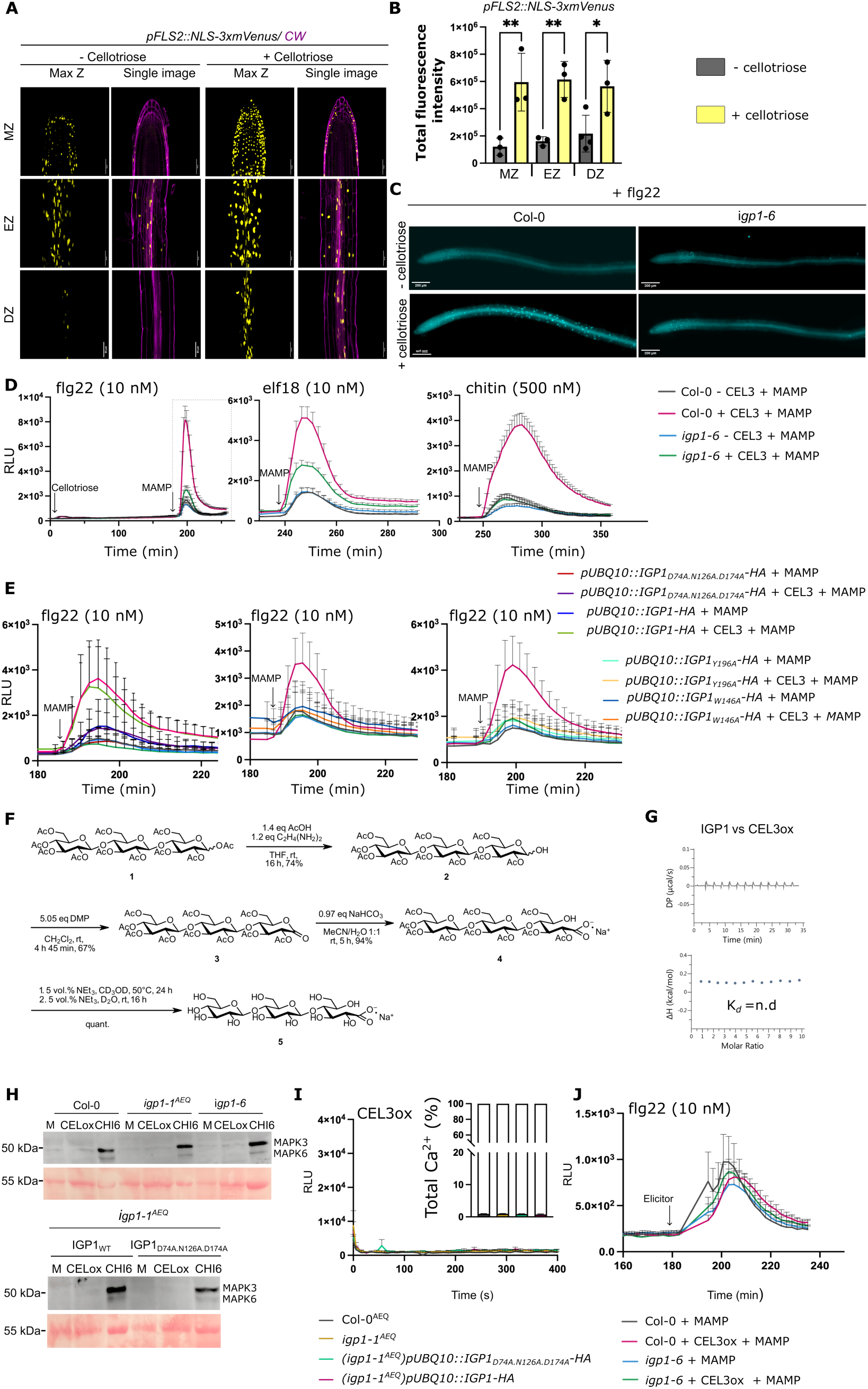
IGP1 alerts the immune system in response to cello-oligomers perception. (**A**) Cellotriose triggers defense gene expression. Confocal images of *pFLS2::NLS-3xmVENUS* seedlings (6 d) show stronger nuclear *mVENUS* signals after 100 μM cellotriose in meristematic (MZ), elongation (EZ), and differentiation (DZ) zones. Maximal z-projections (left) and single planes (right); n = 3. *mVENUS* (yellow) overlaid with calcofluor white (purple). Scale bar, 50 μm. (B) Cellotriose significantly increases nuclear signal intensity. Quantification of FLS2 reporter activation upon cellotriose treatment. Raw intensity density was measured using a maximally projected image. The value represents the total fluorescence intensity of all nuclear signals in the imaged area. Bars show mean ± SD (n = 3–4 roots); *P < 0.05, **P < 0.01 (ANOVA with Tukey’s test). (**C**) Cellotriose enhances callose deposition after MAMP challenge in an IGP1-dependent manner. Representative aniline blue–stained roots 16 h after flg22 treatment; Col-0 vs igp1-6. Scale bar, 200 μm (n ≥ 10). (**D-E**) IGP1-mediated response amplifies MAMP-induced ROS production. Col-0 seedlings pretreated with cellotriose (100 μM, 3–4 h) show stronger ROS production after MAMP challenge, compared to *igp1-6* and IGP1 pocket-binding mutants. n ≥ 10 seedlings, means ± SEM, representative of ≥ 3 experiments. Legends apply to D and E graphs. **(F)** Synthetic scheme for cellotrionic acid (oxidized cellotriose, CEL3ox) chemical synthesis. **(G)** CEL3ox fails to bind IGP1 (ITC), n.d., no binding detected. (**H-J**) CEL3ox fails to activate enhanced IGP1-dependent immune responses. CEL3ox (10 μM) treatment does not induce MAPK phosphorylation (**H**) or cytoplasmic Ca²⁺ burst (**I**) in Col-0, *igp1* alleles, or IGP1 binding-pocket mutants, indicating loss of downstream signaling. Ponceau S serves as loading control. Total Ca²⁺ discharge was normalized after addition of 1 mM CaCl₂. Ca²⁺ burst traces show mean ± SEM. (n = 8). Data shown are representative of three independent experiments. (**J**) Oxidized cellotriose fails to activate an enhanced defense response upon elicitor perception. Pretreatment with CEL3ox (100 µM, 3 h) does not enhance ROS burst after MAMP challenge, in contrast to native cellotriose (**D**). ROS data: n ≥ 10 seedlings, mean ± SEM, representative of ≥ 3 experiments.

We next tested whether this transcriptional reinforcement translates into functional activation of canonical defense outputs. In Col-0 wild-type seedlings, pretreatment with cellotriose markedly enhanced elicitor-induced callose accumulation along root tissues compared to untreated seedlings (fig. S15). This enhancement effect induced by cellotriose, was evident across treatments with different MAMPs, including flg22 and elf18 peptides, and was also detectable using lower concentrations of cellotriose (10 μM). Notably, this response was completely absent in *igp1-6*, demonstrating that IGP1 is indispensable for cello-oligomer–induced activation of callose deposition (Fig. 4C and fig. S21). A similar pattern was observed for reactive oxygen species (ROS) production. Cellotriose alone elicited little to no ROS burst (fig. S15) (*17*); however, seedlings pretreated during 3-4 hours with cellotriose displayed a robust amplification of ROS production upon subsequent challenge with flg22, chitohexaose, or elf18, whereas simultaneous application of cellotriose and the MAMP did not elicit this effect (Fig. 4D and fig.S21). This boosting effect was strictly dependent on IGP1, as it was absent in *igp1-6, igp1-1^AEQ^* and in IGP1 pocket-variant mutant lines defective in cellotriose sensing (Fig. 4D and fig. S21). As observed for callose deposition, ROS enhancement was evident at lower cellotriose pretreatment concentrations (10 μM) and was similarly reproduced with cellopentaose pretreatment (Fig. 4D and fig. S21). Together, these results indicate that IGP1 functions as a sentinel, translating cell wall damage into a system-wide state of immune readiness—positioning cello-oligomer perception through IGP1 as a key evolutionary strategy to alert and amplify defences against pathogens.

### Cellotriose oxidation impairs IGP1-mediated danger perception

Cellotriose is not only released during pathogen-driven cell wall degradation but is also generated naturally during cell wall remodeling processes such as growth and expansion (*53*). To prevent its uncontrolled accumulation in the apoplast and avoid the triggering of IGP1-mediated defences during development or over-immune activation during infection, plants must fine-tune cellotriose homeostasis. A family of cellodextrin oxidases belonging to the FAD-binding berberine bridge enzyme–like (AtBBE-like) superfamily have been proposed to catalyze the oxidation of the anomeric carbon at the reducing end of cello-oligomers (CEL3–CEL6), dampening their immune activity (*54*, *55*). However, the functional consequences of this modification on IGP1 recognition are unknown. To test this directly, we developed a novel chemical synthesis of oxidized cellotriose (cellotrionic acid, CEL3ox) and assayed its immunogenic capacity. Peracetylated cellotriose obtained from cellulose degradation (*56*) was deacetylated at the reducing end and oxidized using a Dess-Martin-protocol (*57*). The resulting lactone was selectively hydrolysed and then deacetylated under mildly basic conditions to give the corresponding sodium salt in high purity after precipitation (Fig. 4F and fig. S22-S25). ITC experiments showed no detectable binding to of CEL3ox to IGP1 (Fig. 4H), consistent with the IGP1–CEL3 crystal structure (Fig. 2, A-B and D; and Fig. S2), which predicts that oxidation at the reducing end disrupts ligand coordination within the receptor binding pocket. Moreover, CEL3ox failed to induce MAPK activation, cytosolic Ca²⁺ influx, or ROS amplification in Col-0 or native IGP1-complemented *igp1* lines (Fig. 4, H–J) compared to CEL3 and CEL5 (Fig. 3, C-D; and fig. S6 and S21). These findings demonstrate that the integrity of glucopyranose ring is essential for IGP1 recognition and reveal that enzymatic oxidation of cello-oligomers provides a biochemical mechanism to neutralize their activity as DAMPs. This couples cell wall metabolism to precise receptor-mediated immune surveillance that prevents unwarranted activation (*58*) while preserving robust responsiveness under large pathogen-driven cell wall degradation.

### IGP1 mediates enhanced plant immunity against pathogens

To close the loop from perception to protection, we next asked whether cello-oligomers (CEL3– CEL5) are naturally generated by pathogens and during infection, and whether their detection by IGP1 confers enhanced disease resistance. To address this, we first assessed cellulose-degrading activity by plant pathogens. Congo red and cellulose–azure assays revealed strong cellulase activity in exudates from *Xanthomonas campestris*, but not in a type II secretion system (T2SS) mutant (*xpsxcs*) (*59–62*) or in *Pseudomonas syringae* pv. tomato DC3000 (Fig. 5, A–B). Consistent with these results, thin-layer chromatography of azu-cellulose digested by *Xanthomonas* exudates detected the specific IGP1 ligands such as CEL3 (Fig. 5C). Similarly, high-performance anion-exchange chromatography with pulsed amperometric detection, HPAEC–PAD analysis of oligosaccharides released upon cellulose incubation with mycelium of the necrotrophic fungus *Plectosphaerella cucumerina* BMM (*Pc*BMM) confirmed time-course accumulation of cellotriose, together with other carbohydrates (Fig. 5D) (*63*). We next assesed whether these *Pc*BMM-derived cello-oligomers are biologically active and perceived by IGP1. Ca²⁺ assays showed that time-course collected fractions from *Pc*BMM-digested cellulose induced a robust cytosolic Ca²⁺ burst in Col-0 wild-type seedlings, but not in *igp1-3^AEQ^* mutant, mirroring responses elicited by pure CEL3 or cello-oligomers derived from cellulose digested with commercial cellulase (Fig. 5, E–G, and fig. S11 and S26). To determine whether this perception translates into functional resistance, we performed Arabidopsis disease resistance assays in Col-0 and *igp1-6*, using *Pseudomonas syringae*, a pathogen with weak cellulase activity (Fig. 5A) and thus little capacity to generate cello-oligomers during infection. Pretreatment of Col-0 plants with CEL3 significantly reduced proliferation of *P. syringae* at 72 h post-inoculation, whereas oxidized CEL3ox or buffer controls provided no protection (Fig. 5H). This PTI-triggering effect was abolished in *igp1-6* plants, confirming that CEL3-mediated resistance is IGP1 dependent (fig. S27). Furthermore, *igp1-6* mutants displayed enhanced susceptibility to the root-colonising bacterium *Ralstonia solanacearum* compared to Col-0 wild-type plants (*64*). Symptom progression and survival reduction in *igp1-6* were similar to those observed in the hypersusceptible mutant *bak1-5/bkk1-1* (Fig. 5, I–J, and fig. S27) (*65*, *66*). Together, these data demonstrate that by detecting cello-oligomers as damage cues, IGP1 acts as a sentinel receptor, alerting the plant to mount a rapid and robust defense against invading pathogens.

**Fig. 5.**
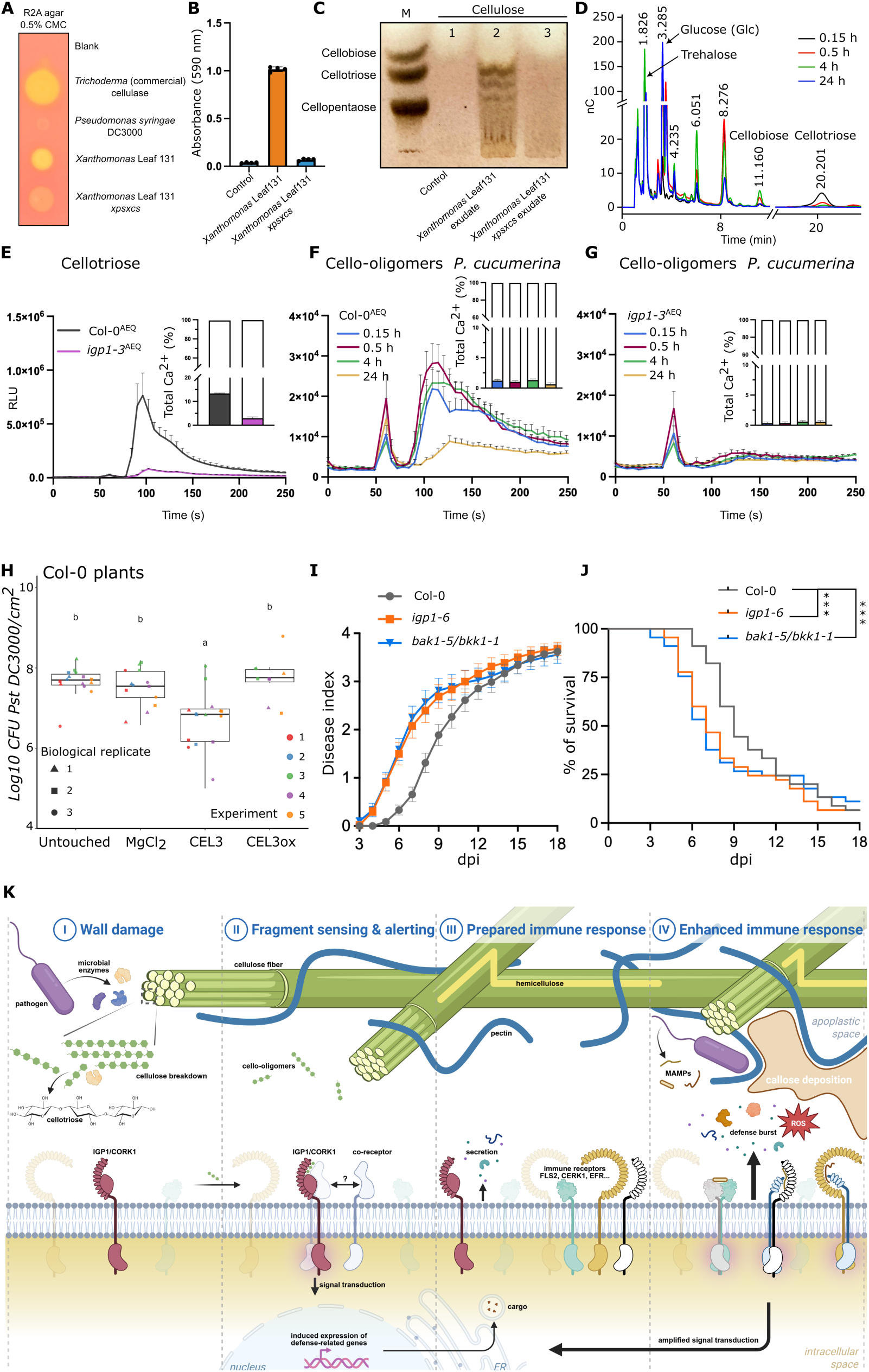
IGP1 senses pathogen-released cello-oligomers to enhance plant immunity and disease resistance. (**A-D**) Plant pathogens with cellulase activity generate cell-wall derived IGP1 ligands. Cell-wall–degrading pathogens such as *Xanthomonas campestris* and *Plectosphaerella cucumerina* (*Pc*BMM) release cello-oligomers during infection. (**A**) Congo red assays show strong cellulase activity for commercial cellulase, *Xanthomonas* Leaf131, but not for the *Xanthomonas* type II secretion (T2S) mutant (*xpsxcs*) or *Pseudomonas syringae* DC3000. (**B**) Quantification of cellulase activity using cellulose–azure dye shows that wild-type *Xanthomonas* exudates degrade cellulose, while the T2S mutant fails to release detectable enzyme activity (absorbance at 590 nm). (C) Thin-layer chromatography of *Xanthomonas* exudate-digested cellulose shows production of cellotriose and cello-oligomers specifically recognized by IGP1. A reference ladder of cellobiose, cellotriose, and cellopentaose was run alongside. (**D**) HPAEC–PAD analysis of time-course (0.15-24 h) degradation of cellulose digested by *Pc*BMM shows accumulation of cellotriose (DP3), as well as other carbohydrates. Identified glycans in HPAEC–PAD are indicated. Data are representative of one of three independent experiments. (**E-G**) IGP1 mediates calcium signaling triggered by cello-oligomers released by *Pc*BMM degradation of cellulose at different time points (0.15-24 h). (**E**) Cellotriose (10 µM) induces a robust Ca²⁺ burst in Col-0^AEQ^ but not in *igp1-3^AEQ^* seedlings. (**F-G**) Fractions from *Pc*BMM-digested cellulose (0.15-24 h) elicit Ca²⁺ influx in wild-type plants in an IGP1 dependent manner. Ca²⁺ flux is shown as relative luminescence units (RLU) over time; values are mean ± SEM (n = 8). (**H**) Cellotriose prepares disease resistance responses in plants. Pre-treatment of Col-0 wild-type plants with CEL3 reduces growth of *Pst* DC3000 after 72 dpi. Untouched plants, MgCl_2_ solution and CEL3ox were used as controls. Statistical significance was analysed by Kruskal–Wallis test followed by Dunn’s post hoc test with Bonferroni correction (p < 0.001). Replicates were performed 3 to 5 times independently. (**I-J**) i*gp1-6* mutant shows enhanced susceptibility to *Ralstonia solanacearum* GMI1000. Col-0, *igp1-6*, and the hypersusceptible *bak1-5/bkk1* mutant were soil-drenched with bacterium suspension. Disease index (0–4 scale; 0 = healthy plant, 4 = dead plant) were scored daily for 18 days (mean ± SEM, n = 45, 3 independent assays) and used to calculate symptom progression (**I**) and survival rates (**J**). Survival curves were analyzed by log-rank and Gehan–Breslow–Wilcoxon tests; **-p< 0.01; ***-p< 0.001;(p-value for *igp1-6*=0.0023; p-value for *bak1-5/bkk1-1*=0.0009). (**K**) Model for IGP1/CORK1-mediated immune activation. I,II Pathogen-secreted cell wall–degrading enzymes release cellulose fragments that are perceived by the plasma membrane receptor kinase IGP1/CORK1. III, This perception triggers an alerting state characterized by transcriptional reprogramming and accumulation of immune receptors, phytokines, and defense genes. IV, Subsequent recognition of elicitors (e.g., flg22, chitin) and phytokines in IGP1-CEL3 boosted cells results in an accelerated and amplified PTI immune response, including amplified ROS burst, callose deposition, and defense gene expression which leads to disease resistance.

## Discussion

The ability of plants to sense structural damage and translate it into immunity is crucial for survival against microbial attack. Our structural and biochemical analysis of the IGP1/CORK1 receptor reveals a highly selective perception mechanism in which glycan chain length and stereochemistry, are integrated to enable precise recognition of cellotriose. Aromatic clamps within the pre-formed binding pocket lock β-linked glucans into place, creating a stringent filter that distinguishes genuine cell wall-generated cello-oligomers from structurally similar glycans, including other cell wall-derived DAMPs (*14*, *17*). It is conceivable that this selectivity is further reinforced by tissue-specific metabolism that might control cello-oligomers homeostasis. In expanding tissues, where cell wall remodeling is particularly active, cellodextrin oxidases are highly expressed and activated (*58*). Oxidation of the reducing end of short oligomers such as cellotriose could therefore occur in situ (*54*, *55*), effectively neutralizing these remodeling byproducts before they accumulate to immunogenic levels. Such a mechanism would prevent inappropriate immune activation during normal growth while maintaining the system’s responsiveness to the higher and more persistent levels of cello-oligomers that accompany pathogen-triggered wall degradation. Moreover, the conservation of these sugars recognition features across flowering plants highlights a widely shared evolutionary strategy for monitoring cell wall–derived oligosaccharides as universal DAMP indicators of colonization of plants by pathogens. Interestingly, similar structural features are observed in certain carbohydrate recognition domains of mammalian immune receptors, where CH–π interactions and hydrogen-bonding networks enforce strict glycan selectivity (*67*, *68*). This highlights the convergent use of conserved biochemical motifs to detect structurally defined carbohydrate danger cues across kingdoms.

Functionally, our results establish IGP1 as a sentinel receptor that converts cello-oligomer perception into enhanced immune activation (Fig. 5K). IGP1 induces a preparatory state characterized by transcriptional reinforcement and receptors accumulation at the membrane, all without activating costly responses (*69*, *70*). This reinforced state enables plants to mount faster and stronger immunity upon subsequent detection of canonical MAMPs, such as flg22 or chitohexaose, producing amplified ROS bursts, enhanced callose deposition, and boosted defense gene expression. By recognising cellulase-released cell wall fragments as actionable danger signals, IGP1 directly couples pathogen-mediated wall degradation to enhanced disease resistance. These findings strengthen the role of β-1,4-D-glucan oligomers as DAMPs, establishing a clear mechanistic link between cell wall damage and immune readiness. Notably, this newly defined immunity layer may also explain how wall damage amplifies local defenses by triggering the targeted deployment of defense responses (*44*). The structural characterization of IGP1–CEL3 recognition advances our understanding of how carbohydrate-derived MAMPs and DAMPs are perceived by plant immune receptors, expanding the diversity of glycan-binding ectodomains to include LRR–malectin receptor proteins.

The structural mechanism described here, highlights cello-oligomers as central danger signals that can contribute to sustainable crop protection and points to IGP1 as a promising tool for crop improvement, offering the potential to enhance disease resistance without growth penalties. Our findings thus position IGP1 as part of a broader family of surveillance receptors that orchestrate immune enhancement through cell wall–derived danger signals, providing plants with a decisive evolutionary strategy: turning structural vulnerability into an alerting defence mechanism.

## Acknowledgments

The authors thank V. Olieric for providing beam time and the staff of beamline PXIII at the Swiss Light Source (SLS), Villigen, for their technical assistance during data collection. We are grateful to Niko Geldner (University of Lausanne) for providing the transcriptional reporter lines. We also thank Philippe Reymond and Edward Farmer for their valuable input on the manuscript. Our thanks extend to Dr. Antje Potthast (Institute of Chemistry of Renewable Resources, BOKU University) for supplying peracetylated oligocelluloses with low DP as starting material for the synthesis of cellotrionic acid, and to Gemma López (CBGP) for her technical support.

## Author contributions

Conceptualization: JS, PJS, AM, LJ, AM, FP

Methodology and Investigation: PJS carried out structural analyses and mutant design. PJS performed protein purification and crystallization, X-ray diffraction and data collection, structure determination and conservation analysis, analysis of public RNA-seq datasets, GO analysis, figure and schematic generation, protein mutant design, data analysis and interpretation. JS and PJS performed crystallization experiments with assistance from ET. PJS, CB, and OK produced and purified proteins and conducted binding assays. CB, and OK performed MAPK assays and western blotting, CB, OK, and ET generated transgenic plant lines. HKL, OK, and MS conducted ROS and callose assays and microscopy. ET, HKL, and OK performed confocal microscopy of reporter lines and assessed membrane localization in *N. benthamiana*. OK, and CB validated mutant lines and differentially expressed genes by qRT-PCR. OK, and CRL carried out *Pst DC3000* infection assays. HKL extracted bacterial exudates and evaluated cellulase activity. KC contributed to mapping and characterization of *igp1* alleles and performed the Ca^2+^ assays. MA performed the cellulose degradation assays with *Pc* BMM mycelium and with commercial cellulase, determined oligosaccharides profiles by HPAEC-PAS in these experiments, and determined Ca^+2^ burst responses in Col-0^AEQ^ vs *igp1-3*^AEQ^ by oligosaccharides derived from cellulose degradation. DJB contributed to the mapping of *igp1* alleles and contributed to the setting up of Ralstonia resistance assays. MMD contributed to the isolation and mapping of *igp1*alleles. MAT contributed to the isolation and mapping of *igp1* alleles and the setting up of Ralstonia experiments. VK contributed to mapping and genetic characterization of *igp1* alleles. PFC contributed to mapping and genetic characterization of *igp1* alleles. FP and UO designed and executed the new chemical synthesis of CEL3ox. APM and LZ designed and perfomed the Ralstonia disease resistance assays shown in the study.

Funding acquisition: JS, AM, LJ, APM, FP Project administration: JS, AM, LJ, APM, FP Supervision: JS, AM, LJ, AM, FP, APM, MAT

Writing – original draft: JS

Writing – review & editing: AM, LJ, APM, PJS, HKL, OK, CB, ET, UO, FP, MAT, CRL

## Data and materials availability

All data are available in the main text or the supplementary materials. RNAseq data of CEL3 are deposited in NCBI under the BioProject number PRJNA1073490. The structures are deposited at the PDB. Materials and raw data files are available upon request to the corresponding author or to Antonio Molina.

