## Supplementary information for "Glycan recognition by a plant sentinel immune receptor"

Current addresses:

**The PDF file includes:**

Materials and Methods
Figs. S1 to S27
Tables S1 to S2
References (70–103)

### Materials and Methods

#### Protein expression and purification

Codon-optimized synthetic genes corresponding to the ectodomain of IGP1 (AT1G56145, residues 25–630) and mutants, were ordered from Invitrogen GeneArt and cloned into a modified pFastBac donor vector (Geneva Biotech) harboring the 30K *Bombyx mori* (*71*) secretion signal peptide, and with a TEV (tobacco etch virus protease) cleavable C-terminal StrepII-9xHis tag. SERK1 (AT1G71830, residues 24–213) and BAK1 (AT4G33430, residues 1-220) were cloned in the same vector harboring azurocidin and native signal peptide, respectively. Baculovirus vectors were generated in DH10 MultiBac *E. coli* cells (Geneva Biotech). Virus amplification was carried out in *Spodoptera frugiperda* Sf9 cells (Geneart, Thermo Fisher Scientific) and was used to infect *Trichoplusia ni* Tnao38 cells (*72*) for protein expression. The cells were grown for 1 day at 28°C and for 2 days at 21°C with gentle shaking at 120 rpm. The secreted proteins were subjected to tandem affinity purification, using $Ni^{2+}$ (HisTrap excel, equilibrated in 25 mM KPi, 500 mM NaCl, pH 7.8; Cytiva) and Strep columns (Strep-Tactin Superflow high-capacity; IBA) equilibrated in 25 mM Tris, 250 mM NaCl, 1 mM EDTA, pH 8.0. Affinity tags were removed using His-tagged TEV protease in a 1:50 ratio at 4°C overnight. Separation of cleaved tags and aggregated proteins was performed using size-exclusion chromatography on a Superdex 200 Increase 10/300 GL column (Cytiva) equilibrated in 20 mM sodium citrate, 150 mM NaCl, pH 5.0. Proteins were further analyzed for purity and structural integrity by SDS-PAGE.

#### Analytical size-exclusion (SEC) chromatography

Analytical size-exclusion experiments were performed using a Superdex 200 Increase 10/300 GL column (Cytiva) equilibrated in 20 mM sodium citrate, 150 mM NaCl, pH 5. 10 to 20 µM of proteins was injected using a 500 µl loop, and the sample was eluted with a flow of 0,5 mL $min^{-1}$. UV absorbance at 280 nm was used to monitor the elution of the proteins. The peak fractions were analyzed by SDS-PAGE followed by QuickBlue Protein Stain (LubioScience LU001000).

#### Crystallization and data collection

Purified apo IGP1 ectodomain was concentrated to 9.1 mg/mL and used for the crystallization. For the IGP1in complex with cellotriose, the concentrated protein was mixed with cellotriose at a 1:2 protein: ligand ratio (≈ 117 µM IGP1: 234 µM cellotriose). Diffracting crystals were obtained for the apo form and IGP1 in complex with cellotriose using the sitting drop method (vapor diffusion) using a 1:1 drop ratio on a condition composed of 0.1 M sodium acetate trihydrate pH 4.6, 0.2 M ammonium sulfate, and 30 % w/v PEG 2000 growing at 18°C. Additionally, 20 % v/v ethylene glycol was used as a cryoprotectant before snap freezing in liquid nitrogen. Single crystals were shot at 100 K and data collection and processing was carried out using the DA+ data acquisition and automatic data processing software that uses XDS (***73*, *74***) at the beamline X06SA at the Synchroton Light Source (SLS) of the Paul Scherrer Institute, Villigen, Switzerland.

#### Structure determination and refinement

Diffraction data was scaled with AIMLESS (*75*). The structure of apo IGP1 and IGP1 in complex with cellotriose was determined by molecular replacement using PHASER (*76*) using the CCP4 software suite (*77*). Initial searching model was generated using the AlphaFold model of AtIGP1 (https://alphafold.ebi.ac.uk/ entry: C0LGH4, residues 25-620). Crystallographic refinement was performed with REFMAC5 (*78*) using the following TLS parameters (4 groups, residues 28-285, 286-467, 468-495, 496-622) determined by the TLS motion determination server (*79*). Manual building was performed in Coot (*80*) based on electron density map interpretation. Solvent molecules were added when supported by electron density and local chemistry. Geometrical restrains for cellotriose were generated using eLBOW (*81*) and used in the refinement of IGP1 in complex. The quality of the models were evaluated using the validation tools in Coot, the wwPDB validation server (https://validate-rcsb-1.wwpdb.org/) and Molprobity (*82*). Supplementary Table S1 shows the data collection and refinement statistics of both apo IGP1 and IGP1 in complex with cellotriose structures.

#### Conservation and structural analysis

The interface between the LRR and malectin domains of IGP1 was analyzed using the PISA server (https://www.ebi.ac.uk/pdbe/pisa/), with interface residues colored based on their relative B-factor values. Multiple sequence alignment of IGP1 was conducted using Clustal Omega (https://www.ebi.ac.uk/jdispatcher/msa/clustalo) (*83*), while the conservation analysis was performed using the ESPript 3.0 server (https://espript.ibcp.fr/ESPript/ESPript/) (*84*). The structural analysis of IGP1 and generation of figure panels were performed using PyMOL (The PyMOL Molecular Graphics System, Version 2.6 Schrödinger, LLC.).

#### Generation of sugar diagrams

The diagrams of the different oligosaccharides used in this study were generated using ACD/ChemSketch (ChemSketch, version 2024.1.1, Advanced Chemistry Development, Inc. (ACD/Labs), Toronto, ON, Canada, www.acdlabs.com.

#### Isothermal titration calorimetry (ITC)

Experiments were performed at 25°C using a MicroCal PEAQ-ITC (Malvern Instruments) equipped with a 200 µL standard cell and a 40 µL titration syringe. Proteins were gel-filtrated into ITC buffer (20 mM sodium citrate, 150 mM NaCl, pH 5.0). Sequencial injections of 3 µL of each ligand (CEL2, CEL3, MAL3, XYL3, XYL4 and CELox) were injected at concentration ranging from 150 to 500 µM into the ITC cell containing the extracellular domain of IGP1 ECD at 9 µM. To assess the interaction between IGP1-CEL3 complex and receptor kinases SERK1 and SERK3/BAK1, an initial ITC experiment was performed between CEL3 and IGP1 (10 µM) as described above. This was followed by a second ITC experiment in which 100 µM of SERK proteins were titrated into a solution containing IGP1 (9 µM) pre-saturated with cellotriose. A total of 13 injections were performed at 150-s intervals with a stirring speed of 500-rpm. Dilution heat was corrected using the thermograph of the titration of the ligand into the cell containing only buffer as a control. Experiments were performed in duplicate, and data were analyzed using the Microcal PEAQ-ITC analysis software. The N values were fixed at 1 during the data fitting and analysis.

#### Carbohydrates used in the experiments

For ITC and MAPK experiments, the following sugars were used: hexaacetyl-chitohexaose (O-CHI6; Neogen), cellobiose (22150; Sigma), cellotriose (O-CTR ; Neogen), cellopentaose (O-CPE; Neogen), maltotriose (J66491;ThermoFisher Scientific), xylotriose (O-XTR; Neogen), xylotetraose (O-XTE; Neogen), cellotriose oxidized (CELox) ( synthetized for this study).

**Selection and genotyping of *igp1* mutant alleles**

Plants used for cytoplasmatic $Ca^{2+}$ measurements were grown in 96-well plates (1 seedling per well) under long-day conditions (14 hours of light) at 19ºC-22ºC in liquid 1/2 MS medium (Martin-Dacal et al.., 2023) (*17*). Ethyl Methanesulfonate (EMS) mutagenized Col-0$^{AEQ}$ seeds (Martin-Dacal et al., 2023) were screened to detect additional *igp1* alleles, as previously described for the *igp1-1*$^{AEQ}$ identification (Martin-Dacal et al., 2023): seedlings were grown *in vitro* for 8 days, and cytoplasmic $Ca^{2+}$ influxes were assessed using a Varioskan Lux Reader luminometer (Thermo Scientific) upon treatment with 100 µM MLG43. Seedlings exhibiting a low response to MLG43 were transferred to soil, self-crossed and subsequently tested for $Ca^{2+}$ burst in F1 seedlings to confirm the impaired response to MLG43 (100 µM) and CEL3 (10 µM). *igp* mutant previously selected (*igp6*$^{AEQ}$: Martin-Dacal et al., 2023) was sequenced and correspond to *igp1-2*$^{AEQ}$ allele in this study, whereas *igp1-3*$^{AEQ}$, *igp1-4*$^{AEQ}$ and *igp1-5*$^{AEQ}$ alleles are described here for the first time (Figure S1) Total $Ca^{2+}$ discharge was performed by treating seedlings with 1M $CaCl_2$ and the $Ca^{2+}$ burst was quantified using the luminometer. *igp*$^{AEQ}$ mutant alleles were backcrossed with Col-0$^{AEQ}$ for genotyping. Leaves from 50 F2 *igp*$^{AEQ}$ x Col-0$^{AEQ}$ segregating plants with an impaired response to CEL3, together with leaves from the Col-0$^{AEQ}$ control line, were harvested and pooled. Genomic DNA was extracted for whole-genome sequencing to identify Single Nucleotide Polymorphism (SNPs) associated with the *igp*$^{AEQ}$ phenotypes. Sequencing (150 bp pair-end reads) was performed on an Illumina platform (Macrogen, Seoul, South Korea) to reach a coverage of 30 million reads (https://www.bioinformatics.babraham.ac.uk/projects/fastqc/). Reads were aligned against *Arabidopsis thaliana* TAIR10 genome reference using HISAT2 with standard parameters except for-no-spliced-alignment (*85*) Reads mapping to multiple locations were removed using samtools v1.7 (parameter -q 5), duplicated reads were removed using picard-tools Mark Duplicates (http://broadinstitute.github.io/picard) and indels were realigned using GATK Realigner-Target-Creator and Indel-Realigner v3.8 (*86*). Variants were called with GATK Unified-Genotyper with default parameters and the resulting files filtered for bi-allelic SNPs using bcftools (*87*). We removed variants present in the Col-0$^{AEQ}$ from the pooled mutant samples and calculated frequencies of alternate reads for each variant in the mutant pool. Frequencies were plotted along the chromosome using R and genomic regions with high frequencies of alternate alleles were further studied to look for candidate genes. Point mutations of *igp1-2*$^{AEQ}$ to *igp1-5*$^{AEQ}$ alleles were confirmed by PCR amplification of the genomic DNA of *IGP1* gene in these mutants and sequencing using oligonucleotides indicated in Table S2. *igp1-6* is a knock-out mutant (SALK_101924C) as proved by qRT-PCR using oligonucleotides indicated in Table S2.

**Plant material and generation of transgenic lines**

The *igp1-1*$^{AEQ}$ mutant line (*17*) was complemented using the following constructs: *pUBQ10::IGP1-3xHA*, *pUBQ10::IGP1*$_{D74A.N126A.D174A}$*-3xHA*, *pUBQ10::IGP1*$_{D829N}$*-3xHA*. Additionally, the *igp1-6* T-DNA insertion mutant (SALK-101924C) was complemented with *pUBQ10::IGP1-3xHA*, *pUBQ10::IGP1*$_{D74A.N126A.D174A}$*-3xHA*, *pUBQ10::IGP1*$_{D829N}$*-3xHA,* *pUBQ10::IGP1*$_{Y196A}$*-3xHA* and *pUBQ10::IGP1*$_{W146A}$*-3xHA*. The IGP1 coding sequence was amplified from *Arabidopsis thaliana* wild-type ecotype Columbia (Col-0) cDNA. Colony screening of over 100 clones consistently revealed only splicing variant 1 as susceptible to cloning (fig. S12). This splicing variant localizes correctly to the membrane, retains all essential kinase domains necessary for enzymatic activity, and maintains signaling capacity upon ligand perception (fig. S12 and S13). Protein expression of IGP1 and its variants in all transgenic lines was assessed by anti-HA-HRP conjugated (Biotec, 130-091-972) immunoblot analysis (fig. S12). To generate an IGP1 transcriptional reporter line, 2527 bp promoter region upstream of IGP1 ATG codon was cloned and fused to NLS-3xmGFP (*pIGP1::3xNLS-mGFP*) in the *Arabidopsis thaliana* Col-0 background. Constructs were generated using the GreenGate cloning system (*88*) and introduced into *Agrobacterium tumefaciens*. Transgenic *Arabidopsis thaliana* plants were then generated via the floral dip transformation method. Transgenic lines were selected using the Fast Green selection marker. The primers used for cloning are listed in Table S2.

**$Ca^{2+}$ response assays**

$Ca^{2+}$ measurements were performed in 96-well plates (1 seedling per well) under long-day conditions (14 hours of light) at 19ºC-22ºC in liquid 1/2 MS medium as described previously (*17*). Seedlings were grown *in vitro* for 8 days. then cytoplasmic $Ca^{2+}$ influxes were assessed, after loading the seedlings o/n with 10 µM coelenterazine, using a Varioskan Lux Reader luminometer (Thermo Scientific) upon treatment with the corresponding sugars indicated in each experiment. Total $Ca^{2+}$ discharge was performed by treating seedlings with 1M $CaCl_2$ and the $Ca^{2+}$ burst was quantified using the luminometer specified in (*17*). $Ca^{2+}$ discharge values were used to standardize the $Ca^{2+}$ burst between the different genotypes tested upon treatment with the different MAMPs/DAMPs.

**MAPK assays**

MAPK activation was determined in 10-day-old Arabidopsis seedlings grown on half-strength liquid MS medium in long day conditions (16 hours light, 22ºC) and treated either with water (mock) or different oligosaccharides for 15 minutes. The sugar concentrations used were 10 µM in the case of cellotriose, maltotriose, cellopentaose and cellotriose oxidized; and 50 µM for chitohexaose. Then, seedlings were flash-frozen and homogenized in the following protein extraction buffer (50 mM Tris-HCl, 200 mM NaCl, 1 mM EDTA, 10 mM NaF, 2 mM sodium orthovanadate, 1 mM sodium molybdate, 10% (v/v) glycerol, 0.1% (v/v) Tween-20, 1 mM 1,4-dithiothreitol, 1 mM phenylmethylsulfonyl fluoride and phosphatase inhibitor cocktail, pH 7.5). Total protein amounts were quantified by Bradford assay. 30 µg of proteins were separated in 10% acrylamide SDS-PAGE and transferred to nitrocellulose membranes using the Invitrogen iBlot Gel Transfer Device. Membranes were blocked with Protein-Free Blocking Buffer ( Thermo Fisher Scientific, 37571) for 2 hours at room temperature. Membranes were incubated overnight at 4°C in blocking solution, containing phospho-p44/42 MAPK (Erk1/2) (Thr202/Tyr204) antibody (Cell Signaling Technology, 9101) (1:1000). Membranes were washed with TBS-T and incubated with goat-anti-rabbit-HRP (Agrisera, AS09 602) (1:5000) in TBS-T. Blots were finally developed using the ECL western blotting substrate (Cytiva, RPN2235) and imaged using an ImageQuant 800 (Amersham). Membranes were also stained with Ponceau-S Red (AppliChem, A2935.0500).

**Transient protein expression of IGP1 wild-type and variants in Tobacco leaves.**

*Nicotiana benthamiana* plants were grown for 3 weeks before agroinfiltration. *Agrobacterium tumefaciens* cultures transformed with the constructs: *pUBQ10::IGP1-mGFP*, *pUBQ10::IGP1*$_{D74A.N126A.D174A}$*-mGFP*, *pUBQ10::IGP1$_{D829N}$-mGFP*, *pUBQ10::IGP1$_{Y196A}$-mGFP*, *pUBQ10::IGP1$_{W146A}$-mGFP* and *p35S::Lti6b-mCherry;* were grown in yeast extract broth media at 28 °C until reaching an optical density of 0.8. Cultures were centrifuged at 3,000 rpm for 10 min, and the bacterial pellets were resuspended in fresh infiltration medium (10 mM MES, 10 mM $MgCl_2$, 0.5% [m/v] saccharose, 100 µM acetosyringone, pH 5.6). The suspensions were incubated on the shaker for 2 h in the dark before syringe infiltration into N. *benthamiana* leaves. Plants were grown for an additional three days before imaging. For the plasmolysis experiments, leaves were then infiltrated with a 20% glycerol solution and imaged immediately at 400X magnification. GFP fluorescence from IGP1-GFP wild-type and mutant variants was detected using an excitation wavelength of 488 nm and an emission range of 495-545 nm. The plasma membrane marker Lti6b-mCherry was excited at 561 nm, with emission recorded between 590 and 652 nm. Imaging was performed using a Leica Stellaris 5 confocal microscope, and images were processed with ImageJ.

**Gene expression analyses by quantitative RT–qPCR**

*Arabidopsis thaliana* seedlings were grown in liquid ½ MS medium supplemented with 0.25% sucrose under long-day conditions (16 h light/8 h dark, 22 °C). IGP1 transcript levels were analyzed in 10 day-old seedlings from *igp1* mutant lines and pUBQ10-driven T2/T3 complementation lines (fig. S12). For defense-associated gene expression analyses, 5-day-old Col-0 and *igp1-6* seedlings were treated with 100 µM CEL3 for 0, 1, 2, or 4 h and immediately flash-frozen in liquid nitrogen. Total RNA was isolated from 20 mg of tissue using the ReliaPrep™ RNA Tissue Miniprep kit (Promega). First-strand cDNA was synthesized from 1 µg of RNA using the GoScript™ Reverse Transcription System (Promega). Quantitative RT–PCR was performed on a QuantStudio 3 Real-Time PCR System (Thermo Fisher Scientific) using Brilliant III Ultra-Fast SYBR Green mix (Agilent). Transcript abundance was calculated using the $2^{-\Delta\Delta CT}$ method, normalized to SAND (At2G28390) as the reference gene. Primer sequences are provided in Supplementary Table S2. Experiments were performed using several independent biological replicates, each with three technical replicates.

**ROS measurement in *Arabidopsis thaliana* seedlings upon elicitor treatment**

Seeds from *Arabidopsis thaliana* wild-type (Col-0), *igp1* mutants (*igp1-1$^{AEQ}$* or *igp1-6*, and IGP1 mutant variant complementation lines, were stratified (4ºC, two days) and germinated in white, flat-bottom polystyrene 96-well plates (BRAND® microplate BRANDplates®, pureGrade, BR781665) in ½ Murashige and Skoog (MS) liquid medium with 0.25% sucrose. They were grown in a long-day chamber (16 h light, 8 h dark, 22ºC) for 5 days. The ½ MS liquid medium was then replaced with sterile $H_2O$ and left overnight to stabilize. Luminol (L-012, Sigma-Aldrich, SML2236, 0.2 mM final concentration) and horseradish peroxidase (HRP, Sigma-Aldrich, P8375, 20 µg/mL final) were added, and ROS levels were measured in Hidex Sense Multimode Plate Reader. Cellotriose was dissolved in sterile $H_2O$ and used at a final concentration of either 10 µM or 100 µM, as indicated in the different experiments. The PAMPs used in this study were: flg22 (QRLSTGSRINSAKDDAAGLQIA, reconstituted in 100 mM NaCl), elf18 (acetyl-SKEKFERTKPHVNVGTIG, solubilized initially in DMSO and diluted in 1% w/v BSA and 100 mM NaCl), *Ralstonia solanacearum* elf18 (acetyl-AKEKFERTKPHVNVGTIG, reconstituted initially in DMSO and diluted in 1% w/v BSA and 100 mM NaCl) and Hexaacetyl-chitohexaose (Megazyme, CAS#: 38854-46-5, solubilized in 100 mM NaCl). For each plate, $n \geq 10$ seedlings were measured for ROS activity, and three technical replicates were performed for each condition.

**Callose stains on seedlings**

To test whether cellotriose could induce callose without PAMP elicitation, 7-day old *Arabidopsis thaliana* seedlings grown on solid ½ MS were transferred to liquid ½ MS with 0.25% sucrose. The seedlings were treated with 100 µM CEL3 or mock ($H_2O$) and left to develop callose for 16 hours. They were fixed in 3:1 ethanol: glacial acetic acid at room temperature for at least 30 min, were washed in sterile $H_2O$ three times and submerged in 0.1% (w/v) aniline blue o/n at room temperature before being mounted on a slide with water. To assess cellotriose's capacity to enhance callose deposition following MAMP recognition seedlings from the ROS assays underwent the same treatment described above. Aniline blue images were taken using Leica Thunder fluorescence microscope (DM6BZ) (laser 405 nm, 20x objective). Images of the roots only treated with cellotriose (fig. S15) were taken on a Leica Stellaris (laser 405 nm, 20x objective).

**Confocal settings and image processing**

Arabidopsis thaliana Col-0 and transgenic NLS-3xmVenus lines (*pFLS2$_{long}$, pEFR and pCERK1*) from Zhou et al., 2020 (*44*), were gas sterilized, stratified and germinated on either ½MS agar or ½ MS liquid plate without sucrose. Plants were grown in a long-day chamber (16 h light, 8 h dark, 22ºC) for five days before being transferred to ½ MS liquid plates for cellotriose treatment or grown for six days in ½ MS liquid and then treated with 100 µM cellotriose for 6 hours. Following this, fixation and staining were performed using the ClearSee protocol (*89*). Seedlings were fixed in ~4% paraformaldehyde/PBS solution, washed in PBS, and cleared in ClearSee solution for at least 72 hours on a shaker. The samples were then incubated with 0.1% calcofluor white to stain the cell wall. After rinsing, samples were mounted in ClearSee solution for observation under a Leica Stellaris 5 confocal microscope. Images were taken using a 40× oil immersion objective, and Z-stacks were acquired with a step size of 0.5 µm. The excitation and detection windows were as follows: mVenus (514 nm, 520–565 nm) and calcofluor white (405 nm, 415–435 nm). Expression of IGP1 was analyzed using the transcriptional reporter line *pIGP1::3xNLS-mGFP,* both in seedlings and adult plants. GFP fluorescence was detected using an excitation wavelength of 488 nm and an emission range of 502–535 nm. Calcofluor white was used to visualize cell outlines. To quantify fluorescence intensity in the *pFLS2$_{long}$::NLS-3xmVenus* line, confocal images were analyzed using the Fiji software package. After adjusting the threshold values for both mock (control)- and cellotriose-treated seedlings, regions of interest (ROIs) were selected based on gray value intensity to outline the nucleus. The total fluorescence intensity within each selected ROI was measured by calculating the integrated density (total area × mean intensity). Measurements were taken for both mock- and cellotriose-treated samples, with n = 3 seedlings per condition.

**Root growth assays**

*Arabidopsis thaliana* (Col-0) seedlings were grown on 1/2 MS media in long day conditions (16 hours light). Seedlings were transferred after four days to new plates with their respective treatments (mock (H2O), cellotriose (100 µM), PEP1 (1 µM) and maltotriose (100 µM); N ≥ 29 per treatment). Root length was measured on the day of transfer and 3 days after, with the SNT plugin for ImageJ/Fiji software (*90*). Relative growth was calculated and statistical differences were determined by Kruskal-Wallis test followed by Dunn post-hoc test (Bonferroni adjusted).

**Chemical synthesis and analysis of cellotrionic acid (CEL3ox)**

Chemicals used were purchased from Sigma-Aldrich and Fluka and were used as supplied without any further purification unless stated otherwise. Water was taken from a Milli-Q system. For reactions demanding anhydrous conditions, anhydrous solvents were produced by desiccation over freshly activated 4Å molecular sieves for at least 24 h prior to use, Activation of 4Å molecular sieves was performed by heating up the molecular sieves in a Kugelrohr apparatus (Büchi Glass Oven B-580) at 300°C in fine vacuum (typically 0.002 mbar) for at least 30 min before use and subsequently handled under an atmosphere of argon. Residual water content for solvents was determined by coulombometric titration on a Mitsubishi CA-21 Karl Fischer apparatus and was well below 10 ppm. Thin layer chromatography (TLC) was performed using pre-coated TLC-plates SIL G-25/UV254 on glass support from Macherey-Nagel. Visualization was performed using an UV-lamp (254 nm) and subsequent p-anisaldehyde dipping stain (CH-stain) followed by heating to 250°C on a hot plate.

Normal-phase flash chromatography was performed on silica gel using Fluka Kieselgel 60 (230-400 mesh) and pre-packed silica gel colums SI-S-2G/6 from Interchim. Filtrations were performed using Minisart® SRP4 Syringe Filter (PTFE, 0.45 µm) from Sartorius. Celite aided filtrations employed celite 545 from Macherey-Nagel. Organic solvents were removed in vacuo at 40 °C (unless stated otherwise) using rotary evaporators. Volatiles were removed from compounds in fine vacuum for several hours to overnight. Lyophilizations were performed using a Speedvac system at $0.3 \cdot 10^{-3}$ mbar.

NMR spectra were recorded at 297 K in the solvent indicated, using a Bruker Avance III 600 (600.22 MHz for $^{1}$H, 150.93 MHz for $^{13}$C), employing standard software provided by the manufacturer (Bruker TopSpin 3.5 pl 6). $^{1}$H and $^{13}$C spectra were referenced to the corresponding residual solvent peak chemical shift as internal standard ($CDCl_3$: 7.26 ppm $^{1}$H, 77.16 ppm $^{13}$C; $CD_3OD$: 3.31 ppm $^{1}$H, 49.00 ppm $^{13}$C) (*91*).

1H and 13C spectra in D2O were referenced via external calibration to sodium 3-(trimethylsilyl)propane-1-sulfonate (DSS)/dioxane (0.00 ppm 1H, 67.19 ppm 13C) in D2O. Assignments were supported by 1H-1H-COSY, 1H-13C-HSQC, 1H-13C-HMBC and if required CLIP-HSQC and 1H-1H-TOCSY. Anomeric configuration was verified by the 1H-13C coupling constant (*92*). Data processing and the readout of absolute integrals was performed using Bruker TopSpin 3.5 pl 6. Presented NMR-data was processed with MestReNova 16.0.0-39276. LC-MS analysis was performed on a Shimadzu LC10 system with Shimadzu 2020 mass spectrometer and Alltech ELSD 3300 (drift tube temperature: 60°C, receiver gain: 2) using a Hypercarb™ Porous Graphitic Carbon column (100x4.6 mm, 5 µm particle size. Data was processed with the software provided by the manufacturer. Preparative HPLC separation was performed on an Interchim PuriFlash® 4/25 (Flash and preparative HPLC system) using a YMC-pack SIL-06 (250 x 10 mm, D. 5 µm, 6 nm). ESI-HRMS data was obtained using samples dissolved in MeCN/H2O on an Agilent Technologies 6230B LCMS-TOF or a Waters Xevo G2-XS QTof instrument. Datasets were analysed by Mass Hunter Qualitative Navigator 10.00 software or mass-adducts were calculated with Mass Hunter Isotope Distribution Calculator v. 8.0.8208.0 software and the ppm-differences calculated (< ±2.5 ppm).

Preparation of starting material D-cellotriose undecaacetate (Step **1**):

As a starting material a crude mix of low DP (n = 1-30) peracetylated oligocelluloses from $Ac_2O/H_2SO_4$ treated cellulose was used (*56*). The mixture of peracetylated oligocelluloses was suspended in toluene:EtOAc 40/60 and filtered over a plug of celite. This led to the clean elution of mono- and oligomers (n= 1-8), while higher oligo- and polymers remained in a microcrystalline layer that could later be dissolved efficiently with acetone. After concentration of the toluene/EtOAc filtrate the resulting oligosaccharide mixture was purified by column chromatography (product/$SiO_2$ 1:100, toluene/acetone 80/20 to 75/25) to obtain pure peracetylated cellotriose (*91*) as an α/β mixture. Analytical data is in full agreement with literature reported values (*93*).

2,3,4,6-Tetra-*O*-acetyl-β-D-glucopyranosyl-(1→4)-2,3,6-tri-*O*-acetyl-β-D-glucopyranosyl-(1→4)-2,3,6-tri-*O*-acetyl-D-glucopyranoside (Step **2**)

Per-Ac-cellotriose **1** (100 mg, 103 µmol, 1.00 eq) was dissolved in anhydrous THF (1.0 mL) and a freshly prepared suspension of AcOH (8.3 µL, 145 µmol, 1.40 eq) and ethylene diamine (8.3 µL, 124 µmol, 1.20 eq) in anh. THF (1.6 mL) was immediately added. The suspension was stirred overnight (15.5 h). After confirming completion of the reaction by TLC, the mixture was partitioned between EtOAc (10 mL) and water (10 mL). Phases were separated and the organic layer was washed with 2 M HCl, sat. aq. $NaHCO_3$ and brine. After drying over $Na_2SO_4$ it was filtered over glass wool and concentrated to obtain a colorless crude (99 mg). The crude was purified by flash chromatography (2 g $SiO_2$ prepacked, 80/20 toluene/acetone to 70/30). Mixed fractions were treated in the same manner. Hemiacetal **2** (71.0 mg, 74%) was obtained after concentration as a colorless foam. Analytical data is in full agreement with literature reported values (*94*).

2,3,4,6-Tetra-*O*-acetyl-β-D-glucopyranosyl-(1→4)-2,3,6-tri-*O*-acetyl-β-D-glucopyranosyl-(1→4)-2,3,6-tri-*O*-acetyl-D-glucopyranolactone (Step **3**)

Under an atmosphere of argon, hemiacetal **2** (30.0 mg, 32.4 µmol, 1.00 eq) and Dess-Martin periodinane (70 mg, 164 µmol, 5.05 eq) were dissolved in anhydrous $CH_2Cl_2$ (1.0 mL) and the solution was stirred at rt for 4 h and 45 min. The reaction was quenched by addition of a mixture of sat. aq. $NaHCO_3$ and 20 % aq. $NaS_2O_3$ (10 mL, 50/50). Within 5 min all solids dissolved and phases were separated. The aqueous layer was washed with $CH_2Cl_2$ (4x) and the combined organic layer was dried over $Na_2SO_4$ and filtered over glass wool. Concentration gave a colorless crude product (29 mg) which was purified by flash chromatography (2 g $SiO_2$ prepacked, 60/40 toluene/acetone with 1% AcOH) to give lactone **3** as a colorless solid (20.0 mg, 67%). TLC: Rf = 0.4 ($SiO_2$, 60/40 toluene/acetone with 1% AcOH). **$^1$H NMR** ($CDCl_3$, 600 MHz): $\delta$ = 5.52 (dd, $J_{3\text{-}2}$ = 7.5 Hz $J_{3\text{-}4}$ = 7.7 Hz, 1 H, H-3$^{glc\text{-}1}$), 5.14-5.10 (m, 2 H, H-3$^{glc\text{-}2}$ H-3$^{glc\text{-}3}$), 5.10-5.03 (m, 2 H, H-2$^{glc\text{-}1}$ H-4$^{glc\text{-}3}$), 4.90 (dd, $J_{2\text{-}1}$ = 7.9 Hz $J_{2\text{-}3}$ = 9.3 Hz, 1 H, H-2$^{glc\text{-}3}$), 4.86 (dd, $J_{2\text{-}1}$ = 7.8 Hz $J_{2\text{-}3}$ = 9.3 Hz, 1 H, H-2$^{glc\text{-}3}$), 4.60 (d, $J_{2\text{-}1}$ = 7.8 Hz 1 H, H-1$^{glc\text{-}2}$), 4.58 (ddd, $J_{5\text{-}6a}$ = 2.5 Hz $J_{5\text{-}6b}$ = 4.3 Hz $J_{5\text{-}4}$ = 8.6 Hz, 1 H, H-5$^{glc\text{-}1}$), 4.52-4.48 (m, 2 H, H-1$^{glc\text{-}3}$ H-6a$^{glc\text{-}1}$), 4.45 (dd, $J_{6a\text{-}5}$ = 2.0 Hz $J_{6a\text{-}6b}$ = 12.1 Hz, 1 H, H-6a$^{glc\text{-}2}$), 4.35 (dd, $J_{6a\text{-}5}$ = 4.4 Hz $J_{6a\text{-}6b}$ = 12.5 Hz, 1 H, H-6a$^{glc\text{-}3}$), 4.22 (dd, $J_{6b\text{-}5}$ = 4.4 Hz $J_{6b\text{-}6a}$ = 12.5 Hz, 1 H, H-6b$^{glc\text{-}1}$), 4.10 (dd, $J_{6b\text{-}5}$ = 4.9 Hz $J_{6b\text{-}6a}$ = 12.2 Hz, 1 H, H-6b$^{glc\text{-}2}$), 4.06-4.00 (m, 2 H, H-6b$^{glc\text{-}3}$ H-4$^{glc\text{-}1}$), 3.79 (dd, $J_{4\text{-}3}$ = 9.5 Hz $J_{4\text{-}5}$ = 9.3 Hz, 1 H, H-4$^{glc\text{-}2}$), 3.66-3.61 (m, 2 H, H-5$^{glc\text{-}3}$ H-5$^{glc\text{-}2}$), 2.15 (s, 6 H), 2.13 (s, 3 H), 2.08 (s, 3 H), 2.06 (s, 3 H), 2.03 (m, 6 H), 2.01 (s, 6 H), 1.98 (s, 3 H); **$^{13}$C NMR** ($CDCl_3$, 150 MHz): $\delta$ = 170.6 170.31 170.29 170.2 169.8 169.7 169.42 169.41 169.3 169.2 (C=O$^{Ac}$), 164.6

(C-1$^{Glc-1}$), 100.9 (C-1$^{Glc-3}$), 100.5 (C-1$^{Glc-2}$), 76.4 (C-5$^{Glc-1}$), 76.1 (C-4$^{Glc-2}$), 74.7 (C-4$^{Glc-1}$), 73.1 (C-5$^{Glc-2}$), 73.0 (C-3$^{Glc-3}$), 72.6 (C-3$^{Glc-2}$), 72.2 (C-5$^{Glc-3}$), 71.74 (C-2$^{Glc-3}$), 71.70 (C-2$^{Glc-2}$), 70.5 (C-3$^{Glc-1}$), 70.2 (C-2$^{Glc-1}$), 67.9 (C-4$^{Glc-3}$), 62.0 (C-6$^{Glc-2}$), 61.65 (C-6$^{Glc-1}$), 61.60 (C-6$^{Glc-3}$), 20.9 20.8 20.69 20.68 20.65 20.60 ($CH_3^{Ac}$); **HRMS**: calcd. for $C_{38}H_{50}O_{26}$: 923.2663 [$M$+H]$^+$, found 923.2647.

Sodium 2,3,4,6-tetra-*O*-acetyl-β-D-glucopyranosyl-(1→4)-2,3,6-tri-*O*-acetyl-β-D-glucopyranosyl-(1→4)-2,3,6-tri-*O*-acetyl-D-gluconate (Step **4**)

In a screw cap Eppendorf tube, lactone **3** (16.8 mg, 18.2 µmol, 1.00 eq) was dissolved in a mixture of MeCN and $H_2O$ (1.0 mL, 50/50), and $NaHCO_3$ (1.49 mg, 17.7 µmol, 0.97 eq) was added. The solution was mixed by shaking and allowed to stand for 5 h, and then lyophilized. The obtained colorless powder was taken up in EtOAc (50 µL) and addition of toluene (1.0 mL) led to precipitation. After centrifugation and decanting off the liquid, the solid was redissolved, precipitated two more times as described and finally dried in fine vacuum. The obtained pale brown powder was dissolved in $H_2O$ (200 µL) and the solution was passed through a syringe filter (Satorius 0.45 µm) followed by washing of the filter with $H_2O$ (3x 100 µL). The clear filtrate was lyophilized to give acetylated sodium cellotrionate **4** (16.0 mg, 94 %). **$^1$H NMR** ($CD_3OD$, 600 MHz): $\delta$ = 5.63 (dd, $J_{3\text{-}2}$ = 3.5 Hz $J_{3\text{-}4}$ = 7.3 Hz, 1 H, H-3$^{glc-1}$), 5.23 (d, $J_{2\text{-}3}$ = 3.5 Hz, 1 H, H-2$^{glc-1}$), 5.19 (dd, $J_{3\text{-}2}$ = $J_{3\text{-}4}$ = 9.4 Hz, 1 H, H-3$^{glc-3}$), 5.13 (dd, $J_{3\text{-}2}$ = 9.4 Hz $J_{3\text{-}4}$ = 9.2 Hz, 1 H, H-3$^{glc-2}$), 4.99 (dd, $J_{4\text{-}3}$ = 9.5 Hz $J_{4\text{-}5}$ = 10.1 Hz, 1 H, H-4$^{glc-3}$), 4.87 (d, $J_{2\text{-}1}$ = 8.0 Hz, 1 H, H-1$^{glc-2}$), 4.81-4.75 (m, 2 H, H-2$^{glc-3}$ H-2$^{glc-2}$), 4.68 (d, $J_{5\text{-}4}$ = 8.0 Hz, 1 H, H-1$^{glc-3}$), 4.45 (dd, $J_{6a\text{-}5}$ = 2.0 Hz $J_{6a\text{-}6b}$ = 12.0 Hz, 1 H, H-6a$^{glc-2}$), 4.38 (dd, $J_{6a\text{-}5}$ = 4.2 Hz $J_{6a\text{-}6b}$ = 12.5 Hz, 1 H, H-6a$^{glc-3}$), 4.23 (dd, $J_{6a\text{-}5}$ = 3.6 Hz $J_{6a\text{-}6b}$ = 11.5 Hz, 1 H, H-6a$^{glc-1}$), 4.18 (dd, $J_{6a\text{-}5}$ = 5.8 Hz $J_{6a\text{-}6b}$ = 12.1 Hz, 1 H, H-6b$^{glc-2}$), 4.12 (dd, $J_{4\text{-}3}$ = 7.4 Hz $J_{4\text{-}5}$ = 4.3 Hz, 1 H, H-4$^{glc-1}$), 4.08-4.02 (m, 2 H, H-6b$^{glc-1}$ H-6b$^{glc-3}$), 3.93 (ddd, $J_{5\text{-}6a}$ = 3.6 Hz $J_{5\text{-}4}$ = 4.3 Hz $J_{5\text{-}6b}$ = 7.5 Hz, 1 H, H-5$^{glc-1}$), 3.87 (ddd, $J_{5\text{-}6b}$ = 2.2 Hz $J_{5\text{-}6a}$ = 4.2 Hz $J_{5\text{-}4}$ = 10.2 Hz, 1 H, H-5$^{glc-3}$), 3.83 (dd, $J_{4\text{-}3}$ = 9.2 Hz $J_{4\text{-}5}$ = 10.0 Hz, 1 H, H-4$^{glc-2}$), 3.73 (ddd, $J_{5\text{-}6a}$ = 2.2 Hz $J_{5\text{-}6b}$ = 4.2 Hz $J_{5\text{-}4}$ = 10.0 Hz, 1 H, H-5$^{glc-2}$), 2.13 (2 s, 6 H), 2.08 (s, 3 H), 2.05 (s, 3 H), 2.03-2.02 (3 s, 9 H), 2.00 (s, 3 H), 1.98 (s, 3 H), 1.93 (s, 3 H); **$^{13}$C NMR** ($CD_3OD$, 150 MHz): $\delta$ = 173.6 (C-1$^{Glc-1}$), 172.9 172.6 172.19 172.17 171.9 171.6 171.4 171.2 171.0 (C=O$^{Ac}$), 102.4 (C-1$^{Glc-2}$), 101.9 (C-1$^{Glc-3}$), 82.0 (C-4$^{Glc-1}$), 78.0 (C-4$^{Glc-2}$), 75.0 (C-2$^{Glc-1}$), 74.6 (C-3$^{Glc-2}$), 74.5 (C-3$^{Glc-3}$), 73.8 (C-5$^{Glc-2}$), 73.5 (C-2$^{Glc-2}$), 73.1 (C-2$^{Glc-3}$), 72.9 (C-5$^{Glc-3}$), 72.7 (C-3$^{Glc-1}$), 70.3 (C-5$^{Glc-1}$), 69.3 (C-4$^{Glc-3}$), 66.3 (C-6$^{Glc-1}$), 63.8 (C-6$^{Glc-2}$), 62.8 (C-6$^{Glc-3}$), 21.03 21.01 20.92 20.87 20.81 20.65 20.61 20.52 20.49 ($CH_3^{Ac}$); **HRMS**: calcd. for $C_{38}H_{52}O_{27}$: 939.2623 [$M$-H]$^-$, found 939.2622.

Sodium β-D-glucopyranosyl-(1→4)-β-D-glucopyranosyl-(1→4)-D-gluconate (Step **5**)

To the sodium salt **4** dissolved in a NMR tube in MeOH-$D_4$ (0.75 mL) was added $NEt_3$ (37.5 µL, 5% vol.) and the mixture was allowed to stand at 50°C for 24 h. Precipitate had formed, so the mixture was diluted with $H_2O$ and transferred to a screw cap Eppendorf tube, followed by washing of the NMR tube with $H_2O$. After lyophilization of the solution, the residue was taken up in $D_2O$ with 5% $NEt_3$ (37.5 µL). After 16 h at rt, residual acetyl groups had been cleaved off as indicated by $^1$H-NMR, so the reaction mixture was lyophilized. The obtained brown crude product (11.4 mg) was taken up in $H_2O$ (50 µL), and addition of *i*PrOH (1.0 mL) led to precipitation. After centrifugation and decanting off the liquid, the solid was redissolved and precipitated five more times as described, and finally taken up in $H_2O$ (0.1 mL) and lyophilized. The obtained beige powder was redissolved in $H_2O$ (0.25 mL) and passed through a syringe filter (satorius 0.45 µm) followed by washing of the filter with water (3x 0.25 mL). The filtrate was lyophilized and finally redissolved in $H_2O$ (1 mL) and lyophilized two more times to remove any volatile organic compounds. The product sodium cellotrionate **5** (9.11 mg, quant.) was obtained as a colorless powder. **$^1$H NMR** ($D_2O$, 600 MHz): $\delta$ = 4.63 (d, $J_{1\text{-}2}$ = 7.9 Hz, 1 H, H-1$^{glc\text{-}2}$), 4.49 (d, $J_{1\text{-}2}$ = 8.0 Hz, 1 H, H-1$^{glc\text{-}3}$), 4.13 (d, $J_{2\text{-}3}$ = 3.1 Hz, 1 H, H-2$^{glc\text{-}1}$), 4.07 (dd, $J_{3\text{-}2}$ = 3.1 Hz $J_{3\text{-}4}$ = 5.5 Hz, 1 H, H-3$^{glc\text{-}1}$), 3.99 (dd, $J_{3\text{-}4}$ = 5.4 Hz $J_{4\text{-}5}$ =5.3 Hz, 1 H, H-4$^{glc\text{-}1}$), 3.975-3.93 (m, 2 H, H-5$^{glc\text{-}1}$ H-6a$^{glc\text{-}2}$), 3.90 (dd, $J_{6a\text{-}5}$ = 2.4 Hz $J_{6a\text{-}6b}$ = 12.5 Hz, 1 H, H-6a$^{glc\text{-}3}$), 3.83 (dd, $J_{6a\text{-}5}$ = 3.5 Hz $J_{6a\text{-}6b}$ = 12.2 Hz, 1 H, H-6a$^{glc\text{-}1}$), 3.80 (dd, $J_{6b\text{-}5}$ = 5.2 Hz $J_{6b\text{-}6a}$ = 12.4 Hz, 1 H, H-6b$^{glc\text{-}2}$), 3.75-3.70 (m, 2 H, H-6b$^{glc\text{-}1}$ H-6b$^{glc\text{-}2}$), 3.67-3.61 (m, 2 H, H-4$^{glc\text{-}2}$ H-3$^{glc\text{-}2}$), 3.57 (ddd, $J_{5\text{-}6a}$ = 2.4 Hz $J_{5\text{-}6b}$ = 5.3 Hz $J_{5\text{-}4}$ = 7.5 Hz, 1 H, H-5$^{glc\text{-}2}$), 3.51-3.45 (m, 2 H, H-3$^{glc\text{-}3}$ H-5$^{glc\text{-}3}$), 3.42-3.35 (m, 2 H, H-4$^{glc\text{-}3}$ H-2$^{glc\text{-}2}$), 3.29 (dd, $J_{2\text{-}1}$ = 7.9 Hz $J_{2\text{-}3}$ = 9.4 Hz, 1 H, H-2$^{glc\text{-}3}$); **$^{13}$C NMR** ($CDCl_3$, 150 MHz): $\delta$ = 179.0 (C-1$^{Glc\text{-}1}$), 103.4 (C-1$^{Glc\text{-}2}$), 103.2 (C-1$^{Glc\text{-}3}$), 82.2 (C-4$^{Glc\text{-}1}$), 79.1 (C-4$^{Glc\text{-}2}$), 76.6 (C-5$^{Glc\text{-}3}$), 76.1 (C-3$^{Glc\text{-}3}$), 75.4 (C-5$^{Glc\text{-}2}$), 74.8 (C-3$^{Glc\text{-}2}$), 73.82 (C-2$^{Glc\text{-}2}$), 73.76 (C-2$^{Glc\text{-}3}$), 73.0 (C-2$^{Glc\text{-}1}$), 72.4 (C-5$^{Glc\text{-}1}$), 72.2 (C-3$^{Glc\text{-}1}$), 70.1 (C-4$^{Glc\text{-}3}$), 62.6 (C-6$^{Glc\text{-}1}$), 61.2 (C-6$^{Glc\text{-}3}$), 60.6 (C-6$^{Glc\text{-}2}$); **HRMS**: calcd. for $C_{18}H_{32}O_{17}$: 519.1567 [*M*-H]$^-$, found 519.1566; **HPLC** (2.5-40% MeCN + 0.1% AcOH in 25 min, 0.7 mL/min): $R_t$ = 29.289 min, $R_{t\ lactone}$ = 23.330 min.

Purity determination of sodium cellotrionate 5 using external NMR-calibration:

The exact amount of sodium salt **5** in the final sample was determined by external NMR calibration with raffinose pentahydrate. Using identical NMR parameter sets with a fixed receiver gain and 32 scans, absolute integrals for an anomeric proton peak were determined for raffinose samples in D2O (0.97, 2.26, 3.53, 5.24, 8.08, 11.22, 14.59 mg/mL) and the sample of sodium cellotrionate 5 (9.11 mg in 0.70 mL $D_2O$). The absolute integral of sodium cellotrionate **5** was determined at three different proton signals with a relative integral of 1 and then arithmetically averaged and concentration determined with the external calibration curve derived from the raffinose series. We found a purity of ≥90% by weight

**Cellulose digestion experiments**

Commercial cellulase R-10 (Onozuka R-10, from Trichoderma), solubilized in 50 mM sodium acetate buffer (pH 5.5), or Xanthomonas Leaf 131 or Leaf 131 *xpsxcs* exudates were used to digest 1% w/v Azo-CM-cellulose (Megazyme, Ref#: S-ACMC). After overnight incubation at room temperature, the precipitant solution (294 mM sodium acetate trihydrate, 21.8 mM zinc acetate, 80% ethanol, pH 5.0) was added at a 1:5 ratio (1 part cellulase and cellulose to 5 parts precipitant solution), briefly vortexed, and centrifuged at 5,000 × g for 5 min. The supernatants were collected and either measured in a spectrophotometer at 590 nm or used for thin-layer chromatography in a silica plate, with a solvent mixture (2:1:1, n-butanol:acetic acid:water) as the mobile phase and stained with an ethanol-anisaldehyde-sulfuric acid (18:1:1) solution. For the Congo red assay, cellulase (1 mg/mL), Xanthomonas strains, and *Pseudomonas syringae* DC3000 (*Pst* cor$^-$) at $OD_{600}$ 0.5 in $MgCl_2$, were spotted onto R2A agar medium containing 0.5% sodium carboxymethyl cellulose (Sigma-Aldrich, Ref: 419311). After 24 hours of incubation at 28°C, the agar plate was stained with 0.1% w/v Congo red dye dissolved in 50% ethanol for 30 min and destained with 1 M NaCl for 15 min. The presence of a white halo indicates cellulose degradation.

**Generation and HPAEC–PAD profiling of cellulose-derived oligosaccharides by cellulase and *Plectosphaerella cucumerina* BMM**

*Plectosphaerella cucumerina* BMM strain (*Pc*BMM) was grown on potato dextrose agar (PDA) as described in (*63*). For hydrolysis of cellulose (Sigma-Aldrich, C6288) by *Pc*BMM mycelia or cellulase from *Trichoderma reesei* ATCC 26921 (Sigma-Aldrich), cellulose was dissolved in sterilized MilliQ water at a concentration of 3 g/L. *Pc*BMM mycelium was scraped from the PDA plates and added to the cellulose solution, and then incubated at 28 °C. For the enzymatic hydrolysis, cellulase was added to the cellulose solution at a concentration of 0.33 U/mg, and incubated at 40°C. Samples from both the *Pc*BMM mycelium-incubated or cellulase treated were collected at different time points (0.15 h, 0.5 h, 4 h, and 24 h), and hydrolytic activity was stopped by incubating the harvested samples at 90º C for 5 minutes. $Ca^{2+}$ burst response of these samples was evaluated in Col-$0^{AEQ}$ and *igp1-3$^{AEQ}$* lines as described above. To study the released cello-oligosaccharides over time, high-performance anion-exchange chromatography coupled with integrated and pulsed amperometric detection (HPAEC-PAD) was performed in a Dionex ICS-6000 system operated with Chromeleon 7 software (version 7.3.1). Separation of the oligosaccharides was performed at 30 ºC on a CarboPac PA20 analytical column (3 mm x 150 mm) fixed to a CarboPac PA20 guard column (3 mm x 30 mm). Elution was realized using a binary gradient consisting of eluent A (250 mM NaOH) and eluent B (250 mM NaOAc in 100 mM NaOH), at a constant flow rate of 0.5 mL/min. Then, 500 µL of the filtered samples were injected. The following standards were used in this study: hexaacetyl-chitohexaose (CHI6; β-1,4-D-$(GlcNAc)_6$; #O-CHI6), cellotriose (CEL3; β-1,4-D-$(Glc)_3$; O-CTR), cellotetraose (CEL4; β-1,4-D-$(Glc)_4$; #O-CTE), cellopentaose (CEL5; β-1,4-D-$(Glc)_5$; #OCPE) and MLG43 (β-1,4-D-$(Glc)_2$-β-1,3-D-Glc, # O-BGTRIB) from Megazyme (Wicklow, Ireland: https://www.megazyme.com), cellobiose (CEL2; β-1,4-D-$(Glc)_2$; C7252), glucose (Glc), sucrose, maltose, trehalose from Sigma-Aldrich (https://www.sigmaaldrich.com). Experiment was repeated 3 times with similar results.

#### In vivo pretreatment with cello-oligomers and *Pseudomonas syringae* pv. tomato DC3000 pathogen infection assays

The pathogen assay with cellotriose pretreatment was adapted from Roussin-Léveillée et al., 2022 (*95*). Four- to five-week-old *Arabidopsis thaliana* plants grown under short-day conditions (10 h light/14 h dark) were syringe-infiltrated with 10 mM $MgCl_2$ (mock), 100 µM cellotriose (CEL3), 100 µM oxidized cellotriose (CEL3ox), or left untreated 24 h before inoculation with *Pseudomonas syringae* pv. tomato DC3000 (*Pst* DC3000). After infiltration, leaves were blotted dry, allowed to recover for 1–2 h in ambient conditions, and returned to the growth chamber. *Pst* DC3000 was cultured overnight in lysogeny broth with 50 mg $L^{-1}$ rifampicin, a fresh culture was inoculated the day of infection. Once the bacterial density reached $OD_{600} = 0.8$–1.0, the bacteria were pelleted (2,500 g, 3 min, $OD_{600} = 0.8$–1.0), washed once, and resuspended in 10 mM $MgCl_2$ to $OD_{600} = 0.2$, then diluted to $OD_{600} = 0.001$ ($\sim 1 \times 10^6$ CFU $mL^{-1}$) for infiltration. Pre-treated or untreated leaves were infiltrated, blotted dry, and incubated under high humidity (> 95%) by covering trays with a water-misted plastic lid. Three days post inoculation (dpi), leaves were surface-sterilized (70% ethanol, 30 s), rinsed twice in sterile water, and three 0.4 cm discs (one per leaf) were collected per plant. Discs were homogenized in sterile water, and serial dilutions ($10^1$–$10^{-6}$) were plated on LB–rifampicin plates for CFU quantification. Plates were incubated at 28°C and CFUs were counted 2 days after bacterial harvest. Each experiment included three biological replicates with three technical replicates per dilution and was repeated 3–5 times with consistent results.

#### *Ralstonia solnacearum* GMI1000 disease resistance experiments

*Arabidopsis thaliana* seeds were sown in soil (1:1 peat soil:vermiculite mixture) and grown under short-day photoperiod conditions (10h light/14h dark) at 22ºC, 65% relative humidity, and a light intensity of 100–150 µE/m²/s in a plant growth chamber for 7 days. Seedlings were then transferred to hydrated, sterilized Jiffy pots (Jiffy International group) and grown for 4 weeks under the same conditions before bacteria inoculation. *Ralstonia solanacearum* GMI1000 bacterial suspension was prepared to a final concentration of $OD_{600}$ = 0.1 (corresponding to 1 × $10^8$ CFU/ml). 300 mL of inoculum of the bacterial suspension were poured to soak the soil of 15 plants in one plastic tray. Plants were transferred from the bacterial solution to a new tray with a bed of potting mixture soil after 20-minute incubation with the bacterial inoculum (*96*). After soil drenching inoculation, plants were kept in a growth chamber with the following conditions: 75 % humidity, 12 h light, 130 µE $m^{-2}$ $s^{-1}$, 27 °C, and 12 h darkness, 26 °C for disease symptom scoring. To evaluate disease symptoms, a scale ranging from '0' (no symptoms) to '4' (complete wilting) was performed as previously described in Vailleau et al., 2007 (*65*). To perform the survival analysis, the disease scoring was transformed into binary data with the following criteria: the disease index < 2 was defined as '0', while the disease index ≥ 2 was defined as '1', in terms of the corresponding time (days post-inoculation, dpi) (*97*). Statistical analyses for survival data were performed using a Log-rank (Mantel-Cox test) and Gehan-Breaslow-Wilcoxon test.

### Inter-domain interface residues

**malectin domain**

| | HSDC | ASA | BSA | %BSA | $\Delta^{i}G$ |
|---|---|---|---|---|---|
| Lys416 | | 166.81 | 34.93 | 21% | -0.31 |
| Gly417 | | 20.83 | 11.33 | 54% | 0.08 |
| Val418 | | 39.06 | 15.24 | 39% | 0.24 |
| Tyr419 | | 4.67 | 1.25 | 27% | 0.02 |
| Leu446 | H | 107.81 | 10.68 | 10% | -0.12 |
| Gly447 | | 8.37 | 3.01 | 36% | 0.05 |
| Pro448 | | 61.22 | 48.33 | 79% | 0.66 |
| Ala449 | H | 80.81 | 80.81 | 100% | 0.61 |
| Thr450 | H | 46.62 | 26.24 | 56% | 0.08 |
| Phe451 | | 139.42 | 116.12 | 83% | 1.86 |
| Phe452 | | 72.97 | 6.42 | 9% | 0.1 |
| Val453 | H | 61.34 | 60.32 | 98% | 0.63 |
| Ser454 | H | 7.06 | 2.21 | 31% | -0.03 |
| Lys455 | | 190.72 | 29.69 | 16% | 0.4 |
| Gln457 | H | 101.17 | 58.3 | 58% | -0.06 |
| Ala460 | | 6.86 | 6.86 | 100% | 0.11 |
| Val461 | | 1.36 | 1.36 | 100% | 0 |
| Ser462 | H | 26.35 | 22.46 | 85% | -0.1 |
| Asn463 | H | 0.54 | 0.24 | 44% | 0 |
| Val464 | | 76.27 | 67.75 | 89% | 1.08 |
| Gly465 | | 8.13 | 5.66 | 70% | 0.09 |
| Leu466 | | 124.57 | 80.22 | 64% | 1.09 |
| Thr468 | | 104.57 | 72.23 | 69% | 0.59 |
| Ala499 | | 5.81 | 0.24 | 4% | 0 |
| Ser500 | H | 40.75 | 33.84 | 83% | -0.24 |
| Ser501 | | 14.4 | 11.54 | 80% | 0.17 |
| Arg503 | H | 74.41 | 61.82 | 83% | -1.32 |
| Tyr505 | H | 51.36 | 50.98 | 99% | 0.27 |
| Leu507 | | 45.49 | 41 | 90% | 0.66 |
| Gly508 | | 26.6 | 25.28 | 95% | 0.25 |
| Asn511 | H | 64.18 | 5.5 | 9% | -0.06 |
| Ser528 | | 75.86 | 0.61 | 1% | -0.01 |
| Asn529 | | 163.75 | 52.72 | 32% | -0.38 |
| Thr530 | H | 34.7 | 25.6 | 74% | 0.11 |
| Trp531 | H | 133.96 | 117.4 | 88% | 1.28 |
| Ser533 | | 30.77 | 4.3 | 14% | -0.05 |
| Leu534 | | 120 | 79.53 | 66% | 1.27 |
| Arg536 | | 102.87 | 1.69 | 2% | 0.01 |
| Ile538 | | 25.08 | 4.84 | 19% | 0.08 |
| Glu576 | HS | 123.39 | 45.36 | 37% | 0.19 |
| Asn577 | H | 42.22 | 33.51 | 79% | -0.06 |
| Tyr578 | | 86.75 | 44.94 | 52% | 0.5 |
| Glu580 | H | 34.22 | 14.73 | 43% | -0.25 |
| His582 | | 17.5 | 7.81 | 45% | 0.12 |
| Phe584 | | 71.75 | 59.4 | 83% | 0.95 |
| Trp585 | H | 74.92 | 39.47 | 53% | 0.44 |
| Ala586 | | 51.27 | 33.62 | 66% | 0.14 |
| Gly587 | | 53.23 | 52.41 | 98% | 0.08 |
| Lys588 | | 91.53 | 38.52 | 42% | 0.58 |
| Gly589 | H | 31.65 | 28.94 | 91% | -0.13 |
| Thr590 | H | 42.69 | 34.62 | 81% | 0.54 |
| Cys591 | | 68.64 | 43.36 | 63% | 1.44 |
| Cys592 | H | 50.89 | 30.22 | 59% | 0.65 |
| Ile593 | | 31.35 | 31.35 | 100% | 0.5 |
| Gln596 | | 108.44 | 20.97 | 19% | -0.36 |
| Leu620 | | 81.6 | 13.05 | 16% | 0.21 |
| Pro621 | | 161.78 | 2.51 | 2% | 0.04 |

**LRR domain**

| | HSDC | ASA | BSA | %BSA | $\Delta^{i}G$ |
|---|---|---|---|---|---|
| Gly103 | | 43.50 | 10.94 | 25% | 0.05 |
| Asp105 | H | 102.70 | 49.48 | 48% | -0.21 |
| Gln127 | | 75.04 | 48.26 | 64% | 0.12 |
| Asn128 | | 11.34 | 11.34 | 100% | -0.13 |
| Phe129 | | 129.59 | 85.33 | 66% | 1.37 |
| Ala151 | | 28.84 | 27.84 | 97% | 0.37 |
| Asn152 | | 14.21 | 12.02 | 85% | -0.01 |
| Ala153 | | 29.03 | 12.31 | 42% | 0.2 |
| Asp174 | | 22.34 | 3.04 | 14% | -0.05 |
| Met175 | H | 98.16 | 82.80 | 84% | 2.46 |
| Asn177 | | 93.21 | 1.96 | 2% | -0.03 |
| Ser199 | H | 31.21 | 18.90 | 61% | 0.04 |
| Trp220 | | 83.94 | 44.82 | 53% | 0.72 |
| Asn222 | | 37.85 | 26.35 | 70% | -0.28 |
| Asp223 | H | 85.38 | 60.51 | 71% | -0.16 |
| Arg225 | | 127.48 | 11.37 | 9% | 0.18 |
| Arg244 | | 52.64 | 23.16 | 44% | -0.53 |
| Leu246 | | 56.06 | 37.55 | 67% | 0.6 |
| Gly247 | | 36.29 | 31.82 | 88% | 0.3 |
| Glu266 | | 25.99 | 0.12 | 0% | 0 |
| Arg268 | | 61.85 | 31.50 | 51% | -0.58 |
| Glu271 | | 57.85 | 25.18 | 44% | 0.24 |
| Arg294 | H | 95.10 | 82.54 | 87% | -0.48 |
| Asn295 | | 54.24 | 23.50 | 43% | -0.18 |
| Gln314 | | 46.24 | 26.60 | 58% | -0.3 |
| Asp316 | | 10.93 | 0.86 | 8% | -0.01 |
| Ser318 | | 5.67 | 3.52 | 62% | 0.01 |
| Phe319 | | 94.56 | 66.12 | 70% | 1.06 |
| His338 | | 67.92 | 34.59 | 51% | -0.2 |
| Phe340 | | 65.25 | 59.16 | 91% | 0.95 |
| Asn343 | H | 52.67 | 13.95 | 26% | -0.21 |
| Asn360 | | 29.24 | 14.24 | 49% | -0.06 |
| Ile361 | | 3.86 | 3.86 | 100% | 0.01 |
| Asp362 | | 25.88 | 23.18 | 90% | 0.13 |
| Ser364 | | 15.36 | 11.58 | 75% | -0.08 |
| Tyr365 | H | 94.39 | 59.61 | 63% | 0.23 |
| Leu381 | | 27.46 | 0.86 | 3% | -0.01 |
| Gln382 | H | 121.92 | 82.59 | 68% | -0.67 |
| Leu383 | H | 32.56 | 28.63 | 88% | -0.28 |
| Asn384 | H | 61.32 | 61.32 | 100% | -0.27 |
| Leu385 | | 0.77 | 0.44 | 57% | 0 |
| Ile386 | | 44.00 | 43.84 | 100% | 0.7 |
| Leu400 | | 28.38 | 11.45 | 40% | 0.18 |
| Pro401 | | 83.88 | 18.63 | 22% | -0.07 |
| Arg402 | H | 168.89 | 117.01 | 69% | -1.14 |
| Leu403 | | 19.42 | 18.75 | 97% | 0.3 |
| Cys405 | | 33.66 | 30.57 | 91% | 0.66 |
| Leu406 | | 71.61 | 71.61 | 100% | 1.15 |
| Lys408 | | 76.75 | 1.96 | 3% | -0.02 |
| Phe410 | | 94.94 | 81.99 | 86% | 1.18 |
| Cys412 | | 58.07 | 39.27 | 68% | 0.54 |
| Asn413 | H | 156.53 | 118.03 | 75% | -0.59 |
| Arg414 | HS | 176.47 | 66.65 | 38% | -1.13 |
| Gly415 | H | 121.79 | 76.84 | 63% | 0.22 |

**Fig. S1. A large network of polar and hydrophobic interactions governs IGP1 multidomain stability.** Table of residues involved in the interaction between the malectin and the LRR domain of IGP1. HSDC, indicate the residues making **H**ydrogen/**D**isulphide bond, **S**alt bridge or **C**ovalent links. ASA, represents the accessible surface area in $A^2$. BSA and %BSA, depicts the buried surface area. $\Delta^{i}G$, indicate the solvation energy effect, kcal/mol, according to the PISA server analysis (*98*).

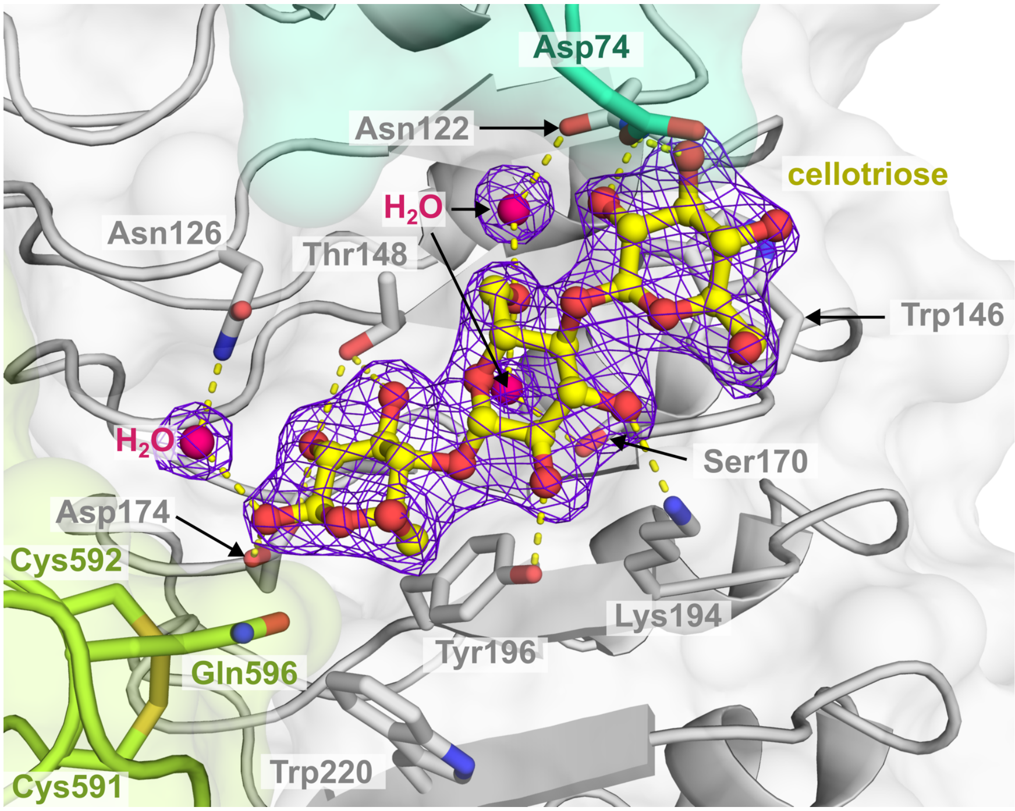


**Fig. S2. Cellotriose omit map in IGP1 binding pocket.** The electron-density omit map (|Fo|−|Fc|), contoured at 3σ (purple mesh), highlights the cellotriose ligand (yellow sticks) and crystallographic water molecules (hot pink) involved in the hydrogen-bond network (yellow dashed lines). Residues Cys591, Cys592, and Gln596 located at the malectin domain, shape the pocket without forming direct hydrogen bonds with the carbohydrate ligand.

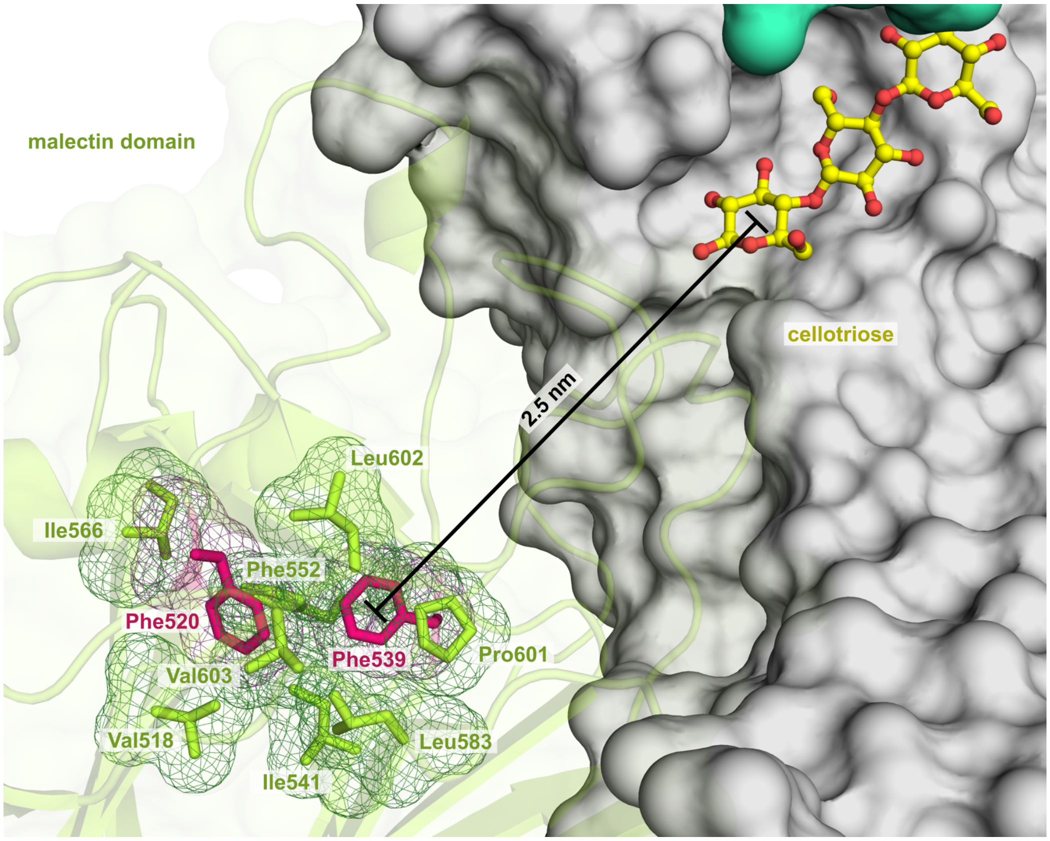


**Fig. S3 Phe residues 520 and 539 are essential for maintaining the structural integrity of the malectin domain in IGP1.** The core of this domain is stabilized by a network of hydrophobic residues (represented as green sticks), with the bulky Phe520 and Phe529 (highlighted in magenta) playing a key role. These residues are buried within the hydrophobic core, shielded from solvent exposure, and positioned distally from the cellotriose ligand-binding pocket (25 Å), which resides within the LRR domain (depicted as a gray surface) of the receptor.

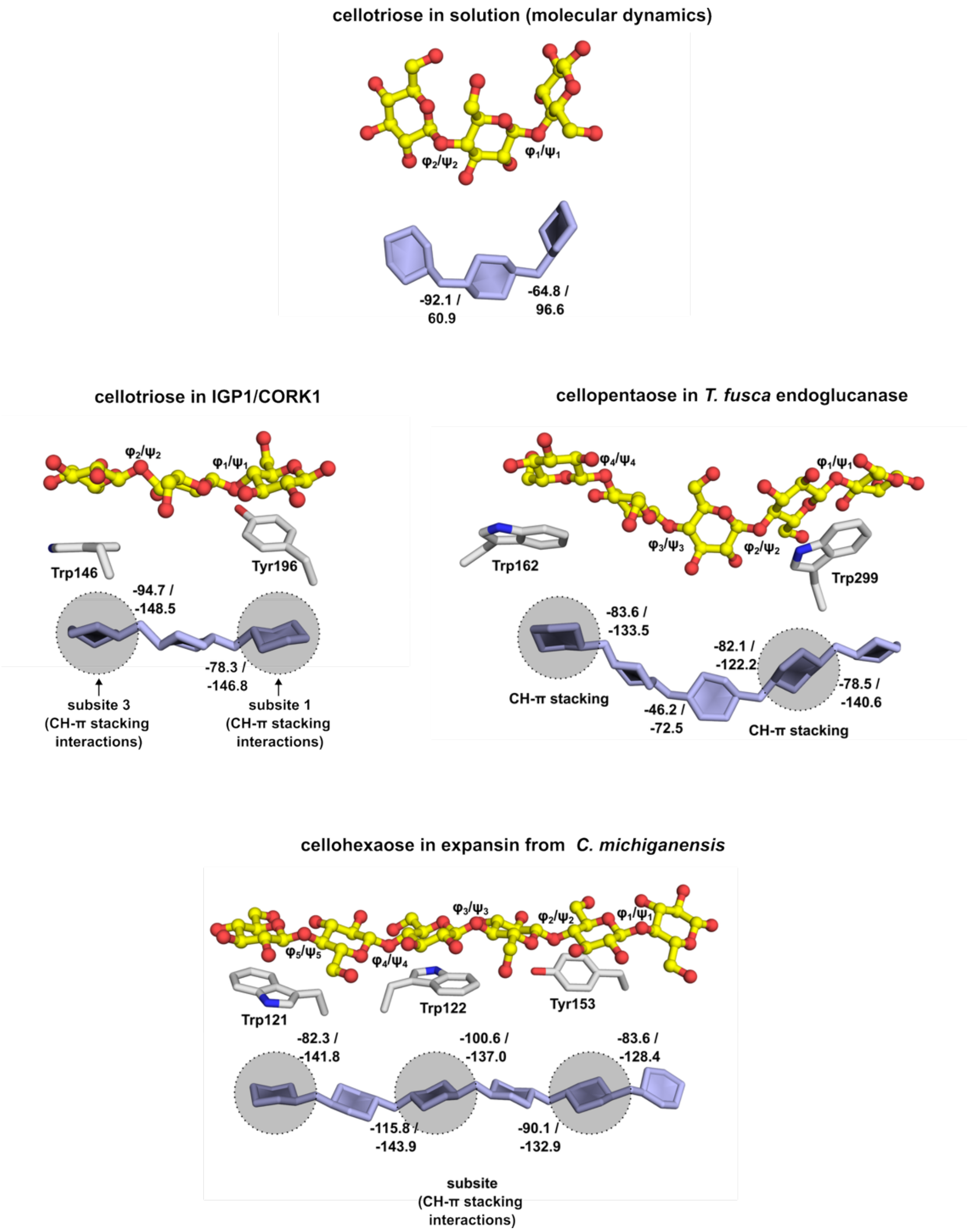


**Fig. S4 CH-π stacking interactions anchor and shape flexible cello-oligomers.** Comparison of cello-oligomer conformations in various protein binding pockets. Top: cellotriose in solution, modeled using molecular dynamics (ATB ID: 31965) (*99*) , showing high conformational flexibility of glucopyranose rings defined by dihedral (torsion) angles Phi (φ) and Psi (ψ). Rings are shown in light blue. Middle left: cellotriose bound to IGP1 adopts a planar conformation stabilized by CH-π interactions with Tyr196$^{IGP1}$ and Trp146$^{IGP1}$ (this study). Middle right: cellopentaose in *T. fusca* endoglucanase (PDB ID: 2CKR) adopts a twisted conformation at the active site, shaped by CH-π interactions. Bottom: cellohexaose on *C. michiganensis* expansin (PDB ID: 4L48) adopts a flat conformation, mediated by CH-π interactions. CH-π interactions mediated by aromatic residues are highlighted by grey circles.

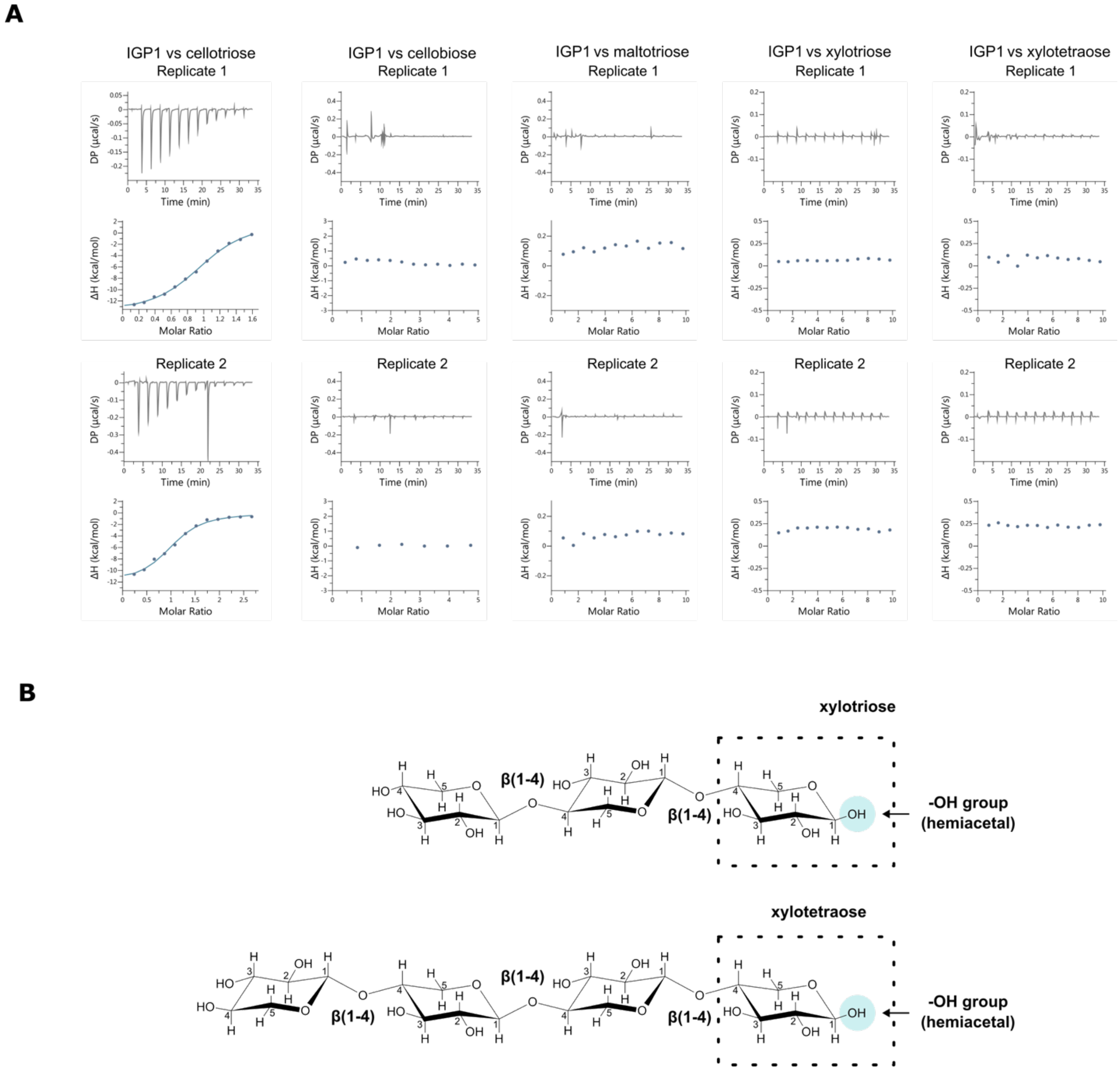


**Fig. S5 Isothermal titration calorimetry (ITC) assays of IGP1 with various cello-oligomers, stereoisomers, and xylans, as summarized in Fig. 2H.** (A) ITC thermograms from independent experiments analyzed in Fig. 2H. (B) 2D representation of oligomers of β-xylose (β-D-xylopyranose). The hydroxyl group (-OH) of the hemiacetal is highlighted with a blue circle and the reducing end is enclosed within the dashed rectangle.

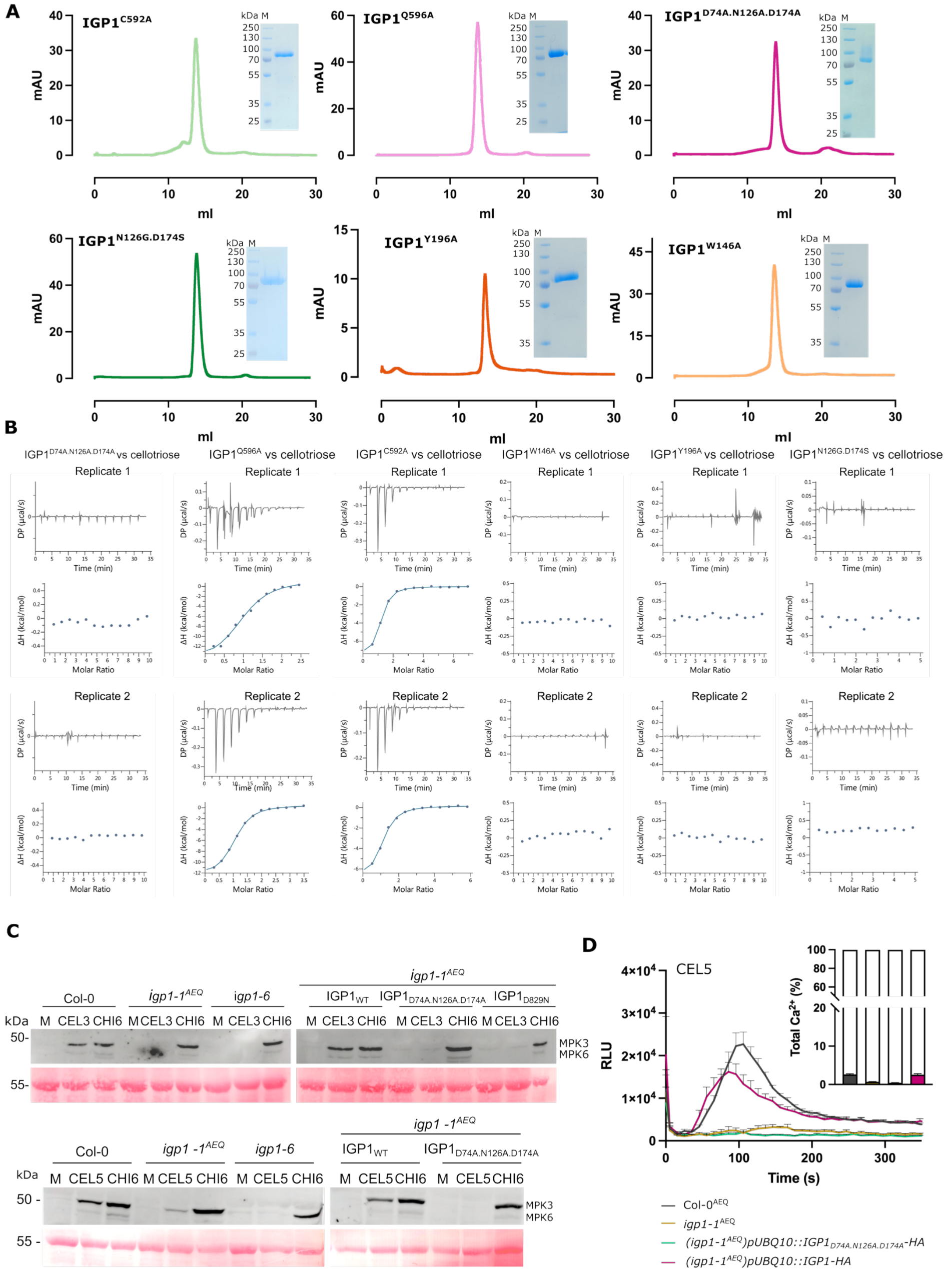


**Fig. S6 IGP1 specifically binds cello-oligomers and activates signaling responses**. (A) Size-exclusion chromatography profiles and SDS–PAGE of extracellular domain (ECD) variants of IGP1 expressed in insect cells. (B) ITC thermograms of independent replicates indicated in Fig. 2J. (C) MAPK activation in response to cellotriose (CEL3) cellopentaose (CEL5) in *igp1-1*$^{AEQ}$ allele background. Seedlings of Col-0, *igp1-1*$^{AEQ}$, *igp1-6*, and complemented lines expressing *pUBQ10::IGP1-HA* or the mutant variant *pUBQ10::IGP1*$_{D74A.N126A.D174A}$*-HA* were treated with 10 µM CEL3, CEL5, 50 µM CHI6, or water (mock). Phosphorylated MAPKs were detected 15 min after treatment using anti-pTEpY (anti-p42/44) antibodies; Ponceau staining was used as a loading control. (D) Cytosolic $Ca^{2+}$ burst in response to CEL5. Luminescence traces (RLU) over time are shown for Col-0$^{AEQ}$, *igp1-1*$^{AEQ}$, and complemented lines after 10 µM CEL5 treatment. Data show mean ± SEM. (n = 8) from one of three independent experiments with similar results. Total $Ca^{2+}$ was discharged by the addition of 1mM $CaCl_2$ to the wells and these values were used for the calculation of the total $Ca^{2+}$ % induced by CEL5.

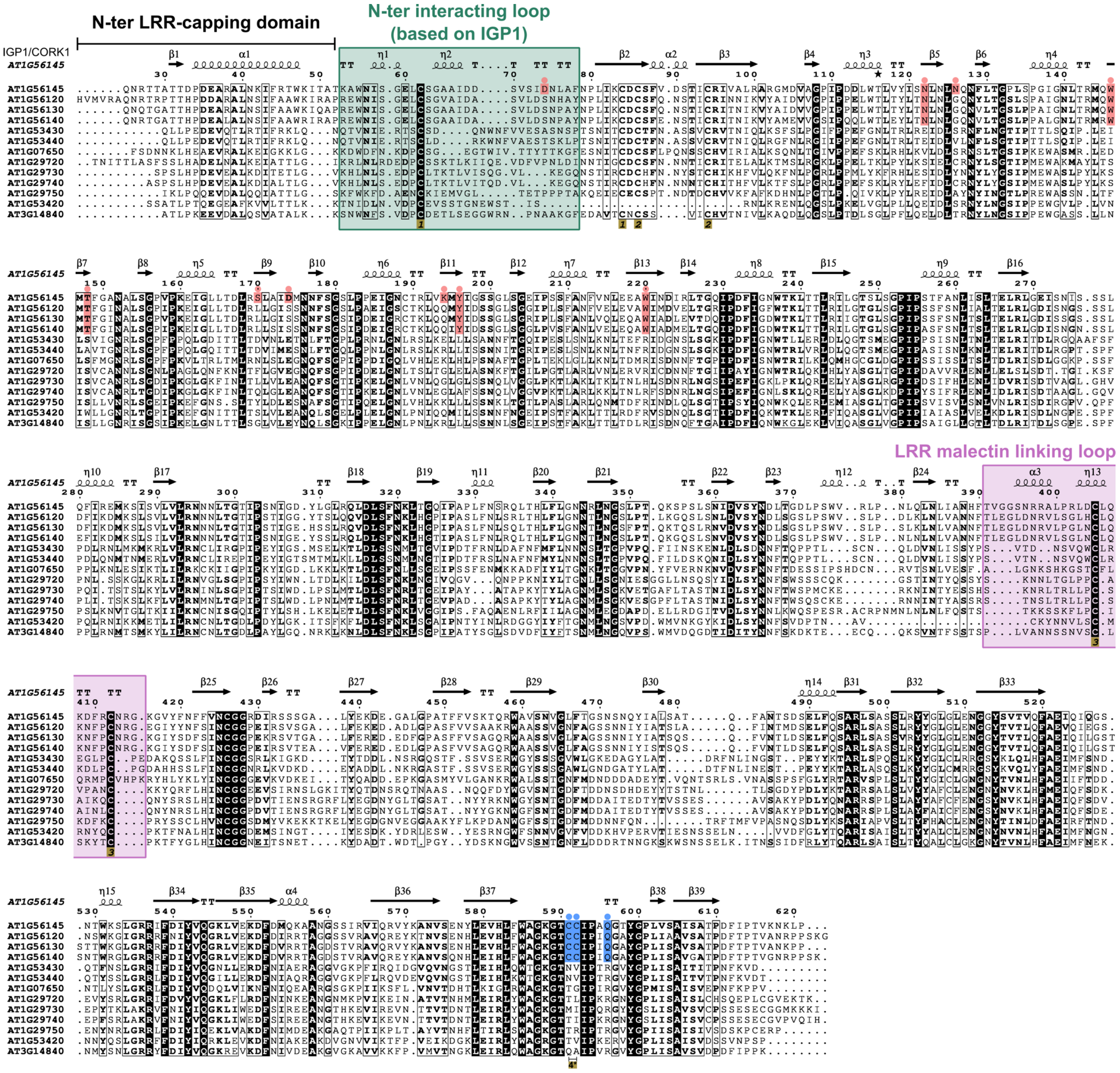


**Fig. S7 Conservation of structural features for precise carbohydrate recognition and shaping in IGP1 across the IGP1-like receptor subfamily**. Ectodomain sequence alignment of the LRR-malectin receptor family in *A. thaliana*, including secondary structure information and residue numbering based on the IGP1-cellotriose crystal structure (PDB: 9HHX, this study). Ligand-binding residues coordinating cellotriose are highlighted in pink salmon, with the same coloring applied to conserved residues in other IGP1-like receptors. The conserved residues in the malectin domain that shapes the pocket are highlighted in blue. Three conserved sulfur bridges, indicated with numbers, are present across the LRR-malectin family, with one unique to IGP1-like members (highlighted in Fig. 2B). Cys-Cys pairs are numbered in gold yellow at the bottom of the alignment.

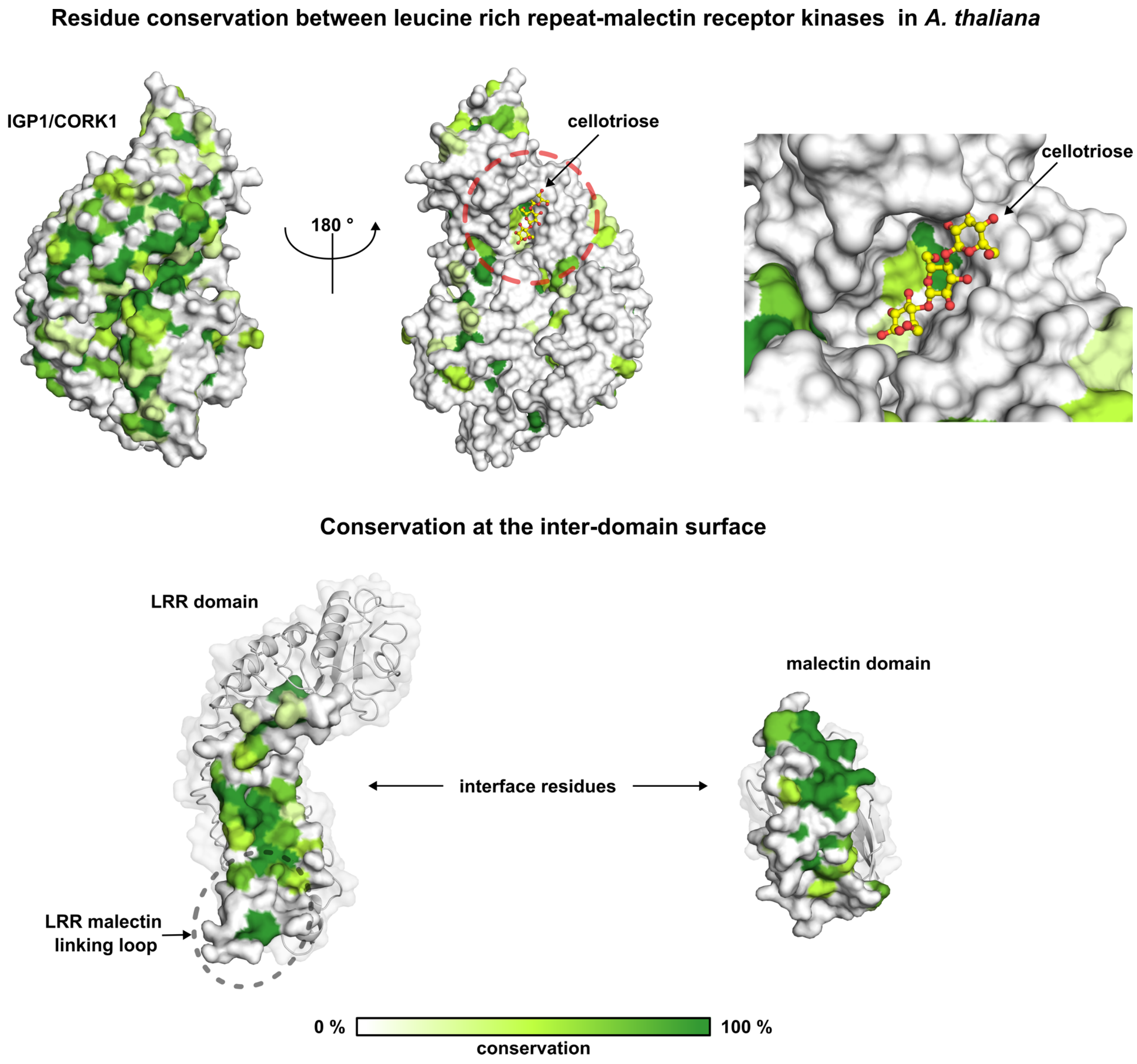


**Fig. S8 Conservation analysis reveals a highly diverse ligand-binding pocket within the Arabidopsis LRR-malectin receptor superfamily, while the inter-domain surface remains notably conserved.** Top: surface representation and side views of the IGP1 ectodomain, colored by residue conservation. Right: close-up view of the IGP1 ligand pocket in complex with cellotriose, showing low conservation within the LRR-malectin family binding-site. This suggests the potential recognition of chemically diverse ligands. Bottom: close-up view of the inter-domain surface is highly conserved, indicating its relevant role in protein folding and interdomain arrangement.

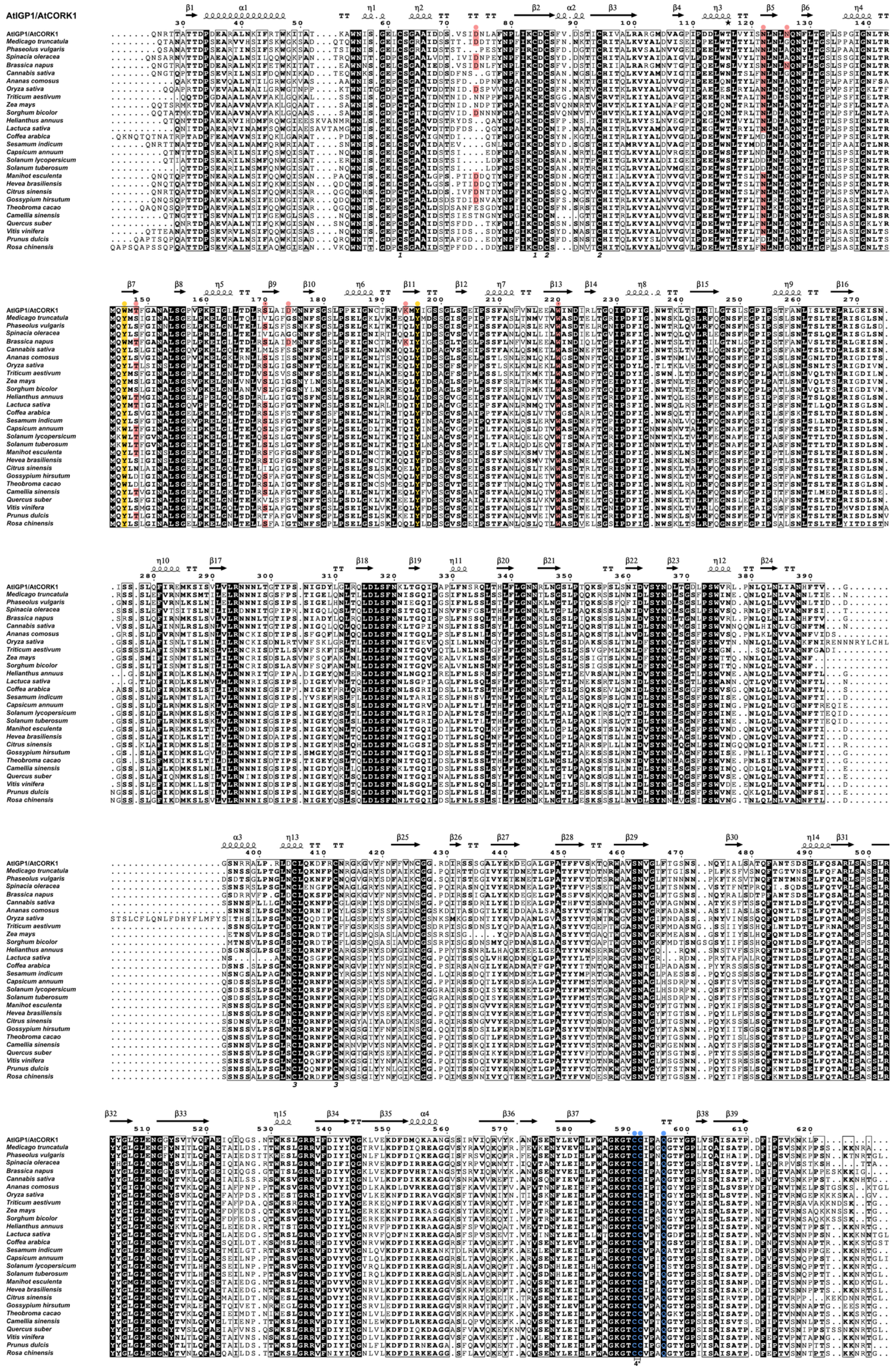

**Fig. S9 Sequence alignment of the ectodomain of *At*IGP1 and IGP1-like receptors from different plant species**. Protein sequences correspond to the IGP1-like orthologs identified in Fig. S10. Conserved aromatic residues essential for CH-π stacking interactions with β-linked sugars are highlighted in yellow. Conserved residues involved in hydrogen bonding between *At*IGP1 and cellotriose are shown in pink salmon, with the same coloring applied to corresponding residues in IGP1-like orthologs. The conserved residues in the malectin domain that shapes the pocket are highlighted in blue. Cys-Cys pairs are numbered at the bottom of the alignment.

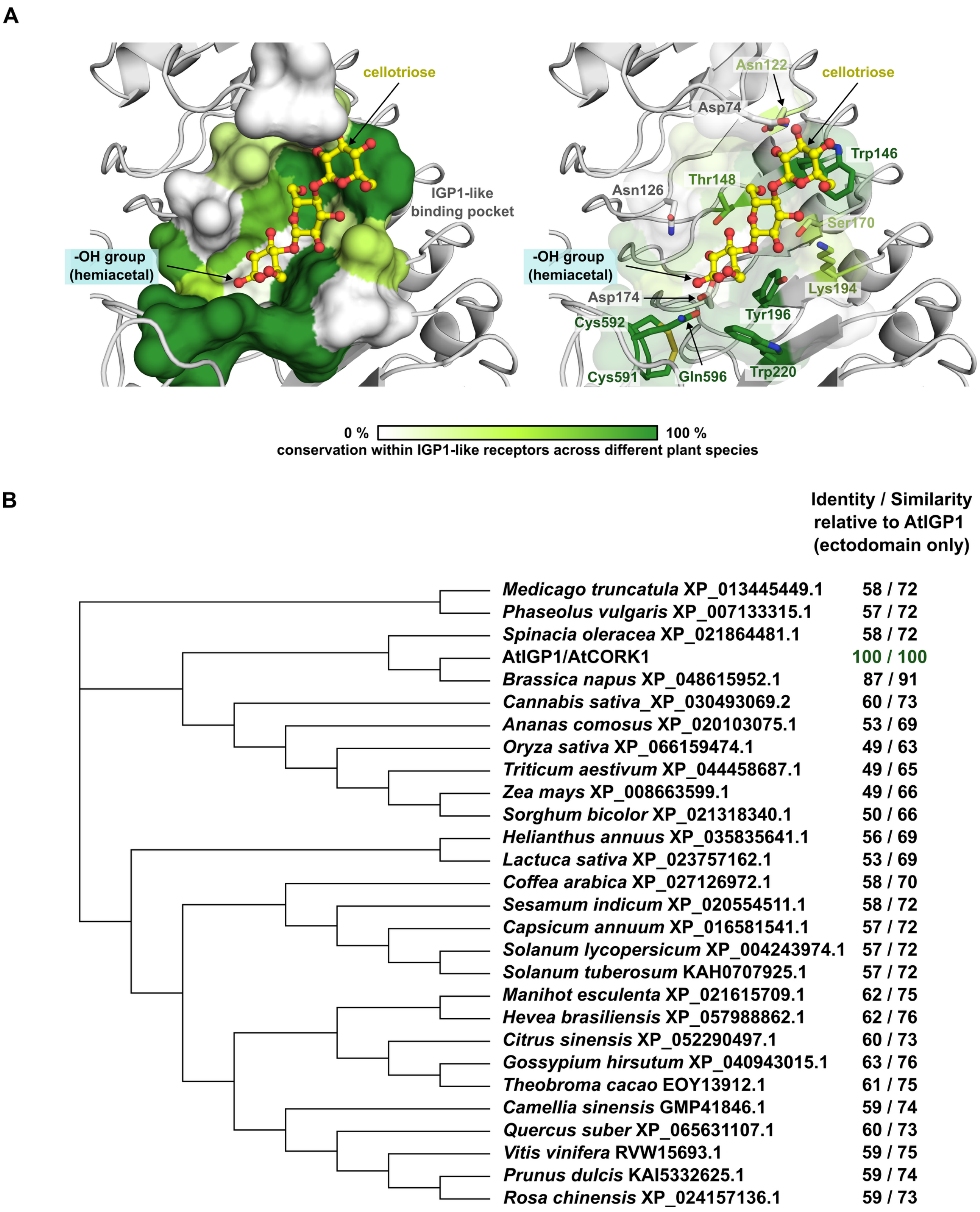


**Fig. S10 Conservation of the IGP1-like receptor subfamily across plant species.** (A) Conservation analysis of the sugar-binding pocket in *A. thaliana* IGP1 and its orthologs. Aromatic residues coordinating β-pyranose rings at subsites 1 and 3 are invariant, suggesting a conserved mechanism for recognizing and binding oligosaccharides with β-glycosidic linkages. (B) Phylogenetic analysis of IGP1-like orthologs, with identity scores relative to *At*IGP1 indicated.

**A**

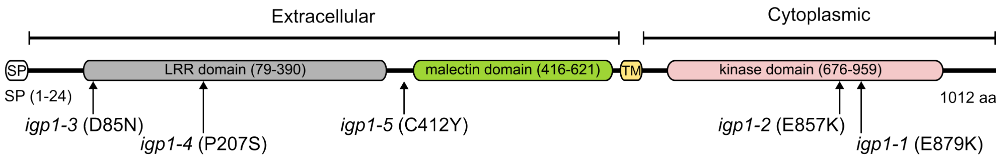


| Allele | gene_id | Position | Change | Col-0$^{AEQ}$ pool_AD1 | allele pool_AD2 | allele pool_FREQ | funct_consequ | Splicing variant 1 | Splicing variant 2 |
|---|---|---|---|---|---|---|---|---|---|
| ***igp1-2*** | AT1G56145 | 21008767 | C to T | 221 | 162 | 1 | nonsynonymous SNV | exon23:c.G2569A:p.**E857K** | exon22:c.G2650A:p.E884K |
| ***igp1-3*** | AT1G56145 | 21013481 | C to T | 149 | 116 | 0.97 | nonsynonymous SNV | exon2:c.G253A:p.**D85N** | exon2:c.G253A:p.D85N |
| ***igp1-4*** | AT1G56145 | 21012628 | G to A | 190 | 22 | 0.85 | nonsynonymous SNV | exon7:c.C619T:p.**P207S** | exon7:c.C619T:p.P207S |
| ***igp1-5*** | AT1G56145 | 21010951 | C to T | 172 | 23 | 1 | nonsynonymous SNV | exon16:c.G1235A:p.**C412Y** | exon16:c.G1235A:p.C412Y |

**B**

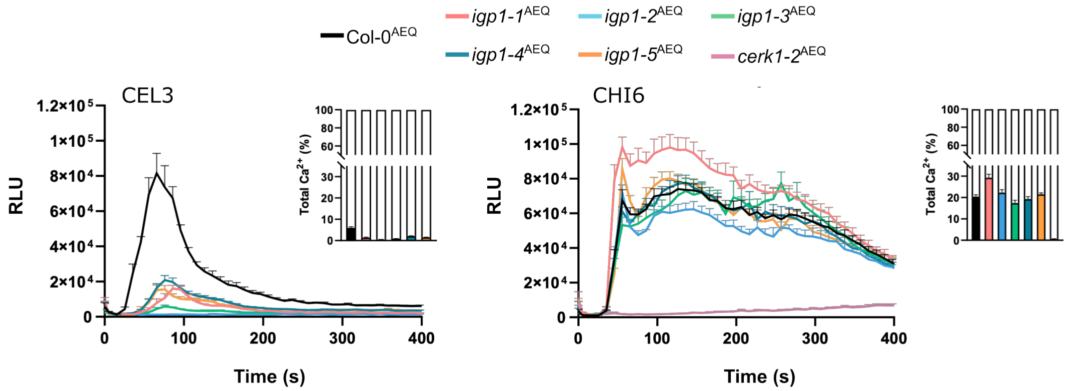


**C**

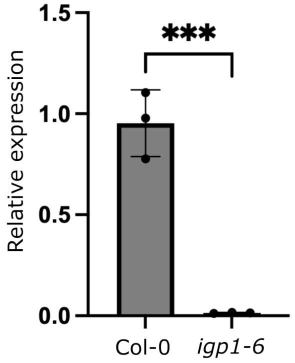


**Fig. S11 Mutations in the extracellular and kinase domain of *Arabidopsis thaliana* IGP1 impair $Ca^{2+}$ responses to cellotriose.** (A) Schematic representation of the IGP1/CORK1 receptor topology. The leucine-rich repeat (LRR) domain is shown in grey, the extracellular malectin-like domain in green, the transmembrane (TM) domain in yellow, and the intracellular kinase domain in pink. Point mutations in IGP1 are indicated with arrows: *igp1-1$^{AEQ}$* (*17*), and *igp1-2$^{AEQ}$* to *igp1-5$^{AEQ}$* (this study). A table summarizes the mutations, including chromosome position, reference and alternate alleles, SNP frequency, and predicted amino acid changes. (B) $Ca^{2+}$ influx measured in response to 10 µM cellotriose (CEL3) or 50 µM chitohexaose (CHI6) in wild-type (Col-0$^{AEQ}$), *igp1$^{AEQ}$* mutant alleles, and the *cerk1-2$^{AEQ}$* line (defective in CHI6 perception), expressed as relative luminescence units (RLU) over time. Data represent means ± SEM (n = 8) from one of four independent experiments with similar results. Total $Ca^{2+}$ discharge was induced by 1 mM $CaCl_2$ addition and used to calculate the percentage of $Ca^{2+}$ release triggered by cellotriose or chitohexaose treatments (see insets and calculation details in (*17*). (C) IGP1 expression is disrupted in the T-DNA knockout line *igp1-6* (SALK_101924). Expression relative to SAND (At2g28390). Bars represent the average of three biological replicates, *** p-value < 0.005 (T-test).

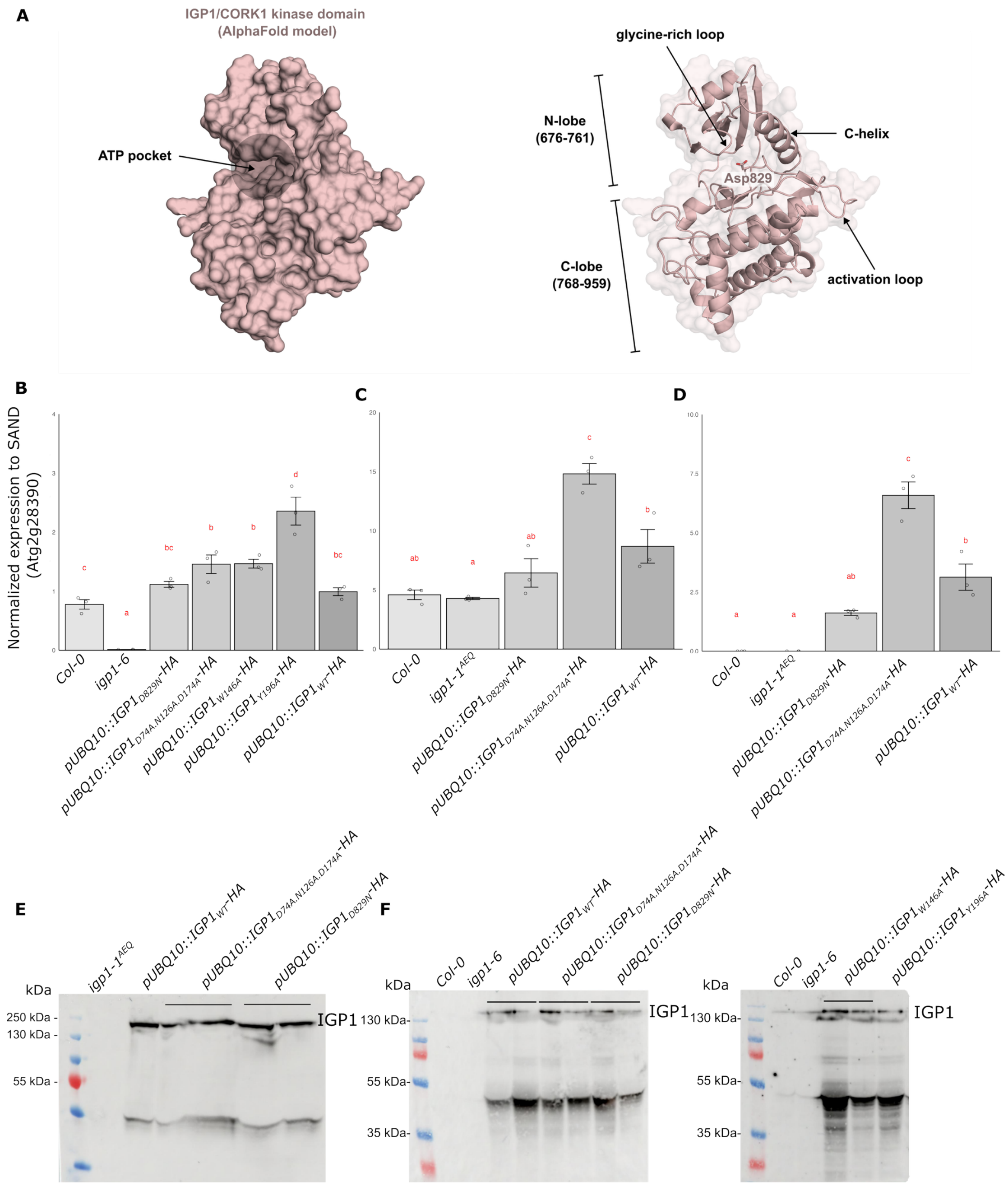


**Fig. S12 Complementation lines with the IGP1 splicing variant 1 kinase domain retain all essential elements of a functional receptor kinase.** (A) Left: Surface representation of the IGP1 kinase domain (splicing variant 1) based on an AlphaFold prediction (model ID: C0LGH4). The ATP-binding pocket is highlighted by a grey circle. Right: Cartoon representation of the N-lobe and C-lobe regions, with key structural and regulatory elements indicated by arrows. The conserved metal-coordinating residue Asp829 (DFG motif) is shown as sticks at the ATP-binding pocket entrance. Mutation of Asp829 to asparagine inactivates the kinase, disrupting IGP1 downstream signaling (Fig. 3C). (B) qRT-PCR analysis of expression levels of IGP1 wild-type and variants in *igp1-6* knockout background transgenic lines used in this study (primers listed in Table S2). (C–D) qRT-PCR expression analysis of IGP1wild-type and variants in *igp1-1$^{AEQ}$* EMS background (*17*) transgenic lines used in this study. (D) Transgene-specific expression assessed using HA tag–discriminating primers. Bars represent the average of 3 technical replicates. Letters correspond to different statistical groups (ANOVA followed by Tukey test, p-values < 0.05). (E-F) Western blot analysis of IGP1 wild-type and variant proteins expressed in complementation lines in the *igp1-1$^{AEQ}$* and *igp1-6* backgrounds generated in this study.

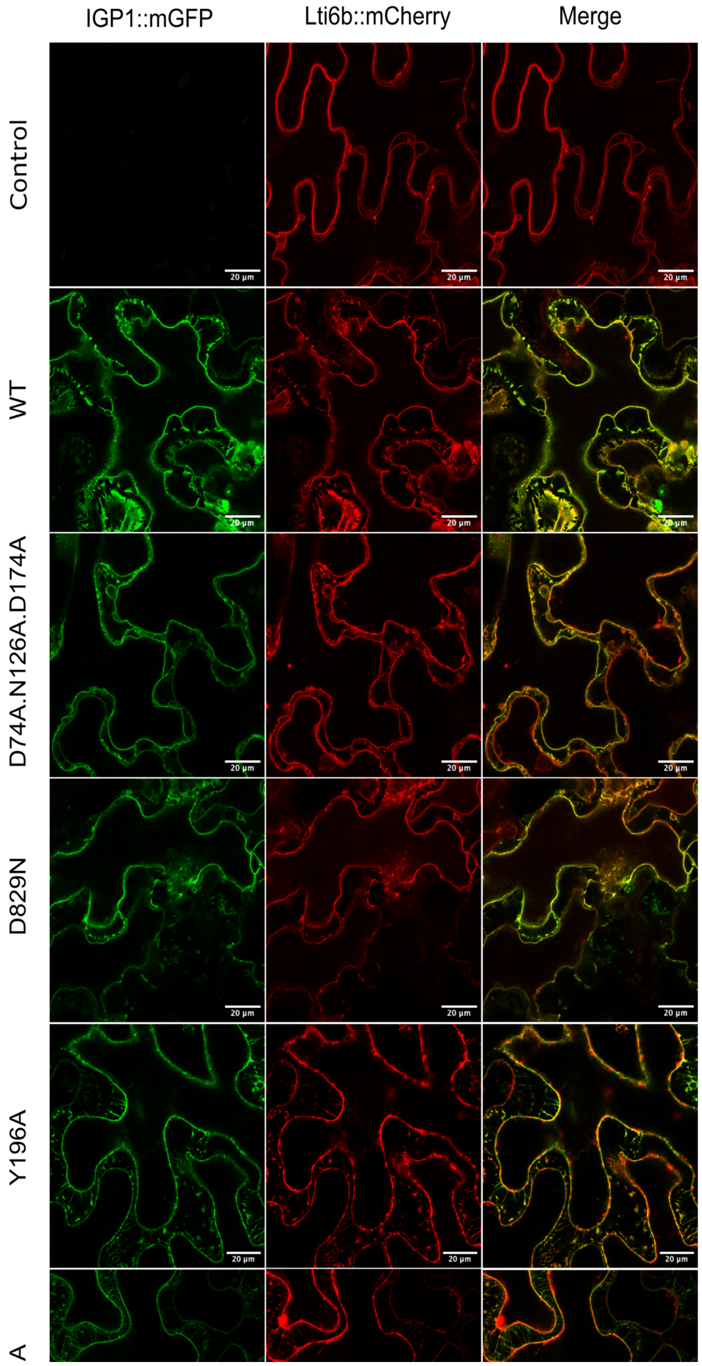


**Fig. S13 *In planta* plasma membrane localization of IGP1 wild-type and mutant variants.** (A) Subcellular localization of *pUBQ10::IGP1-mGFP* wild-type and mutants was analyzed via transient expression and plasmolysis assays in *N. benthamiana* leaves. The *p35S::Lti6b-mCherry* plasma membrane marker was used to visualize the plasma membrane during plasmolysis. The first row shows control samples, including the plasma membrane marker alone and non-infiltrated samples. Scale bar = 20 µm.

A

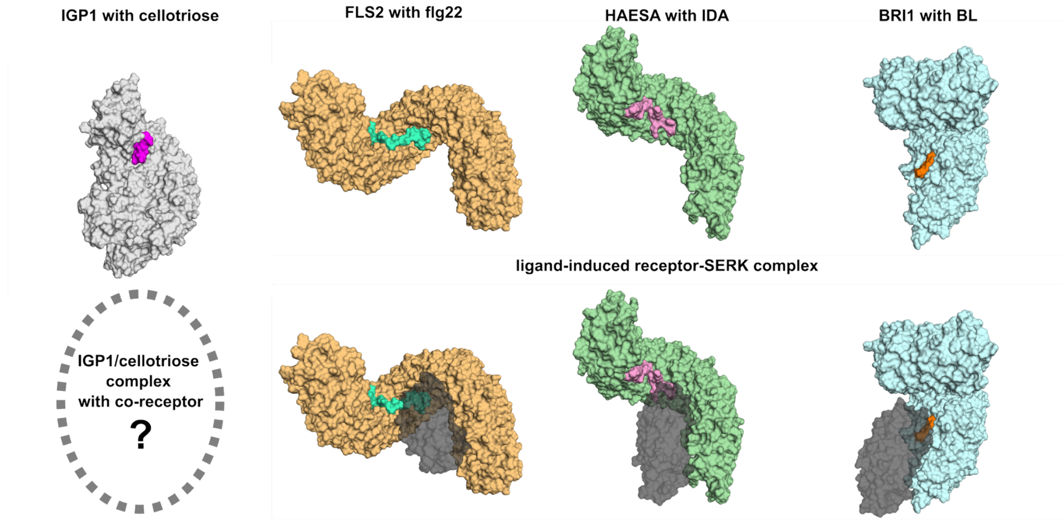


B

| | IGP1/cellotriose | FLS2/flg22 | HAESA/IDA | BRI1/BL |
|---|---|---|---|---|
| **ligand SASA in protein pocket, Å$^2$** | 195.4 | 1456.4 | 891.5 | 145.5 |
| **ligand buried SASA by co-receptor, Å$^2$** | unknown | 166.1 | 216.5 | 118.9 |

C D

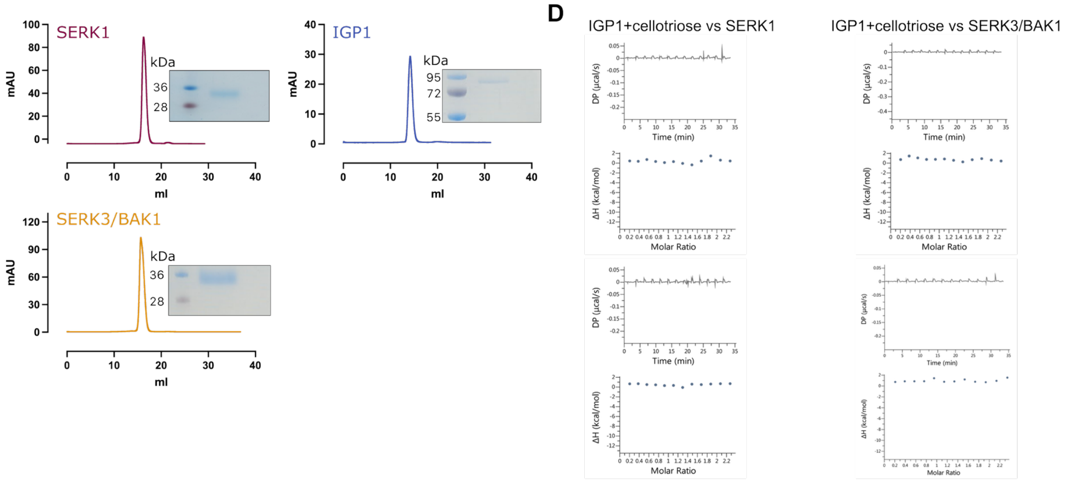


E

| Ligand (syringe) | Protein (cell) | $K_d$ (μM) | ΔH (kcal/mol) | N |
|---|---|---|---|---|
| SERK1 | IGP1 + Cellotriose | n.d | n.d | |
| SERK3 (BAK1) | IGP1 + Cellotriose | n.d | n.d | |

**Fig. S14 Cellotriose binding to IGP1 creates a potential interaction surface for a co-receptor.** (A) Surface representation of IGP1 bound to cellotriose (magenta) (this study PDB: 9HHX) compared to plant plasma membrane receptors FLS2-flg22 (PDB ID: 4MN), HAESA-IDA (PDB ID: 5IYX), and BRI1-BLD (PDB ID: 4LSX) and their ligand-induced complexes with SERK co-receptors (dark grey, bottom). (B**)** Calculated Solvent Accessible Surface Area (SASA) of ligands, showing cellotriose's occluded area is comparable to SERK-bound complexes. (C, D) The SERK membrane-kinase family are not co-receptors of IGP1-cellotriose. Size-exclusion chromatography (SEC) profiles and SDS-PAGE of BAK1 (SERK3) and SERK1. (D) ITC binding experiments and summary table of SERK proteins versus IGP1 + cellotriose. n.d indicates non-detected binding. (E) ITC summary table of SERK proteins versus IGP1 + cellotriose. n.d indicates non-detected binding.

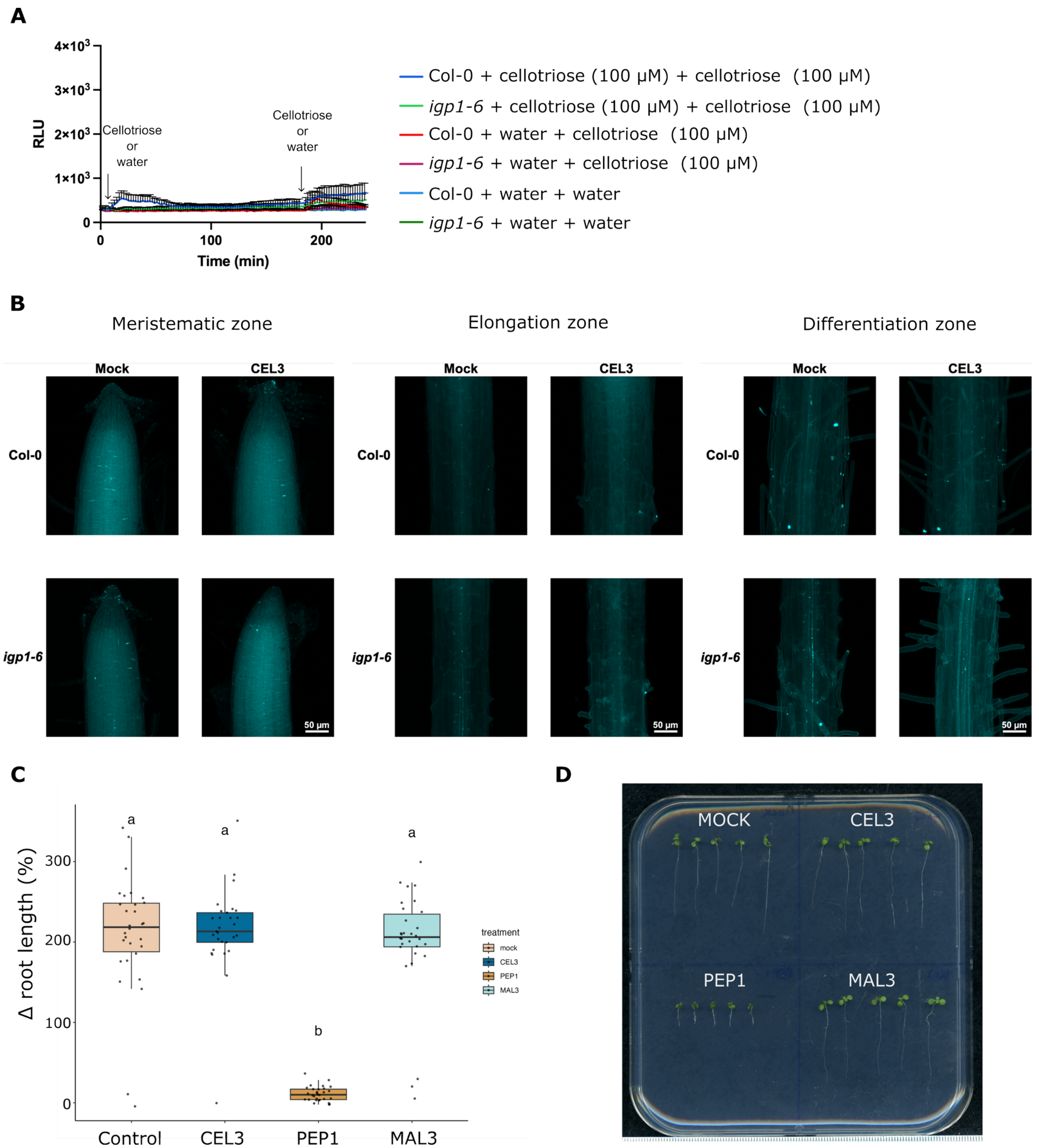


**Fig. S15 Cellotriose does not elicit canonical MAMP responses or growth inhibition.** (A) Cellotriose treatment (100 μM, 3 h) fails to induce a strong IGP1-dependent ROS response. ROS production was monitored over time in Col-0 and *igp1-6* seedlings ($n \geq 10$ per treatment, mean ± SEM., representative of $\geq 3$ independent experiments). (B) Cellotriose does not trigger IGP1-dependent callose deposition. Shown are representative aniline blue–stained roots 16 h post-treatment with 100 μM cellotriose (Col-0 vs. *igp1-6*). Scale bar, 50 μm ($n \geq 3$). (C) Cellotriose does not impair growth. Col-0 seedlings transferred 4 d after germination to plates containing mock ($H_2O$), cellotriose (100 μM), PEP1 (1 μM), or maltotriose (100 μM) were assayed for root elongation ($n \geq 29$). Relative growth was calculated after 3 d; statistical significance was assessed by Kruskal–Wallis test with Dunn post hoc (Bonferroni corrected). (D) Representative seedlings from the experiment in (C).

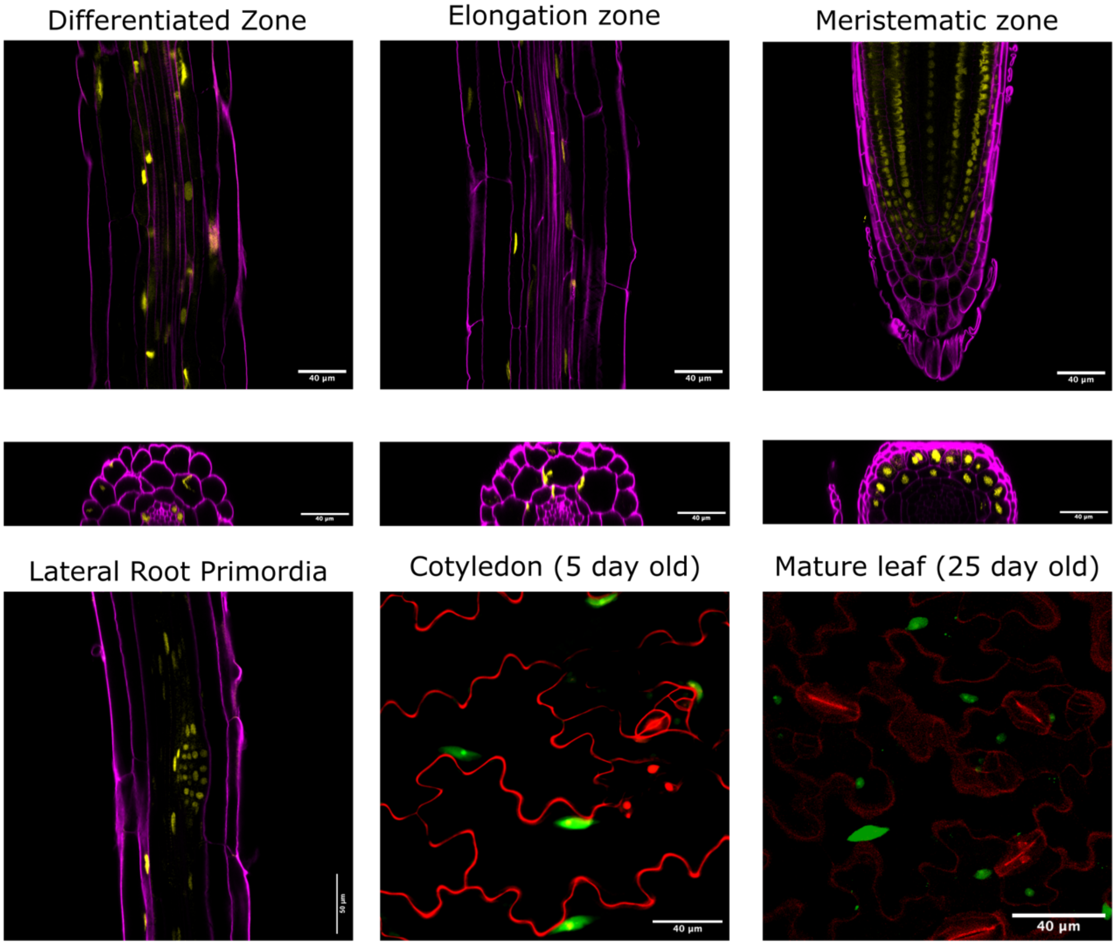


**Fig. S16 Expression pattern of IGP1 in seedling roots, cotyledons and adult leaves.** Representative confocal images of the *pIGP1::3xNLS-mGFP* transcriptional reporter line. Single-plane images of longitudinal (top) and orthogonal view of longitudinal root sections (bottom) display nuclear GFP signals (green) across all root cell layers. Propidium iodide (PI, red) staining outlines cell structures. Scale bars: 40 or 50 µm; (n > 3). Six 5-day old seedlings per line from 3 independent lines were imaged for IGP1 expression showing similar results.

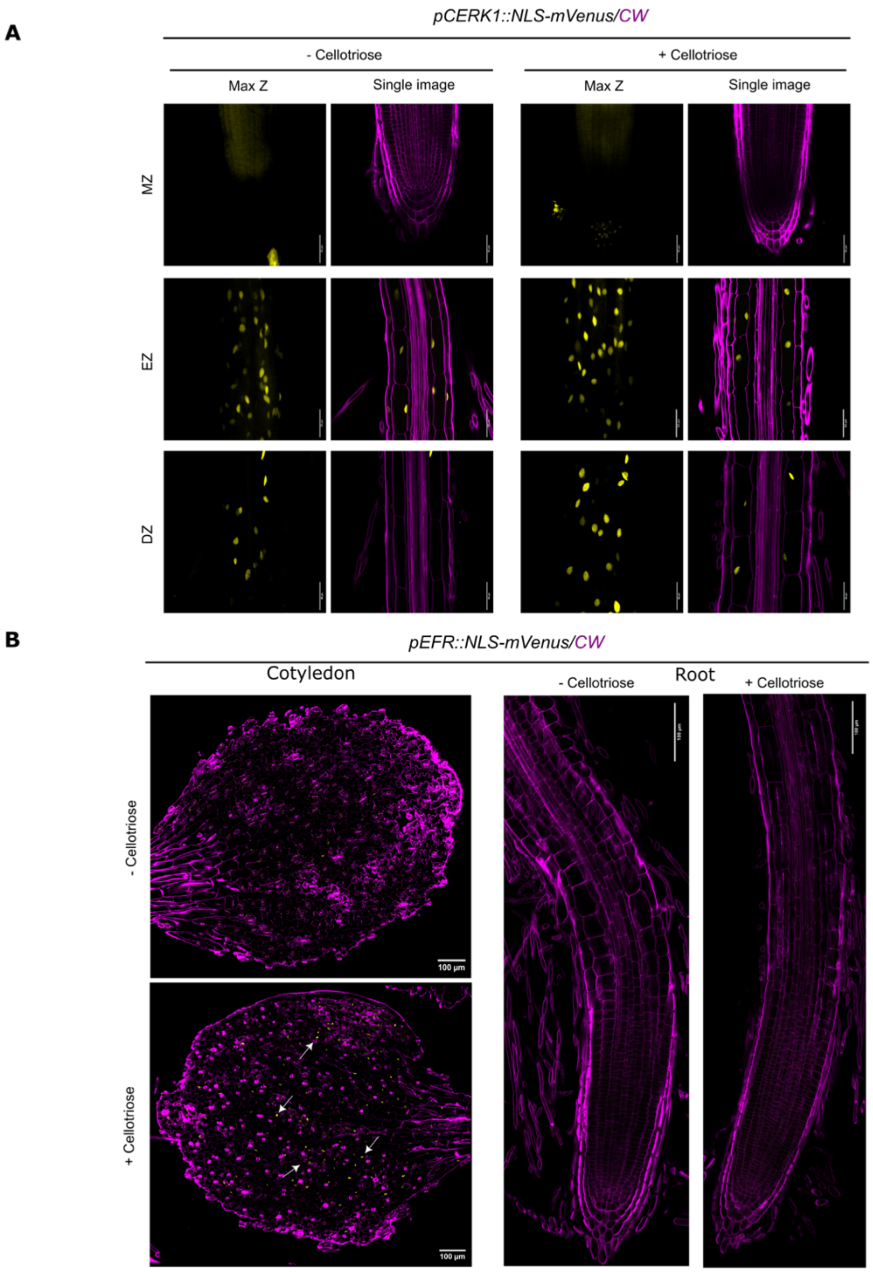


**Fig. S17 Cellotriose activates the expression of multiple pattern recognition receptors.**
(A) Representative expression pattern of the chitin receptor CERK1 upon 100 µM cellotriose treatment. Confocal images of 6-day-old *Arabidopsis* roots expressing the reporter line *pCERK1::NLS-3xmVENUS.* Three different zones were imaged: meristematic zone (MZ), elongation zone (EZ), and differentiation zone (DZ). Maximal projections of z stacks (max z, left) and single confocal sections (single image, right) are presented. Scale bar, 50 µm, n=3 seedlings were imaged. (B) Representative expression pattern of the pathogen-associated receptor EFR upon 100 µM cellotriose application. Single layer of cotyledons and roots were imaged. Scale bar = 100 um. Nuclear-localized *mVENUS* signals (yellow) are co-visualized with calcofluor (CW, purple).

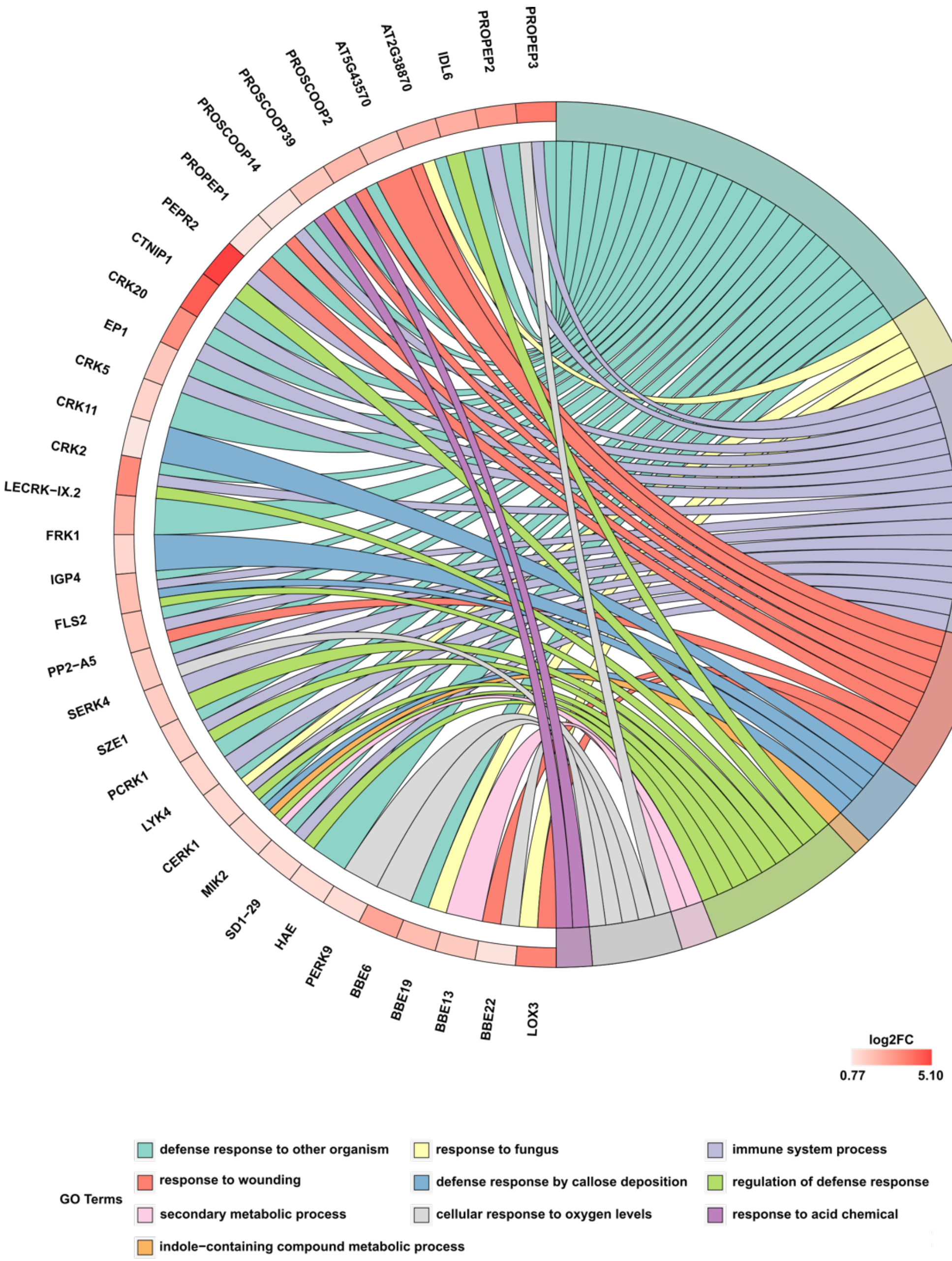


**Fig. S18 Functional analysis of a root RNA-seq dataset reveals activation of general immune responses following cellotriose treatment.** GOplot representation of immunity items enriched among differentially expressed genes from the publicly available dataset accession no. GSE198092 (Gene Expression Omnibus (GEO) database) (*36*). The analysis compares Arabidopsis seedling roots from Col-0 treated with 10 µM cellotriose vs water (mock treatment). Key upregulated categories include plasma membrane receptors, signaling peptides, and other molecular components involved in immune response modulation. Additionally, proteins involved in oligosaccharide redox homeostasis, such as members of the Berberine-like enzymes (BBE-like), are highlighted. The GOplot was generated using the GraphBio server (http://www.graphbio1.com/) (*100*).

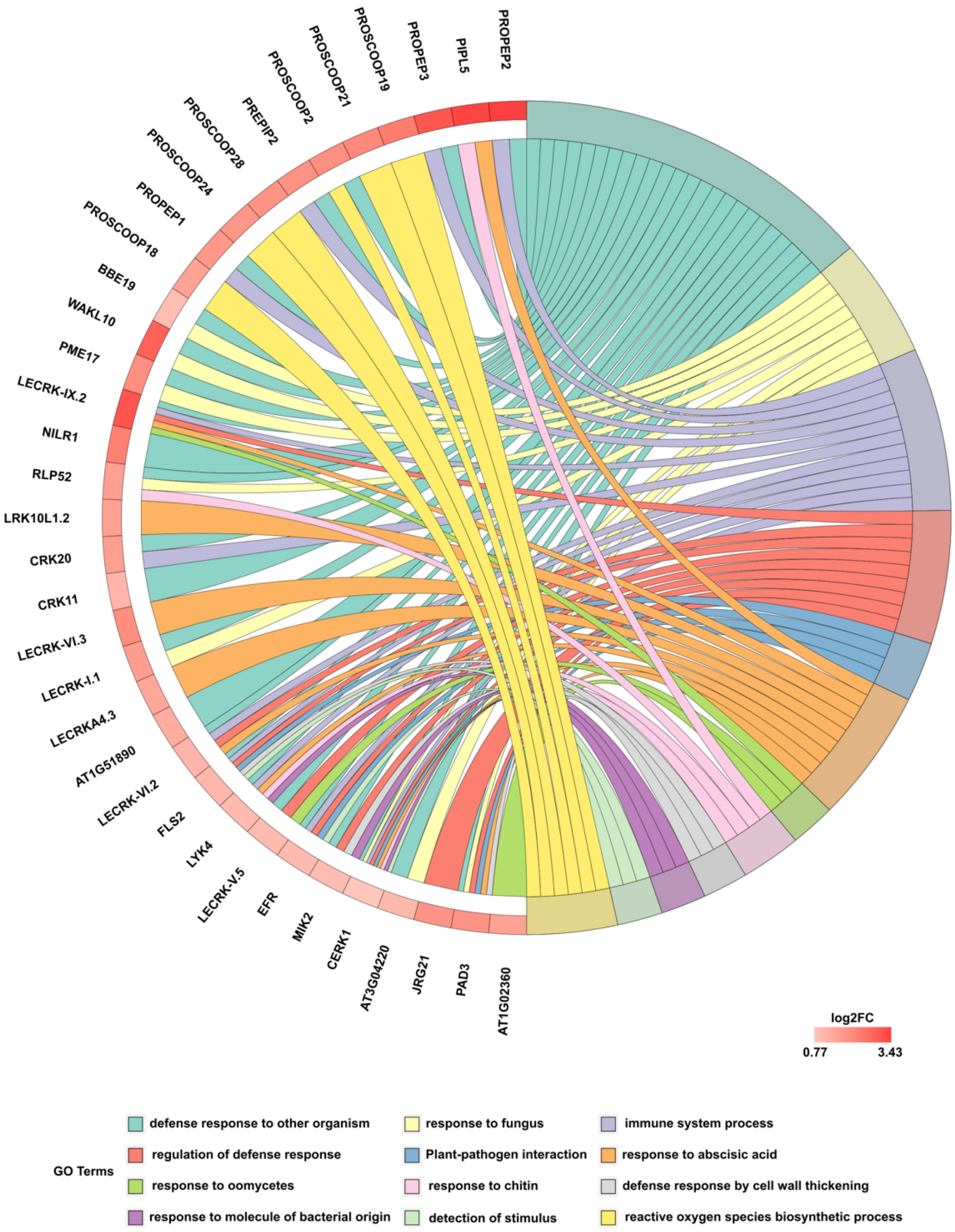


**Fig. S19 Functional analysis of full seedlings RNA-seq dataset shows the activation of general immune responses upon cellotriose treatment.** GOplot representation of immunity items enriched among differentially expressed genes from the publicly available dataset accession no. PRJNA1073490 (NCBI BioProject) (*14*). The analysis compares Arabidopsis seedlings treated with 10 µM cellotriose vs water (mock treatment). Key upregulated categories encompass plasma membrane receptors, signaling peptides, and various molecular components that play a crucial role in modulating immune responses. The GOplot was generated using the GraphBio server (http://www.graphbio1.com/) (***100***).

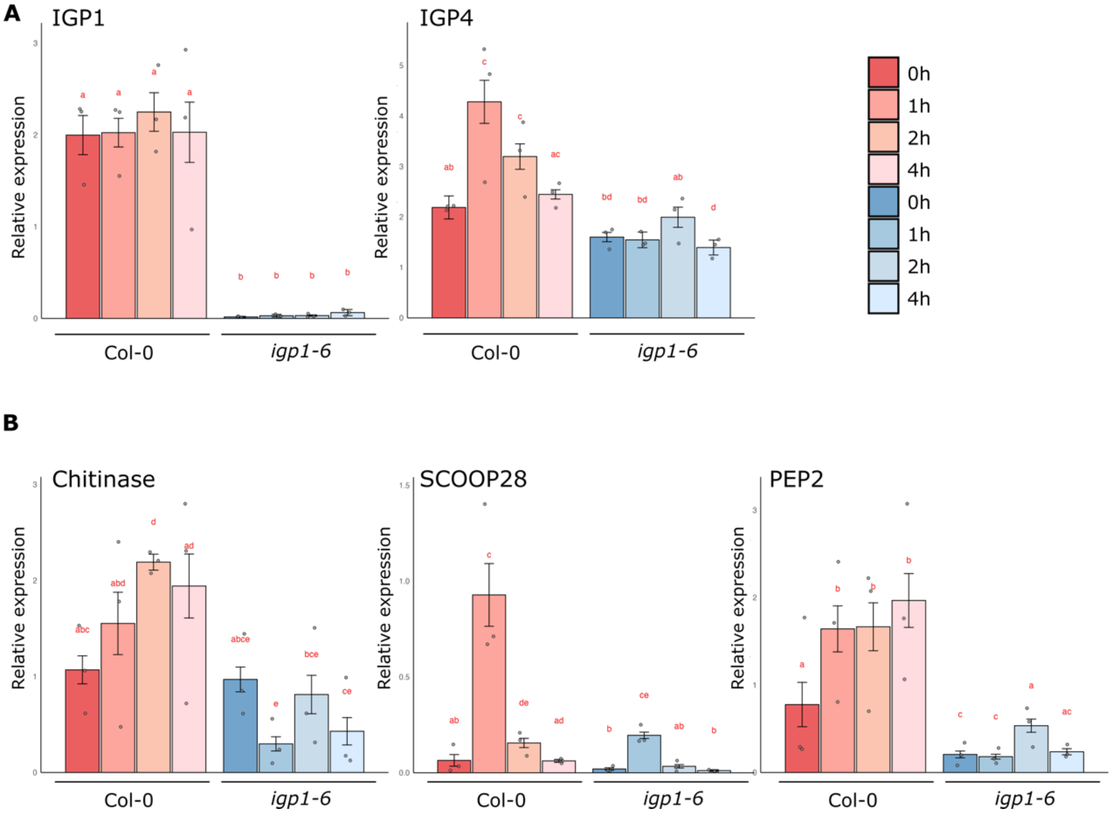


**Fig. S20 Cellotriose-induced expression of defense-associated genes.** (A) Cellotriose treatment strongly induces *IGP4* but not *IGP1*. (B) Broader transcriptional responses to cellotriose, show upregulation of defense-associated genes. Transcript levels were quantified by qRT–PCR in Col-0 and *igp1-6* seedlings treated with 100 µM cellotriose for 4 h and normalized to the reference gene SAND (At2G28390). Bars show mean ± SD from three biological replicates. Letters show different statistical groups; $P<0.05$ (aligned rank transform ANOVA. Post-hoc pairwise contrasts were adjusted using Tukey's method).

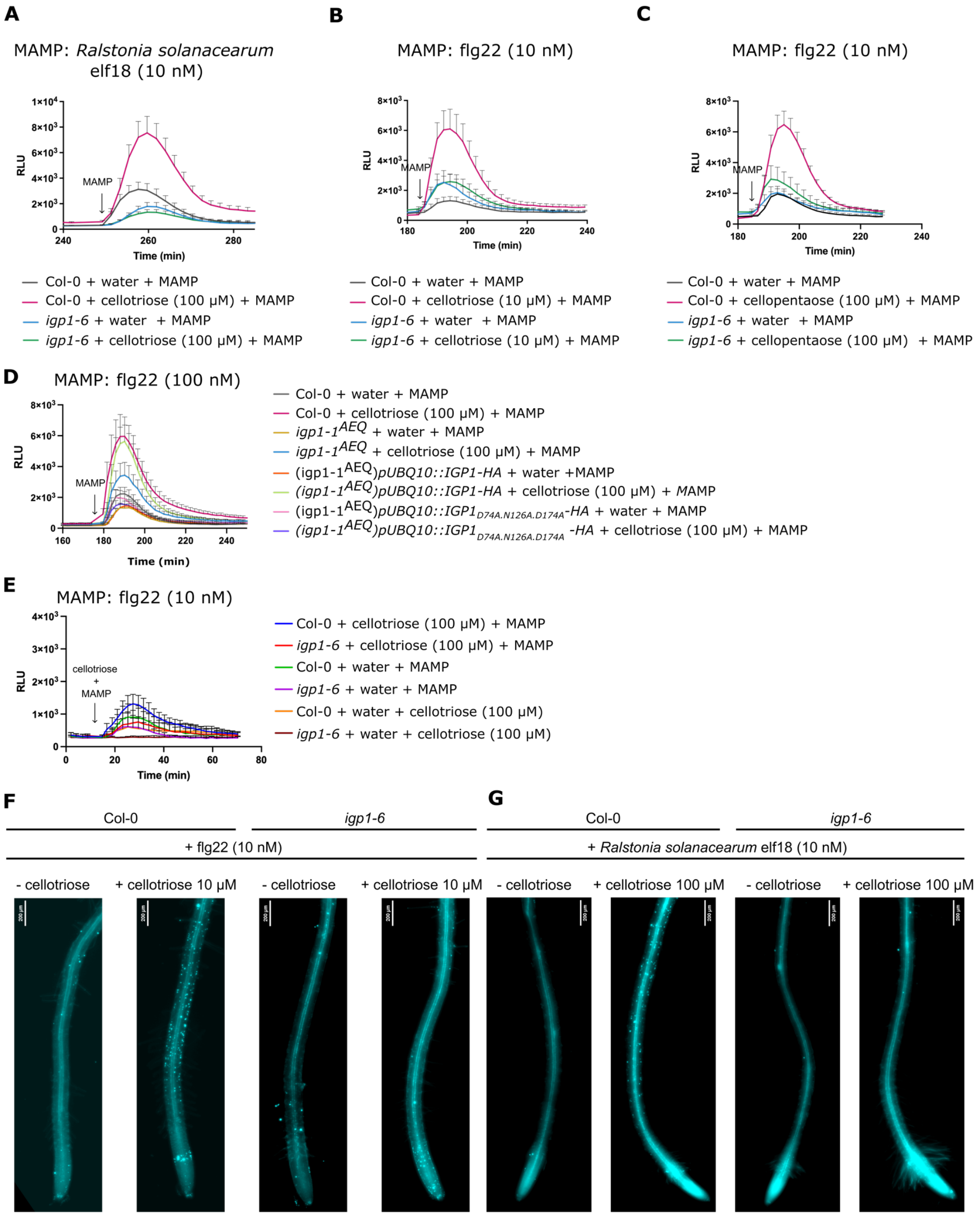


**Fig. S21 Cellotriose perception enhances Pattern-Triggered Immunity (PTI) responses.** (A-E) Pretreatment with cellotriose, sensed by IGP1, amplifies pathogen MAMP–induced ROS burst production in Arabidopsis. Seedlings of Col-0, *igp1-6*, and complemented lines in the *igp1-1*$^{AEQ}$ background (n ≥ 10 per treatment) were treated with mock solution (water) or cellotriose (10 or 100 μM) and challenged with MAMP elicitors 3–4 h later. No amplification was observed when cellotriose and the MAMP were applied simultaneously (E). ROS production was recorded as relative luminescence units; values represent the mean ± SEM of three technical replicates. Data are representative of at least three independent experiments. (F-G) Cellotriose pretreatment promotes callose deposition in roots following MAMP challenge in an IGP1-dependent manner. Shown are representative aniline blue–stained roots of Col-0 and *igp1-6* seedlings harvested 16 h after MAMP treatment from the same assay as in panels A–C. Scale bar, 200 μm (n ≥ 10 seedlings per condition).

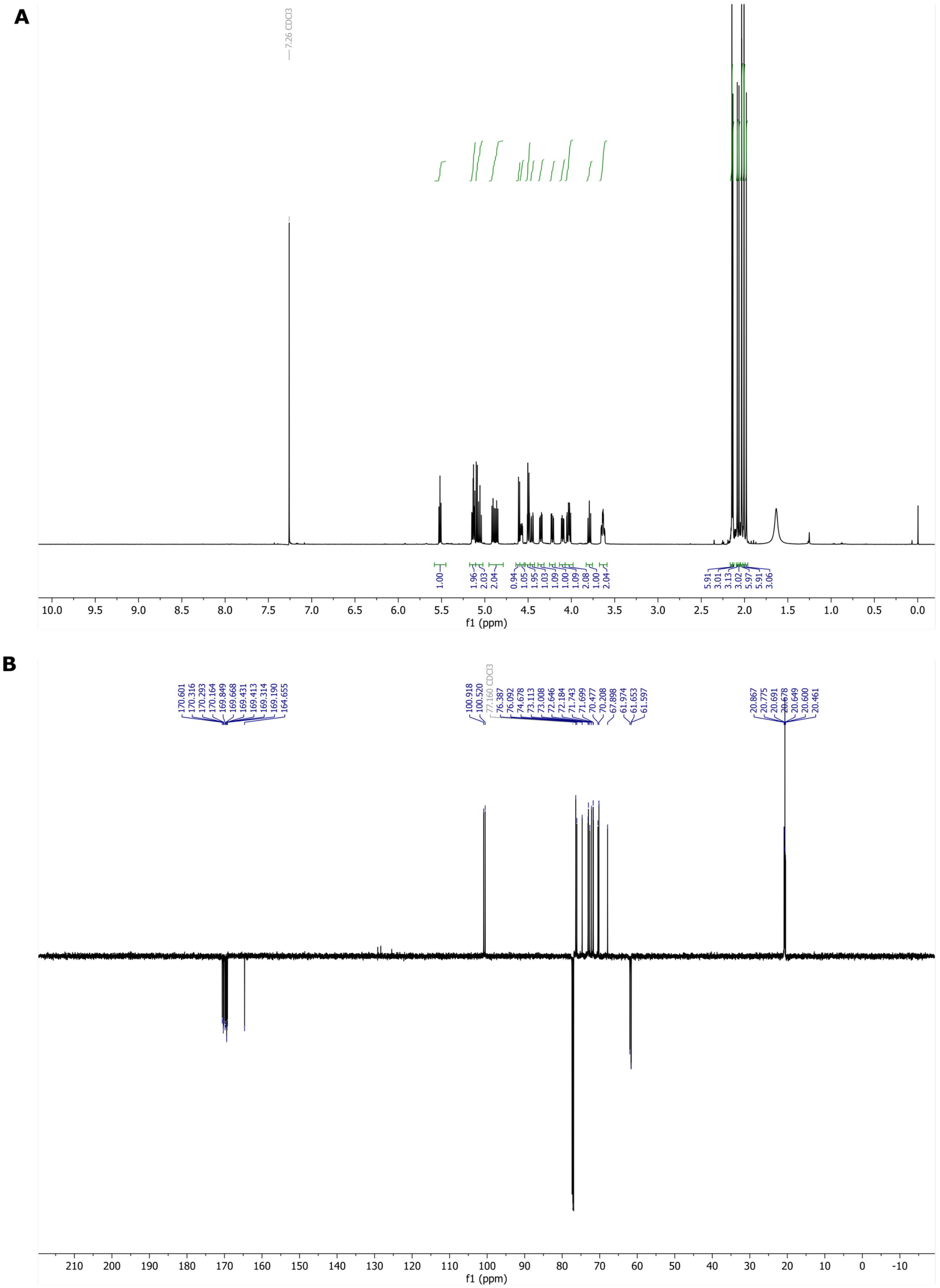


**Fig. S22 NMR spectra of intermediate compounds in the synthesis of cellotrionic acid 5 (CEL3ox).** (A) $^{1}$H NMR of lactone **3** in $CDCl_3$ (600 MHz, 297K). (B) $^{13}C$ APT NMR of lactone **3** in $CDCl_3$, (150 MHz, 297K).

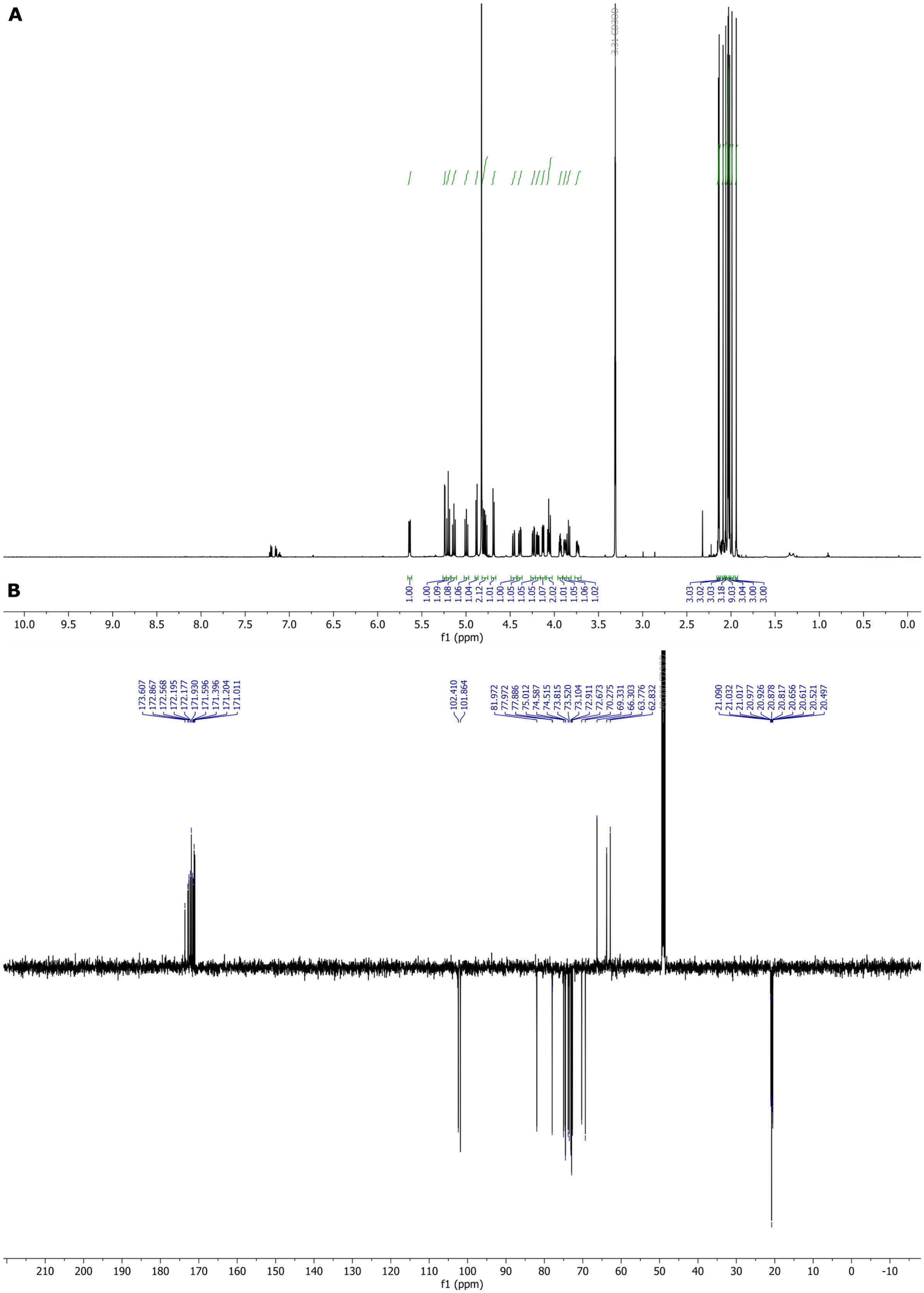

**Fig. S23 NMR spectra of intermediate compounds in the synthesis of cellotrionic acid 5 (CEL3ox).** (A) $^{1}$H NMR of acetylated sodium cellotrionate **4** ($CD_3OD$, 600 MHz, 297 K, 7.16 2.32 toluene). (B) $^{13}$C APT NMR of acetylated sodium salt **4** ($CD_3OD$, 150 MHz, 297 K).

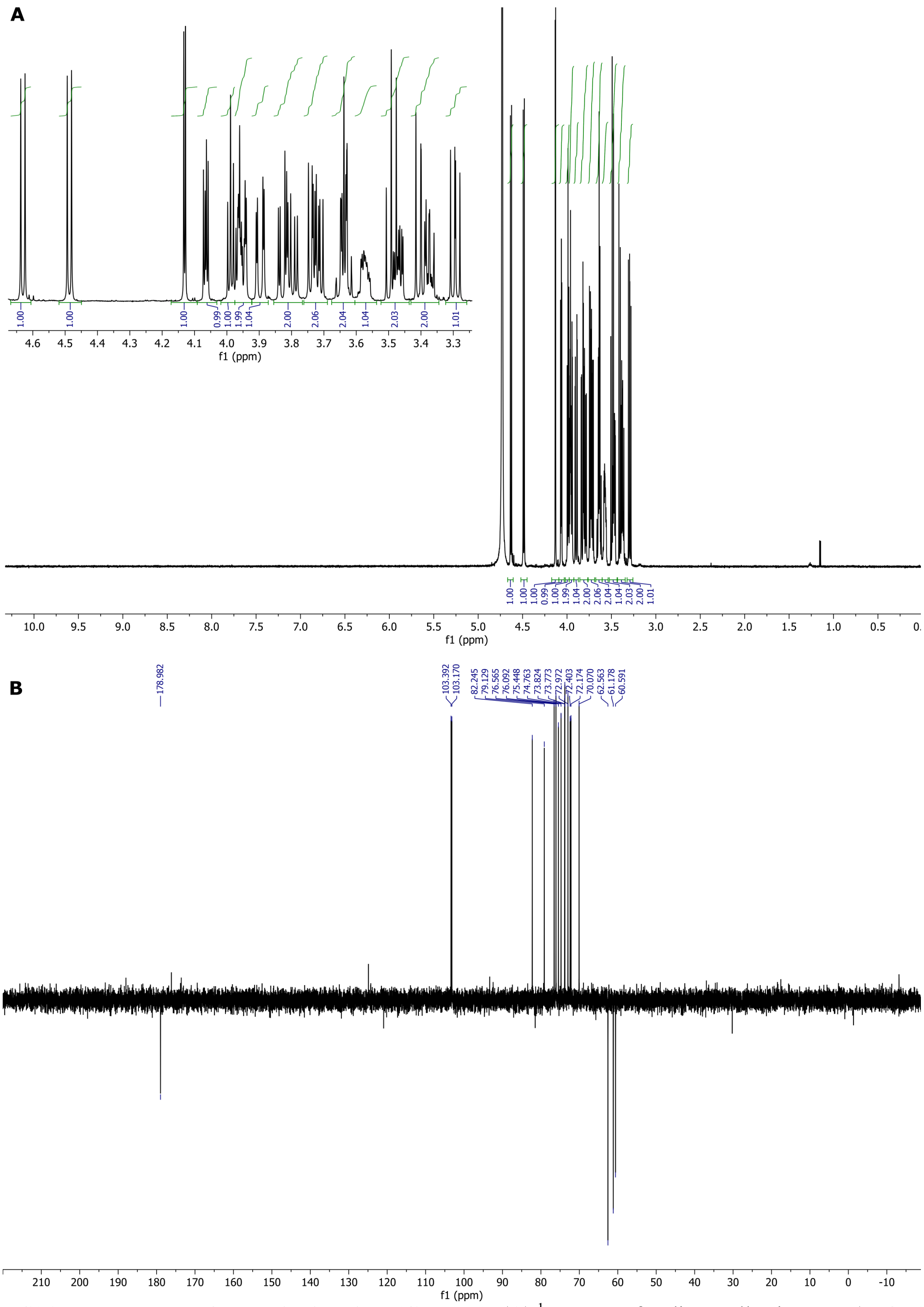


**Fig. S24 NMR spectra of cellotrionic acid 5 (CEL3ox).** (A) $^{1}H$ NMR of sodium cellotrionate **5** ($D_2O$, 600 MHz, 297 K)-. (B) $^{13}C$ APT NMR of sodium cellotrionate **5** ($D_2O$, 150 MHz, 297 K).

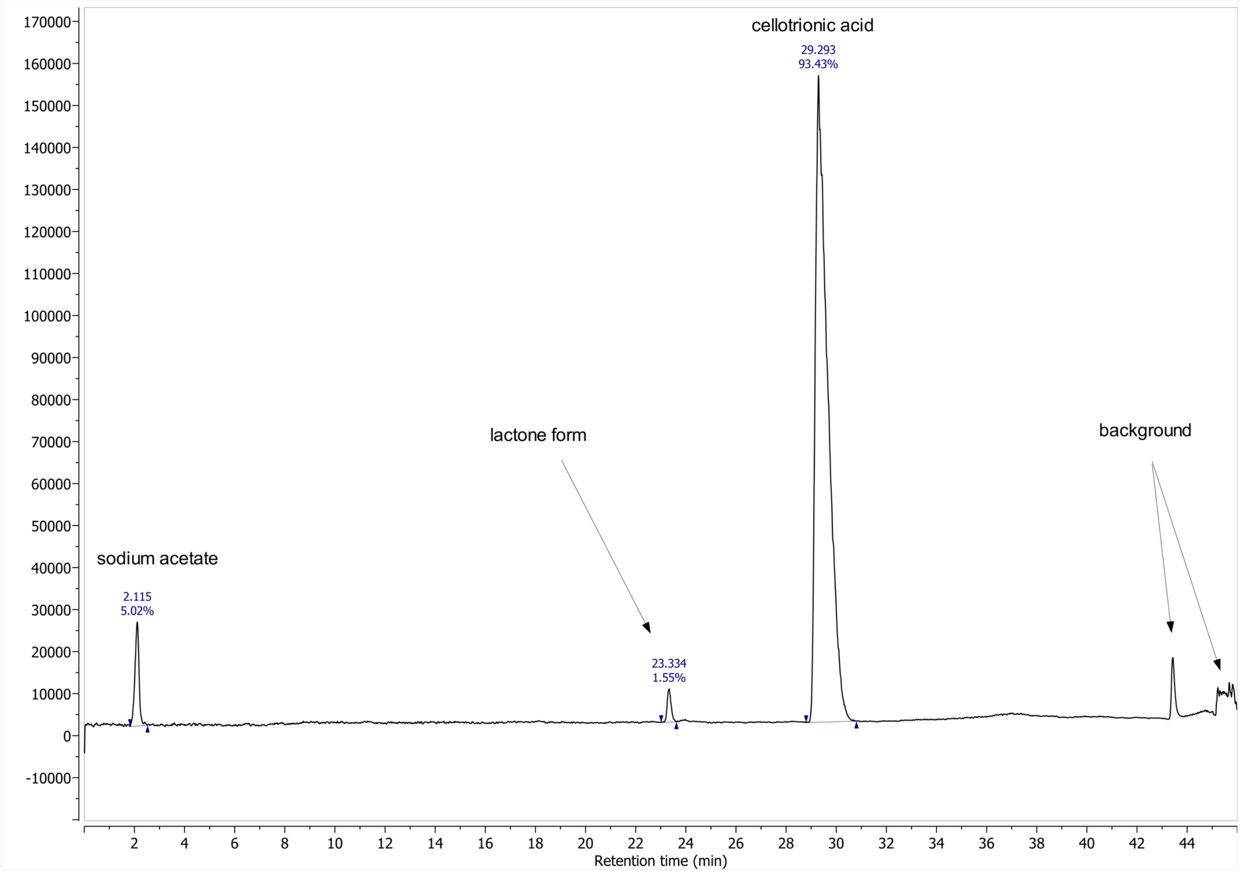


**Fig. S25 HPLC profile of sodium cellotrionate 5 (CEL3ox).** Gradient employed: 0-8 min 2.5% A, 8-33 minutes gradient to 40% A, 33-37 minutes gradient to 100% A, 0.7 mL/min. Solvent A: MeCN + 0.1% AcOH, Solvent B: $H_2O$ + 0.1% AcOH.

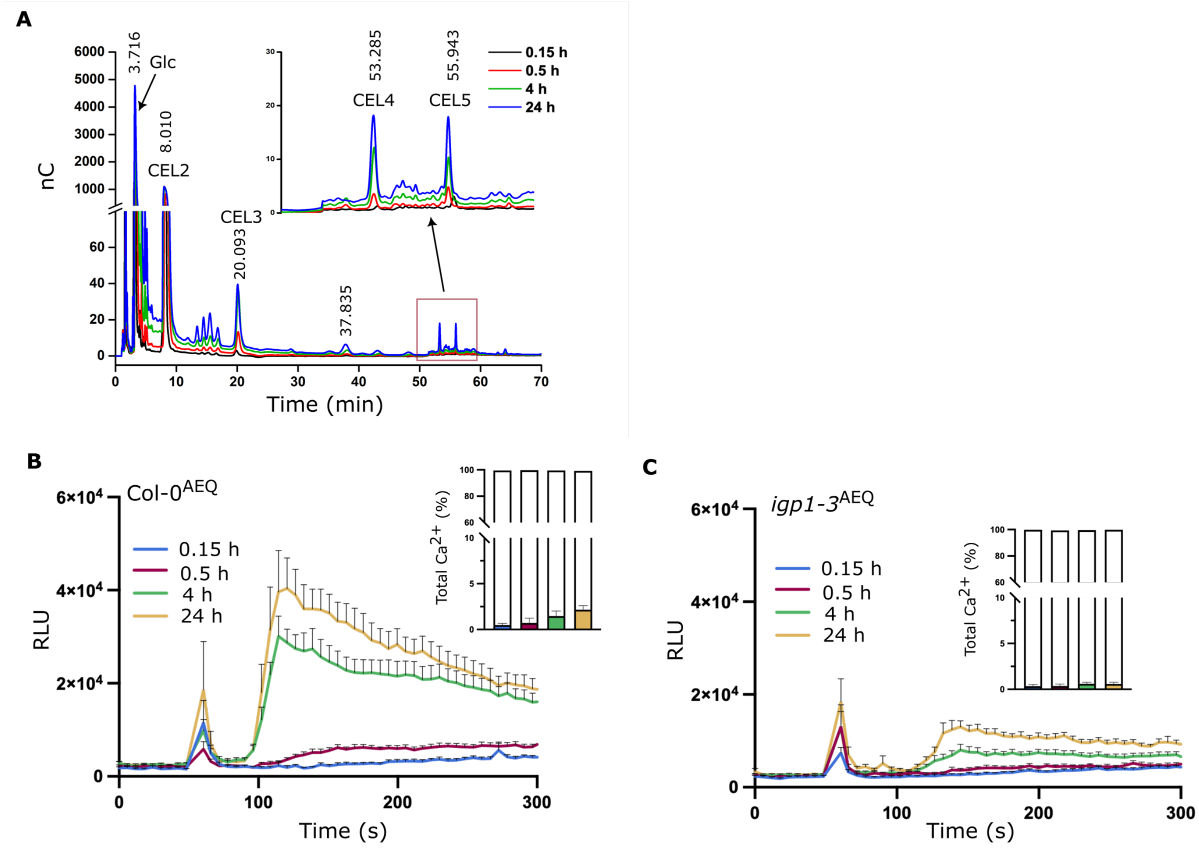


**Fig. S26 Cellulase-generated cello-oligomers do not elicit $Ca^{2+}$ signaling in *igp1-3* seedlings.** (A) HPAEC–PAD profiles of the oligosaccharide mixtures produced by cellulase digestion, with identified cello-oligomers indicated. Data are representative of four independent experiments with similar results. (B-C) Cytosolic $Ca^{2+}$ burst in Col-0$^{AEQ}$ and *igp1-3*$^{AEQ}$ seedlings Cytosolic $Ca^{2+}$ dynamics in Col-0$^{AEQ}$ and *igp1-3*$^{AEQ}$ seedlings treated with soluble fractions released from cellulose by cellulase digestion (0.1, 0.5, 4, and 24 h). $Ca^{2+}$ levels are shown as relative luminescence units (RLU) over time (mean ± SEM, n = 8).

A

B

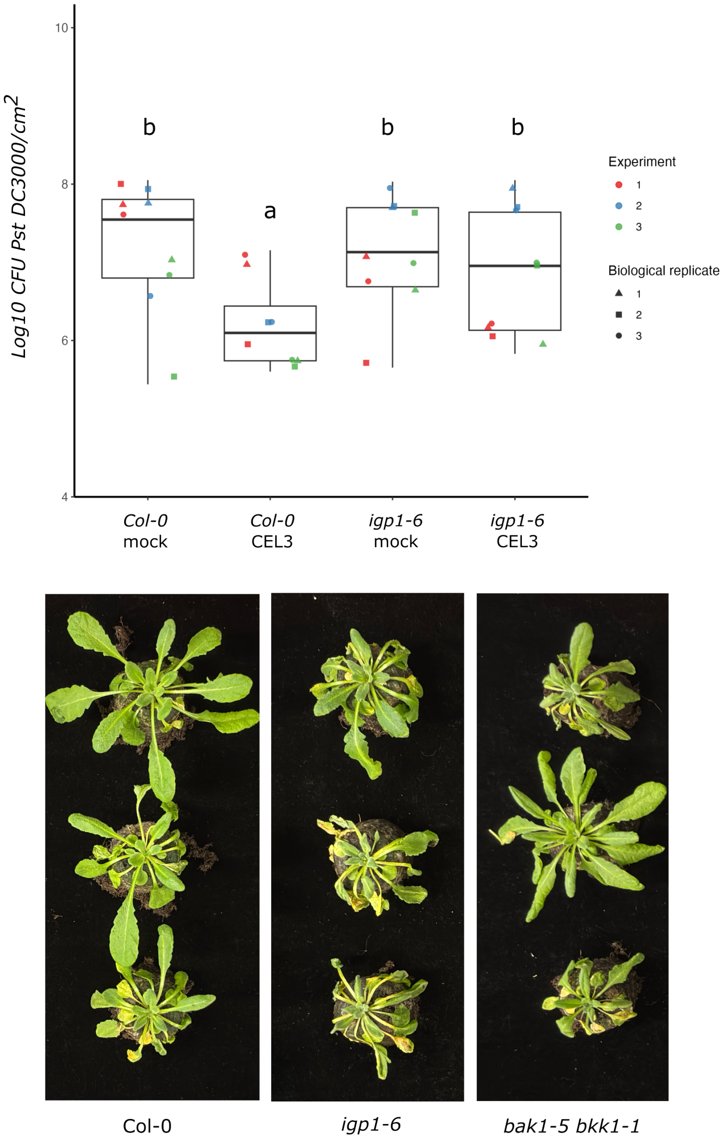


**Fig. S27 IGP1 enhances Arabidopsis immunity against pathogen infection**. (A) Pre-treatment with cellotriose (CEL3) reduces *Pseudomonas syringae* DC3000 growth in an IGP1-dependent manner at 72 hpi; buffer served as mock control. Statistical significance was assessed using a Kruskal–Wallis test followed by Dunn's post hoc test with Bonferroni correction (p < 0.001). Experiments were repeated independently three times. (B) Representative images of disease symptoms at 9 days post–soil drench inoculation in Col-0, *igp1-6*, and the susceptible control mutant *bak1-5/bkk1-1* (pictures from data represented Fig. 5 I-J).

**Table S1. Crystallographic data reduction and refinement statistics.**

| PDB ID | Apo AtIGP1 9HHU | AtIGP1/CEL3 complex 9HHX |
|---|---|---|
| **Data-reduction** | | |
| Space group | *P* $6_1$ | *P* $6_1$ |
| Wavelength (Å) | *1.000010* | *1.033286* |
| Cell dimensions | | |
| *a, b, c* (Å) | 83.78, 83.78, 198.94 | 84.37, 84.37, 198.32 |
| $\alpha, \beta, \gamma$ (°) | 99.00, 90.00, 120.00 | 90.00, 90.00, 120.00 |
| Resolution (Å) | 48.95 – 2.16 (2.23 -2.16) | 49.58 - 2.62 (2.74 - 2.62) |
| $R_{meas}$* | 0.137 (2.472) | 0.475 (2.436) |
| CC(1/2) *(%)** | 99.80 (31.60) | 97.30 (31.40) |
| $I/\sigma I$* | 10.0 (1.0) | 5.9 (1.0) |
| Completeness (%)* | 100.0 (100.0) | 100.0 (100.0) |
| Redundancy* | 7.8 (6.5) | 10.5 (8.7) |
| Wilson B-factor | 45.37 | 35.30 |
| **Refinement** | | |
| Resolution (Å) | 48.99 – 2.16 (2.22 -2.16) | 49.07 – 2.62 (2.69 - 2.62) |
| No. reflections | 42194 | 23962 |
| $R_{work/}$ $R_{free}$$ | 0.169 / 0.210 | 0.182 / 0.256 |
| No. atoms | | |
| Protein | 4456 | 4487 |
| Glycan | 257 | 243 |
| Ligand (CT3) | 34 | - |
| R.m.s deviations$ | | |
| Bond lengths (Å) | 0.015 | 0.012 |
| Bond angles (º) | 2.289 | 2.335 |
| Molprobity results | | |
| Ramachandran outliers (%)# | 0.17 | 0.00 |
| Ramachandran favored (%)# | 95.77 | 95.43 |
| Molprobity score# | 2.01 | 2.21 |

Highest resolution shell is shown in parenthesis.
*As defined in Xia2 /Dials (*101*)
$As defined in Refmac5 (*102*)
#As defined in Molprobity (*103*)

**Table S2. Primers used in the study.**

| Name | Primer sequence |
|---|---|
| IGP1$_{D74A}$ Fw 5’ | GACAGTGTCAGCATCGCCAATTTAGCTTTC |
| IGP1$_{D74A}$ Rv 5’ | GAAAGCTAAATTGGCGATGCTGACACTGTC |
| IGP1$_{N126A}$ Fw 5’ | CTCTAATCTGAACCTAGCTCAGAATTTCTTGACTG |
| IGP1$_{N126A}$ Rv 5’ | CAGTCAAGAAATTCTGAGCTAGGTTCAGATTAGAG |
| IGP1$_{D714A}$ Fw 5’ | GATCATTAGCGATTGCTATGAATAATTTC |
| IGP1$_{D714A}$ Rv 5’ | GAAATTATTCATAGCAATCGCTAATGATC |
| IGP1$_{D829N}$ Fw 5’ | GGTCCCGAAACTCTCAAATTTTGGGTTGGCCAAAC |
| IGP1$_{D829N}$ Rv 5’ | GTTTGGCCAACCCAAAATTTGAGAGTTTCGGGACC |
| IGP1$_{Y196A}$ Fw 5’ | GTG AAA ATG GCC ATT GGA AGT TCA GGA CTT |
| IGP1$_{Y196A}$ Rv 5’ | ACT TCC AAT GGC CAT TTT CAC TAG CCT CGT |
| IGP1$_{W146A}$ Fw 5’ | CGAATGCAGGCGATGACTTTTGGGGCCAAT |
| IGP1$_{W146A}$ Rv 5’ | CCCAAAAGTCATCGCCTGCATTCGAGTGAGATT |
| IGP1 prom Fw 5’ | AACAGGTCTCAACCTAGACGTTGTGCTACTTGC |
| IGP1 prom Rv 5’ | AACAGGTCTCTTGTTCGTCGACGACCAAAGATGTG |
| SAND qPCR Fw 5’ | AACTCTATGCAGCATTTGATCCACT |
| SAND qPCR Rv 5’ | TGATTGCATATCTTTATCGCCATC |
| IGP1 qPCR Fw 5’ | ACC TTC TCG AAT GGG CAT GG |
| IGP1 qPCR Rv 5’ | CTC CAC ATC ACC GGT CAA CA |
| IGP1-HA qPCR Fw 5’ | GAG CTG GAT ATT GCC TGA AAC |
| IGP1-HA qPCR Rv 5’ | GTA TGG ATA AGC ACT CGC ACC- |
| IGP4 qPCR Fw 5’ | GCTTTCAGGAGATGTTGAGGTCAG |
| IGP4 qPCR Rv 5’ | GTGAAGGATTCAGAAGCCTGTG |
| Chitinase qPCR Fw 5’ | CCAATCGTTCGACGCCTATAA |
| Chitinase qPCR Rv 5’ | GTGCTCGGTGAGCAGTATTT |
| SCOOP28 qPCR Fw 5’ | TGAAGGGAAGCTTGTGGTTGA |
| SCOOP28 qPCR Rv 5’ | TGTTCTTATTTCCAAAAGTTTGCGG |
| PEP2 qPCR Fw 5’ | GGACAACAAGGCCAAATCAAAG |
| PEP2 qPCR Rv’ | CCGCGTTGGGTACACTATTA |
| igp1-2F | CTTGGTTGGTCACAGCGTTT |
| igp1-2R | TATCACGCTGCTCTTGGTGT |
| igp1-3 F | GAACCGAACCACTGCCACTA |
| igp1-3 R | GAGTGAGTGTGTTCCAAGTGC |
| igp1-4F | CCGGTTCTCTACCACCTGAG |
| igp1-4 R | TCTCCTGCGGTAAAGCATGT |
| igp1-5F | TCTTCCTTCTTGGGTTCGCC |
| igp1-5R | CCTACATTGCTGACTGCCCA |
